# A comparative study of stress responses elicited by misfolded proteins targeted by bipartite or matrix-targeting signal sequences to yeast mitochondria

**DOI:** 10.1101/2020.08.16.252734

**Authors:** Kannan Boosi Narayana Rao, Pratima Pandey, Rajasri Sarkar, Asmita Ghosh, Shemin Mansuri, Mudassar Ali, Priyanka Majumder, K. Ranjith Kumar, Arjun Ray, Swasti Raychaudhuri, Koyeli Mapa

## Abstract

The double-membrane-bound architecture of mitochondria, essential for ATP production, sub-divides the organelle into inter-membrane space (IMS) and matrix. IMS and matrix possess contrasting oxido-reductive environments and distinct protein quality control (PQC) machineries resulting different protein folding environments. To understand the nature of stress response elicited by equivalent proteotoxic stress to sub-mitochondrial compartments, we fused well-described bipartite or matrix-targeting signal sequences to misfolding and aggregation-prone stressor proteins to target and impart stress to yeast mitochondrial IMS or matrix. We show, mitochondrial proteotoxicity leads to growth arrest of yeast cells of varying degrees depending on nature of stressor proteins and the intra-mitochondrial location of stress. Next, using transcriptomics and proteomics, we report a comprehensive stress response elicited by two types of targeting signal-fused stressor proteins. Among global responses by mitochondria-targeted stressors by both types of signal sequences, an adaptive response of abrogated mitochondrial respiration and concomitant upregulation of glycolysis is uncovered. Beyond shared stress responses, specific signatures due to stress within mitochondrial sub-compartments are also revealed. We report that bipartite signal sequence-fused stressor proteins eliciting stress to IMS, leads to specific upregulation of IMS-chaperones and TOM complex components. In contrast, matrix-targeted stressors lead to specific upregulation of matrix-chaperones and cytosolic PQC components. Finally, by systematic genetic interaction using deletion strains of differentially upregulated genes, we found prominent modulatory role of TOM complex components during IMS-stress response. In contrast, *VMS1* markedly modulates the stress response originated from matrix.

## Introduction

Mitochondria, the cellular powerhouse, imports almost 99% of its constituent proteins as pre-proteins from cytosol. These pre-proteins remain unfolded till they reach their site of function inside the mitochondria where precursor proteins are eventually folded and often assembled (into complexes) to their functional forms by the mitochondrial matrix (MM) and the intermembrane space (IMS)-resident chaperone machineries [1–3]. Thus, mitochondrial IMS and matrix are continuously exposed to non-native unfolded proteins which when combined with other sources of proteotoxic stress, threaten the health of the organelle. Apart from presence of considerable amount of reactive oxygen species (ROS) generated as a by-product of oxidative phosphorylation by the organelle, recent literature shows varied sources of proteotoxic threat to mitochondria like aggregation prone proteins from cytosol which requires clearance by MAGIC pathway [4], mistargeted proteins from neighbouring organelles like endoplasmic reticulum (ER) [5] or higher intra-mitochondrial temperature [6].

To combat such imminent proteotoxic stress, designated chaperones and proteases maintain the folding environment (together known as mitochondrial Protein Quality Control (PQC) machinery) in both the sub-mitochondrial compartments, although the nature of folding machineries is markedly different between matrix and IMS [1–3]. When the amount of unfolded, misfolded or unassembled proteins overwhelms the folding capacity of either of these compartments, the perturbed mitochondrial homeostasis elicit a stress response known as mitochondrial <u>U</u>nfolded <u>P</u>rotein <u>R</u>esponse (mitoUPR) [7, 8].

While mitoUPR has been extensively studied, a vast majority of these studies relied on external stressors that perturb the mitochondrial protein homeostasis indirectly [9–11]. Notably, the evidence of mitoUPR originating exclusively from accumulation of misfolded or aggregated proteins in mitochondria is rare in literature [12, 13]. The concept of compartmentalized proteostasis within mitochondria is even fewer, given the lack of selectivity of small-molecule stressors that are routinely used to impart proteotoxic stress. If only one sub-compartment of mitochondria is affected by proteotoxic stress, will the cellular response be distinct? Can cells distinguish intra-mitochondrial location of the misfolded or aggregated proteins and respond accordingly?

To answers these questions, we fused three different stressor proteins possessing varied misfolding or aggregation-propensities to previously described bipartite signal sequence or its truncated form for targeting to the mitochondrial IMS or matrix, respectively. We show that the stress elicited by these differentially-targeted stressors are at least partially different in terms of the cellular capacity to handle the mitochondrial proteotoxic stress. Although both scenarios culminate into a global mitochondrial phenotype (severe fragmentation), the stress response signature is partly distinct and unique when compared between IMS and matrix stress. A global cellular adaptive response to mitochondrial proteotoxic stress is abrogated mitochondrial respiration which is well corroborated with prominent downregulation of components of Electron Transport Chain (ETC) found in transcriptomics and proteomics data. We show that yeast efficiently cope with mitochondrial stress by suspending mitochondrial respiration and survive by a strategy of altered metabolism. Apart from global adaptive responses due to mitochondrial proteotoxic stress, some specific signatures for IMS and matrix stress are also captured. Transcriptomics data revealed upregulation of IMS-specific protein quality control (PQC) components like *ERV1* and small Tim proteins by bipartite signal sequence-targeted stressors. In contrast, matrix-targeting-signal-fused stressors lead to differential upregulation of matrix-resident PQC components like Hsp60, Ssc1, Tim44 along with components of cytosolic PQC. We have termed the specific responses as IMS-UPR and mito-matrix-UPR respectively. Finally, by systematic genetic interaction experiments, we show that subunits of TOM complex and Msp1 (an extractase of outer mitochondrial membrane for mistargeted tail-anchored proteins [14–16] and a component of mitoCPR pathway[17]) act as specific modulators of IMS-UPR. In contrast, Vms1, a component of Ribosome Quality Control (RQC) [18–21]and Mitochondria-Associated Degradation (MAD) [22, 23] process, exhibits prominent role during mito-matrix UPR. The detailed molecular mechanism of the modulatory role of TOM complex components and Vms1 in IMS and matrix stress respectively, remains to be explored in future.

## Results

### Mitochondrial IMS and matrix are susceptible to proteotoxic stress

We generated mitochondrial proteotoxic stress model in yeast, *Saccharomyces cerevisiae* by expressing model misfolded and aggregation-prone proteins within mitochondria. We took three different exogenous model proteins of varied misfolding properties as stressors (Figure 1A). In parallel, we expressed well folded wild type version of these proteins as controls to dissect the cellular response elicited exclusively due to misfolding or aggregation and not due to overburdening of mitochondrial protein quality control machineries with foreign proteins. Mitochondrial Inter-membrane Space (IMS) and Mitochondrial Matrix (MM)-specific targeting of these model proteins was done by fusing the genes encoding the stressor proteins to well characterized bipartite signal sequence of yeast Cytochrome b2 (Cyb2SS) or its truncated version (lacking the 19 amino acid membrane sorting signal, also known as Cyb2ΔSS), respectively [24, 25] (Figure 1A, lower panel). To capture mitochondria-specific response pathways, simultaneously, we generated yeast strains expressing these stressor proteins in other subcellular compartments including ER, nucleus (N), cytosol (Cyto), apart from mitochondrial matrix or IMS by fusing the proteins with well described targeting signals (as summarized in Figure 1B). All misfolded proteins or the wild type counterparts were expressed from inducible Gal1 promoter with a Cyc terminator for transcription termination (Figure 1B, upper schematic), integrated in the URA3 locus (Figure S1A). Expression of stressor proteins as well control proteins were only observed post-induction, as assessed by western blots (Figure S1B). Proteins targeted to mitochondrial IMS or matrix were checked for cellular localization using some of the stressor (DMMBP and PMD) and control (MBP) proteins (Figure S1 C-F). Except for a minor fraction of Cyb2ΔSS-fused PMD protein (MM-PMD) (Figure S1E) which is found in cytosolic fraction, other proteins are exclusively localized in the mitochondrial fraction (Figure S1 C-F).

**Figure 1:**
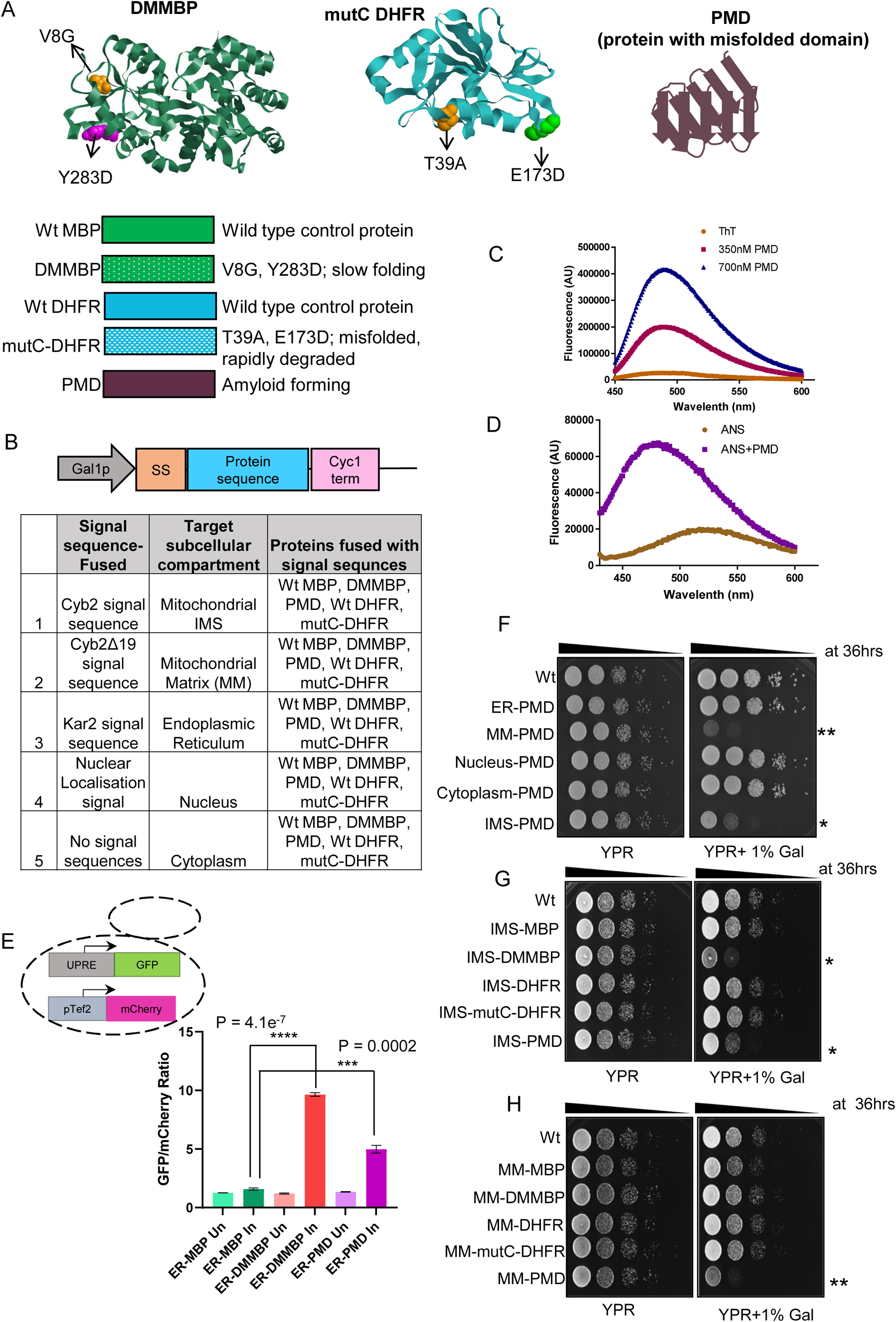
Sub-cellular compartment-specific proteotoxic stress model generation in yeast and its phenotypic characterization. **A**. Upper panel: Left, middle and right panels are depicting ribbon structure of *E. coli* MBP (PDB ID 1OMP, mutations of DMMBP, V8G and Y283D are shown as space fill), mouse DHFR (PDB ID 3D80, mutations of mutC-DHFR, T39A, E173D, are shown as space fill), and schematic picture of amyloid forming PMD protein respectively. Lower panel: Schematic representation of all proteins with their properties used for sub-compartment specific targeted expression in yeast, *Saccharomyces cerevisiae*. **B.** The top schematic represents DNA cassettes used for integration in Wt yeast strains for inducible expression of stressor proteins specifically targeted to different sub-cellular compartments. “SS” indicates <u>S</u>ignal <u>S</u>equences for targeting to different subcellular compartments like endoplasmic reticulum (ER), mitochondrial IMS, mitochondrial matrix and nucleus. Gal1p and Cyc term indicates galactose-inducible Gal1 promoter and Cyc terminator, respectively. Lower table summarizes signal sequences fused to stressor (or control) proteins for targeted expression within different sub-cellular compartments. **C**. Fluorescence emission spectra of free Thioflavin T (ThT) and ThT bound to purified PMD protein in two different concentrations are shown. **D**. Fluorescence emission spectra of 1,8 bis-ANS in buffer and bound to purified PMD protein are shown. **E**. Yeast strain expressing the control protein MBP or stressor proteins DMMBP or PMD specifically targeted to endoplasmic reticulum (ER) were grown till mid log phase. Subsequently, stressor (or control) proteins were induced with 1% galactose for 6 hours. After induction, GFP and mCherry fluorescence were measured by flow cytometry. The ratio of mean GFP and mCherry fluorescence was plotted as bar plots for all strains. A schematic picture of the UPRE-GFP reporter with mCherry under constitutive Tef2 promoter as control of cellular transcription/translation is shown as inset. Error bars represent standard deviation between repeats (n=3). P-value was calculated by unpaired Student’s T test (2­tailed). **F**. Growth phenotype assay using drop-dilution of different yeast strains expressing the amyloid forming PMD protein in different subcellular compartments including mitochondrial IMS and matrix. Yeast strains grown till mid log phase (OD_600_ of 0.5) were serially diluted (1:10) and spotted on without inducer (YPR, <u>Y</u>east <u>E</u>xtract-<u>P</u>eptone-<u>R</u>affinose) and with inducer [YPR+1% gal (<u>Y</u>east-extract-<u>P</u>eptone-<u>R</u>affinose +1% galactose as inducer)] plates. In the YPR+1% gal plates, slow growth phenotype of IMS-PMD and MM-PMD strains are visible compared to Wt strain and other spotted strains while on the control plate without inducer (YPR) all strains grow similarly. MM­PMD phenotype is more severe as indicated by slow growth from the first spots in comparison to IMS-PMD where the phenotype is evident from second spot onwards. These phenotypes have been denoted as “**” and “*” for MM-PMD and IMS-PMD strains respectively to emphasize the more severe phenotype of MM-PMD compared to IMS-PMD in respect to Wt strain. The time mentioned in hours at the upper right corner of the panel indicates the time of incubation of the spotted plates at 30°C before capturing the image. **G-H**. Drop-dilution assay as described in panel F is shown for strains containing mitochondrial IMS targeted DMMBP (with wt MBP as control) and mutC-DHFR (with Wt DHFR as control) (panel **G**). Growth phenotype assay of the strains containing mitochondrial matrix targeted stressor and control proteins similar to panel G (panel **H**). “*” and “**” indicates spots with visible growth phenotype in case of IMS-PMD/IMS-DMMBP and MM-PMD strains respectively. The time mentioned in hours at the upper right corner of the panel indicates the time of incubation of the plates at 30°C.

We preferred exogenous model proteins over aggregation prone mutants of endogenous proteins to capture the stress response arising particularly due to misfolding stress and not due to secondary effects of aggregation prone endogenous proteins that might 1) recruit endogenous interaction partners in the aggregated species leading to loss of function and toxicity, 2) co-aggregation of the wild type copy of the protein, where keeping the endogenous copy of the protein becomes essential, 3) exert dominant negative function due to futile interaction with the interaction partners or 4) lead to rewiring of transcription to upregulate a compensatory or parallel pathway due to dominant negative function of the overexpressed mutant protein.

We chose slow folding mutant version of *E. coli* Maltose binding protein (MBP) (Figure 1A, left panel), also known as double-mutant MBP (DMMBP) as one of the stressors [26, 27]. Due to slow folding rate of DMMBP, at any given time point the concentration of the non-native conformation of this protein is higher than that of a fast-folding protein and is expected to induce proteotoxic stress [28]. Indeed, when DMMBP is expressed in the lumen of endoplasmic reticulum (ER) of yeast cells, it elicits ER-UPR (ER-Unfolded Protein Response) but wild type MBP does not induce ER-UPR (Figure 1E) [28]. A previously described mutant form of mouse dihydro-folate reductase (mutC-DHFR) which is a misfolded protein with a high turnover rate (Figure 1A, middle panel) as second stressor protein [29]. The third stressor protein is an amyloid forming protein and hereafter named as protein with misfolded domain (PMD) (Figure 1A, right panel). Previous studies have shown that artificial beta sheet containing model proteins form toxic aggregates and these aggregates sequester crucial components of cellular protein quality control machinery resulting in toxicity [30]. PMD being much larger in size (~30kDa) than the model beta sheet forming proteins, is comparable to many endogenous yeast proteins in size and was expected to be toxic. PMD is predicted to contain secondary structural elements of approximately 33.3% of α-helices, 39.3% of ß-strands and 27.4% of random coils (Figure S2A and S2B). Recombinant PMD protein was cloned and purified from *E. coli* (Figure S2C) and characterised for misfolding and amyloid-forming properties indicating strong amyloid formation (Figure 1C). Additionally, PMD showed increased fluorescence emission and blue shift of the fluorescent probe 1,8, ANS (Figure 1D) illustrating surface-exposed hydrophobicity, a hallmark of misfolding. When expressed in ER lumen of yeast, PMD elicited ER-UPR like DMMBP (Figure 1E) and mutC-DHFR (Figure S2D) indicating that PMD was recognised as a misfolded protein in the ER and was able to mount ER-UPR (Figure 1E).

Importantly, among the different subcellular compartments tested, mitochondrial matrix (MM) and IMS exhibited substantial growth defects upon expressing the amyloid forming protein, PMD. The observed growth defect is more pronounced in matrix than in IMS (Figure 1F). Expression of the slow folding mutant, DMMBP, imparted growth phenotype only to IMS. Expression of rapidly degrading misfolded protein mutC-DHFR did not impart any growth defect to either of the mitochondrial sub-compartments (Figure 1G and 1H). These results indicate that misfolded proteins trigger heterogeneous cellular responses depending on both their intrinsic misfolding property and their sub-compartmental location. All three stressor proteins did not exhibit any growth defect upon expression at other cellular compartments denoting mitochondria may be more sensitive to proteotoxic insults (Figure 1F, S1G and S1H).

We confirmed the targeting of fused proteins by Cyb2SS or Cyb2ΔSS signal sequences to mitochondrial IMS and matrix respectively, using wild type MBP protein by mitoplasting (removal of the outer mitochondrial membrane by osmotic shock) (Figure 2A). Intact mitochondria isolated from MM-MBP strain show two forms of the MBP protein, the p (precursor form) and the m (mature) forms (Figure 2A). MBP fused to Cyb2SS and targeted to IMS shows p, m and an additional i (intermediated) form (Figure 2Bi). Protease treatment by proteinase K (PK) of intact mitochondria keeps all the forms of MBP protein protected in both matrix and IMS of mitochondria, as expected. Upon solubilization of membranes with detergent (Tx-100) with PK treatment, a smaller band indicating a protease-resistant smaller form of the protein denoted as m* form is detected (Figure 2Ai and 2Bi). After mitoplasting, majority of the Cyb2SS-fused IMS-targeted MBP protein is released from mitochondria due to opening of the outer mitochondrial membrane and major fraction of the protein is found in the supernatant fraction (Figure 2Bii) after re-isolation of the mitochondrial pellet. Correspondingly, only a small amount of three forms of the protein is retrieved in mitochondrial pellet indicating that the protein is localized in the IMS (Figure 2Bi). The equivalent amount of protease digested m* form found in the supernatant after protease digestion of the mitoplasts and detergent-solubilized mitoplasts further confirms that the protein is exposed to protease digestion just after mitoplasting. For matrix targeted protein, upon mitoplasting, majority of the protein is retrieved in mitochondrial pellet after re-isolation (Figure 2A). In contrast to Cyb2SS-fused IMS targeted MBP, MM-MBP mitoplasts show the m* form only upon protease digestion following detergent solubilization. A minor amount of different forms of the MBP protein is found in the supernatant in isotonic buffer or in hypotonic buffer in case of matrix targeted MBP protein where it should not be released from intact membranes which may happen due to problems of integrity of the mitochondria membranes due to overexpressed proteins in it. Similar pattern of localization within mitochondria was observed with DMMBP (data not shown). For PMD, localization experiment was extremely difficult as the protein formed large insoluble aggregates.

**Figure 2:**
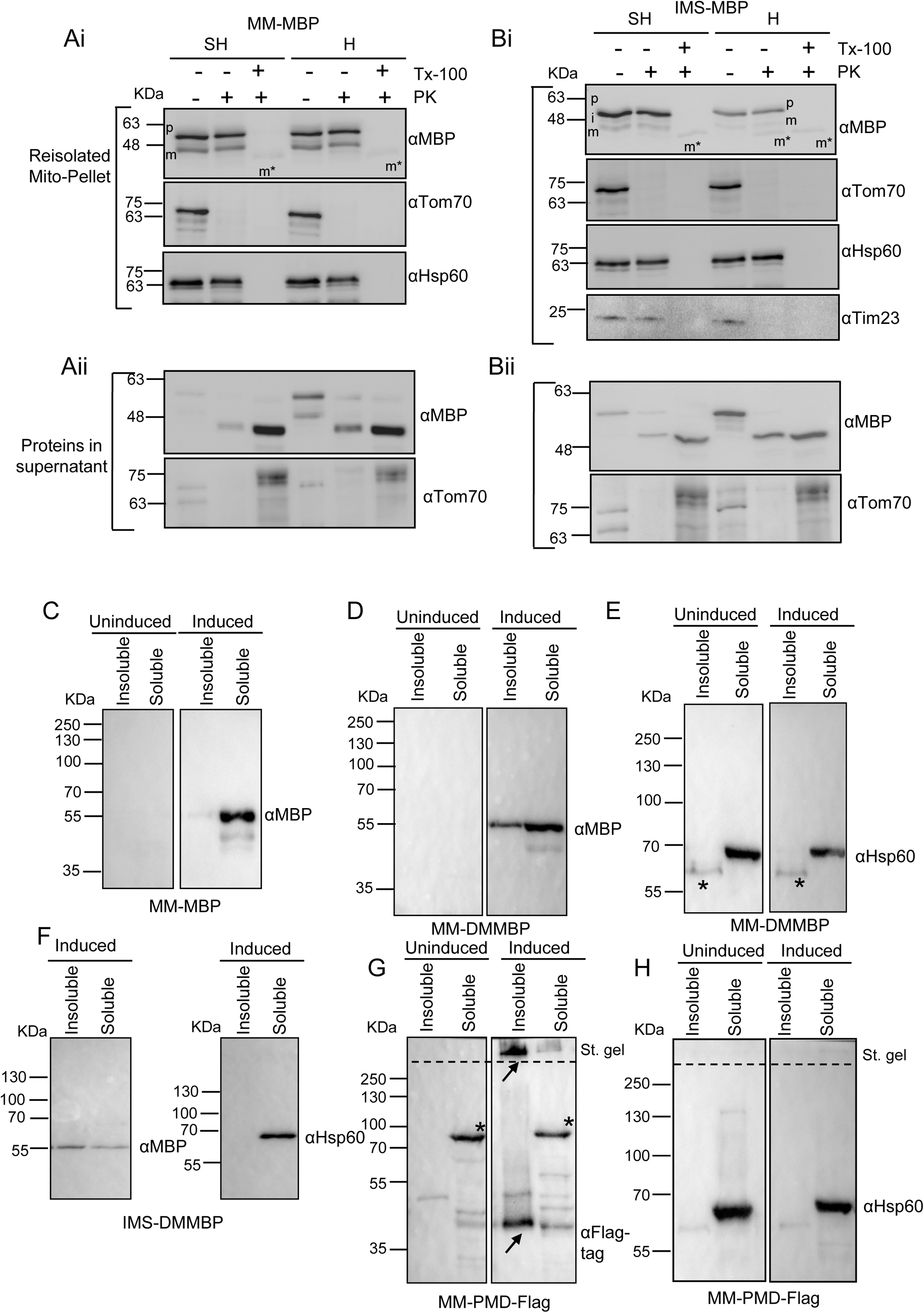
Localization of Cyb2SS and Cyb2ΔSS-fused MBP protein within mitochondria and characterization of intra-mitochondrial misfolding of stressor proteins. **A.** Western blot to confirm the localization of Cyb2ΔSS-fused MBP protein which targets the protein to mitochondrial matrix was performed with anti-MBP antibody using reisolated mitochondrial pellets following treatment of isolated mitochondria from MM-MBP strain (mitochondria isolated after 12 hours of induction with 1% galactose) with protease, Proteinase K (PK) in isotonic buffer (SH buffer; sorbitol-HEPES buffer, details in methods section) or hypotonic (H buffer; HEPES buffer) buffer to prepare mitoplasts. For complete solubilization of mitochondrial membranes, 0.1% Tx-100 was used. Two forms of the MBP protein shown as p (precursor form) and m (mature) forms are observed. In Tx-100 with PK treated samples, a smaller form of the protein indicating a protease-digested, smaller form of the protein denoted as m* form is found. As markers for outer membrane Tom70 antisera was used and as markers for matrix resident proteins, Hsp60 antisera was used. **B.** Similar to panel A, localization of Cyb2SS-fused MBP which targets the protein to mitochondrial IMS, was performed as following. Reisolated mitochondrial pellets after treatment of isolated mitochondria from IMS-MBP strain (mitochondria isolated after 12 hours of induction with 1% galactose) with PK in SH buffer or H buffer were used as described in panel A. For membrane solubilization 0.1% Tx-100 was used. Like matrix targeted MBP, two forms of the MBP in the IMS, p (precursor form) and the m (mature) forms along with an additional i (intermediated) form are observed. After mitoplasting in H buffer with PK treatment, we observed the p, m and the protease-resistant smaller m* form in the IMS targeted protein in the re-isolated mitochondrial pellet (Panel Bi) and majority of the protein is found in supernatant only by mitoplasting with H buffer (Panel Bii). Mitoplasting with PK digestion leads to digestion to m* form which is also found majorly in supernatant after re-isolation of mitochondrial pellet (Panel Bii). In Tx-100 with PK treated samples only the protease digested m*form is found which is retrieved in the supernatant fraction due to solubilization of mitochondrial membranes (Panel Bii). As marker for outer membrane Tom70 antisera, Tim23 antibody as marker for IMS exposed protein and as markers for matrix resident proteins, Hsp60 antisera was used. **C-H.** To show the aggregation propensity of stressor proteins used, mitochondria of IMS-DMMBP (panel F) or MM-DMMBP (panel D) or MM-PMD-FLAG (panel G) were isolated after expressing the proteins for 12 hours with 1% galactose in YPR. As control, well folded wild type MBP protein (panel C) was taken from MM-MBP strain. Isolated mitochondria were subjected to solubilization with 0.1% Tx-100 in SH buffer and was separated into detergent soluble and insoluble fractions by high-speed centrifugation. The pellet (insoluble fraction) and supernatant (soluble fraction) were loaded, and western blot was performed with anti-MBP or anti-FLAG antibody. As a marker for soluble endogenous protein, Hsp60 was taken (panels E and H). As part of MM-PMD-flag was forming big aggregates and were stuck in the stacking gel, western blot was performed with the stacking gel for MM-PMD-FLAG mitochondria (panel G, also for control panel in H). “*” indicates non-specific bands in western blots in panel G and H. Arrowheads in panel G indicates PMD-FLAG protein in resolving and stacking gels.

To assess the misfolding and aggregation propensity of the stressor proteins within mitochondrial IMS and matrix, we solubilized the isolated mitochondria with Tx-100 and fractionated the mitochondrial proteins into detergent-soluble and insoluble parts. Indeed, the stressor proteins, DMMBP and PMD proteins were substantially found in the detergent-insoluble mitochondrial fractions (Figure 2D, F and G) although the control protein MBP (Figure 2C) or mitochondrial Hsp60 (used as marker protein) was found in the soluble fraction (Figure 2E, F-right panel and H). A large portion of the PMD protein remained stuck at the stacking gel indicating formation of large aggregates which is in corroboration of the propensity of this protein to form amyloid aggregates as shown before (Figure 2G).

To differentiate whether growth arrest or cell death leads to observed growth phenotype after expressing Cyb2SS or Cyb2ΔSS-targeted stressors to mitochondria, we performed a recovery experiment. Following induction of the stressor proteins for 16 hours, we washed the culture to remove the inducer (galactose) and spotted the cells on the glucose containing media (YPD) where the Gal1 promoter remains completely repressed. Cells were expected to recover at this point if their growth was arrested due to proteotoxic stress. All strains grew similarly in YPD indicating that the strains with visible phenotype in presence of the inducer (MM-PMD, IMS-PMD and IMS-DMMBP) were growth arrested (Figure S3A). In parallel, to detect any cell death due to mitochondrial proteotoxic stress, yeast cells were stained with propidium iodide (PI). Insignificant cell death was observed (less than 2%) in strains which showed prominent growth phenotypes in presence of inducers (IMS-DMMBP, IMS-PMD and MM-PMD), confirming that the observed growth phenotypes are due to growth arrest of yeast cells (Figure S2E).

Taken together, we show that mitochondrial sub-compartments, matrix and IMS are susceptible to proteotoxic stress although are partially dissimilar in stress handling capacity as evidenced by different ability to tolerate equivalent stress imparted by expression of slow-folding mutant protein, DMMBP. Interestingly, extreme proteotoxic stresses by misfolded and aggregated proteins in mitochondrial IMS and matrix lead to only growth arrest of yeast cells without considerable cell death.

### Proteotoxic stress in IMS or mitochondrial matrix leads to altered mitochondrial dynamics and functions

Proteotoxic stress in mitochondrial sub-compartments led to significant growth defects, hinting an alteration in mitochondrial function. As dysfunctional mitochondria are often associated with altered mitochondrial morphology, we checked for any change in mitochondrial morphology following misfolding stress. Imaging of mitochondrial network was done expressing mitochondria-targeted yeGFP (yeast enhanced Green Fluorescent Protein) in yeast cells. The uninduced, control cells showed tubular network of mitochondria with continuous fission and fusion indicating normal mitochondrial morphology (Figure 3A, minus Gal panels). In contrast, the induced strains with visible growth defect e. g. IMS-DMMBP, IMS-PMD (Figure 3A, middle and right panel) and MM-PMD (Figure S3C, lowest panel) showed severely fragmented mitochondria with prominent disruption of the tubular network. In some strains (e.g., IMS-PMD) disruption of tubular mitochondrial network was extremely severe and the fragmented mitochondria accumulated at one pole of the cells (Figure 3A, right panel). Similarly, upon expression PMD protein with a C-terminal GFP tag in either matrix or IMS, we observed severe disruption of mitochondrial network (Figure S3B). The cells with overexpressed WT proteins (MBP or DHFR) (data not shown) or mutC-DHFR (Figure S3C, upper and middle panel) did not show any alteration in mitochondrial network indicating mere overexpression of foreign proteins from the same inducible Gal1 promoter in mitochondrial IMS or matrix do not lead to altered mitochondrial morphology (Figure S3C). Notably, this altered mitochondrial dynamics upon proteotoxic stress was recapitulated in mammalian cells. By targeting PMD protein to mitochondrial matrix or IMS in HeLa cells [20, 31], we observed severe fragmentation of mitochondrial tubular network in both matrix and IMS stress, more prominently in the matrix stress (Figure 3B). Despite severe fragmentation of mitochondrial network, cell death was negligible in mammalian cells too (Figure S3D). Apart from fragmentation, IMS-PMD and MM-PMD cells were substantially less stained with MitoTracker green (a well-established fluorescent probe which specifically accumulates within functional mitochondria) following induction of proteotoxic stress compared to uninduced cells (Figure S3E). Importantly, MM-PMD cells had far more severe defect in uptake of Mitotracker green probe, after expression of the stressor protein (Figure S3E).

**Figure 3:**
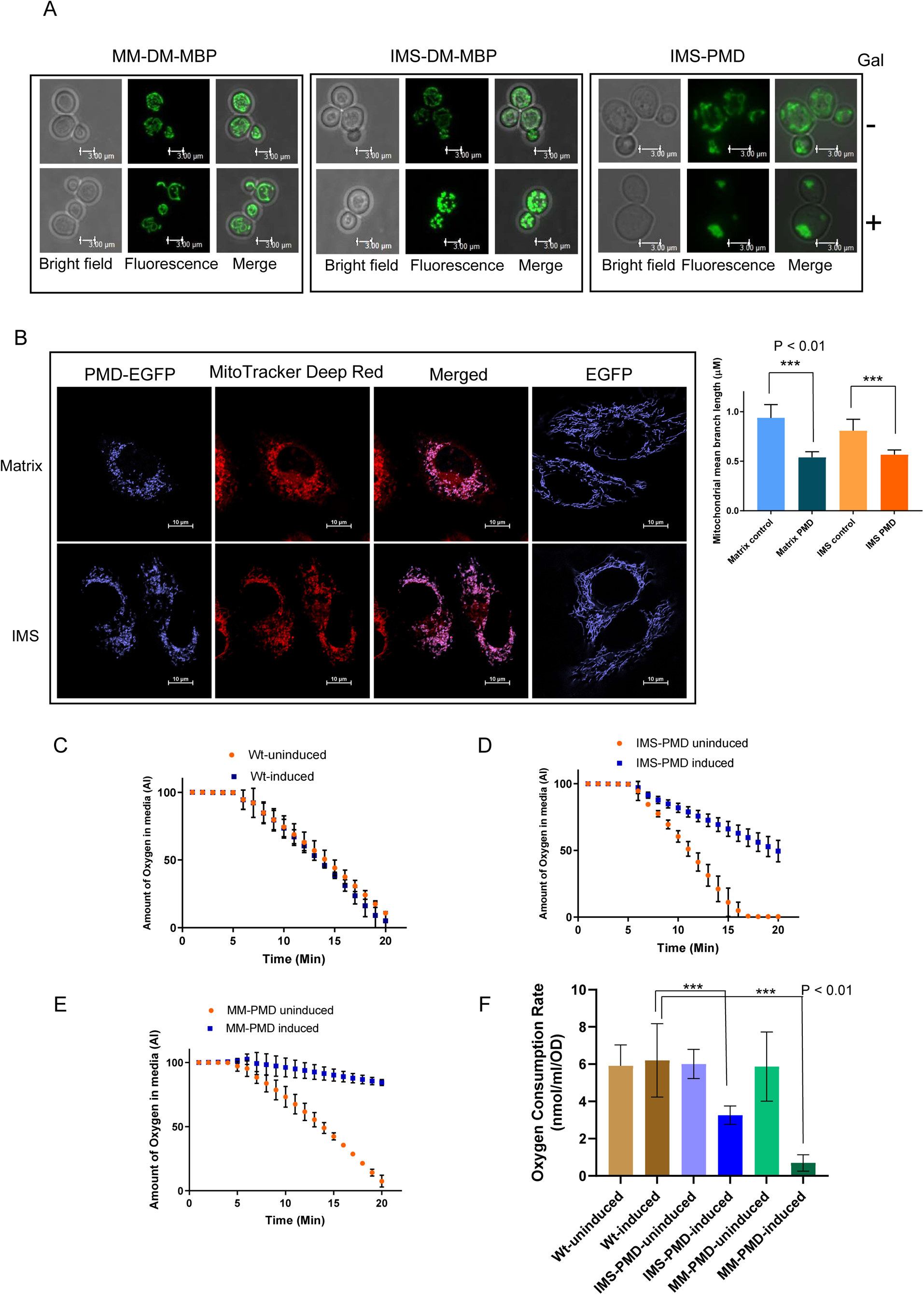
Proteotoxic stress in mitochondrial sub-compartments, IMS and matrix leads to alteration in mitochondrial forms and functions. **A.** Fluorescence confocal microscopy of yeast cells showing green, fluorescent mitochondria expressing mitochondria targeted yeGFP in the MM-DMMBP (Left panel), IMS-DMMBP (Middle panel) and IMS-PMD strains (Right panel) in absence and presence of galactose induction of stressor proteins in respective mitochondrial sub-compartments. **B.** Left panel: Fluorescence confocal microscopy of HeLa cells showing fluorescent mitochondria constitutively expressing matrix and IMS targeted PMD protein tagged with C-terminal EGFP. To show the mitochondria specific localization of PMD­GFP, co-localization with MitoTracker Deep Red dye have been shown. Right panel: Bar plot showing the mean branch length of mitochondria in control cells (cells expressing EGFP in matrix or IMS) and cells expressing PMD proteins specifically in matrix or IMS. Error bars represent standard deviation among different mitochondrial segments (n=15). Statistical analysis was done by Student’s t-test. **C-E.** Representative kinetic traces of Oxygen consumption rate (OCR) of wild type (Wt) (panel C), IMS-PMD (panel D) and MM-PMD (panel E) yeast strains are shown in absence and presence of galactose induction. F. Oxygen consumption rate calculated as nmol of O_2_ consumed per ml of media by per OD of yeast cells is plotted as bar plots. Error bars represent standard deviation between repeats (n=3). P-value was calculated by unpaired Student’s T test (2­tailed).

To ascertain the respiratory status of the mitochondria, we measured the oxygen consumption rate (OCR) of yeast cells expressing the stressor proteins in IMS or matrix. MM-PMD and IMS-PMD strains showed significant reduction in OCR compared to wild type cells following proteotoxic stress; the extent of reduction of OCR is more prominently observed in case of matrix stress (MM-PMD) (Figure 3C, D, E and F).

In summary, we show that proteotoxic stress due to accumulation of misfolded or aggregated proteins in mitochondrial sub-compartments lead to severe mitochondrial fragmentation and significantly reduced mitochondrial respiration. Interestingly, such changes in the organellar morphology and compromised function are well tolerated by yeast as well as human cells as we observed negligible cell death in both the systems. This result indicates presence of robust stress response and quality control mechanisms to cope with mitochondrial proteotoxic stress in both yeast and human cells.

### Identical misfolded proteins elicit distinct stress response when expressed in mitochondrial IMS or matrix

As initial growth assays post-induction of proteotoxic stress indicated that mitochondria are most vulnerable among the tested subcellular compartments and sub-mitochondrial compartments (IMS or matrix) (Figure 1F and S1G-H) are at least partially dissimilar in stress handling capacity, it was interesting to check the cellular responses to equivalent proteotoxic insults specifically imparted in these compartments. we performed transcriptomics of the yeast strains expressing different stressor proteins targeted to different sub-cellular compartments (mito-IMS, mito-MM, ER, cytosol, nucleus) (details of strains are provided in Supplemental Table S2). To identify the specific set of genes that are differentially expressed during misfolding stress in a particular subcellular compartment, we calculated the Z-score of expression for each gene across all the yeast strains (table S2). The genes above or below Z-score 2 at a particular condition were considered as significantly upregulated or downregulated, respectively. Using Z-score analysis, we found 394, 210 and 285 genes to be upregulated upon expression of Cyb2SS-fused PMD, DMMBP and mutC-DHFR, respectively in the IMS. On targeting these proteins to mitochondrial matrix, 227, 222 and 25 genes were upregulated with expression of PMD, DMMBP and mutC-DHFR, respectively. 90 genes were downregulated upon expression of PMD protein in IMS while 15 genes were downregulated when expressed in matrix. Significantly down-regulated genes were lower than the number of upregulated genes.

As PMD protein showed prominent toxic effects in both matrix and IMS, we performed GO enrichment analysis of differentially overexpressed genes of mitochondrial IMS or matrix stress due to expression of PMD protein using Holm-Bonferroni test for multiple comparison (p-value <0.05) by Yeastmine webserver (https://yeastmine.yeastgenome.org/yeastmine). Various pathways related to mitochondrial biogenesis and function (mitochondrial translation, mitochondrial translocation, cytochrome complex assembly, mitochondrial gene expression) were highly enriched during stress by PMD protein in IMS (Figure 4A). In contrast, general protein folding, and refolding pathways were upregulated in matrix stress with PMD protein (Figure 4B). In corroboration to GO enrichment analysis, by performing gene interaction network (physical and genetic) analysis of the upregulated genes, we found prominently distinct network in IMS and matrix stress with the same stressor protein, PMD (Figure S4Aand S4B). Among the network of differentially upregulated genes during IMS stress, we found components of oxidative folding machinery like *ERV1*, small heat shock proteins of mitochondrial IMS like *TIM10, TIM13*, components of TOM complex like *TOM5, TOM6, TOM7* and components for assembly and chaperoning of Cytochrome C oxidase (*COX14, COX17*) machinery (Figure S4A). Apart from these prominent groups, many components of mitochondrial ribosome (components of both large and small subunits) were also found to be upregulated in IMS stress. On Comparing, network analysis of differentially upregulated genes during mitochondrial matrix stress showed distinct network of genes than IMS stress. In matrix stress, matrix resident chaperones and co-chaperones like *SSC1, HSP60, TIM44* were upregulated indicating the stress response is specific to matrix (Figure S4B). Interestingly, a substantial number of chaperones involved in cytosolic quality control like *SSA1, SSA2, SSA4, HSP82, HSC82, YDJ1, DJP1, SIS1, CCT2* were found to be upregulated specifically as a response to mitochondrial matrix stress (Figure S4B). Importantly, component of Ribosome Quality Control (RQC), *VMS1,* which is a peptidyl-tRNA endonuclease and protects mitochondria by antagonising the CAT-tailing activity of *RQC2* [20, 21] and also is an important component of Mitochondria Associated Degradation (MAD) pathway[22, 23], was present in the network of differentially upregulated genes of matrix specific proteotoxic stress response (Figure S4B).

**Figure 4:**
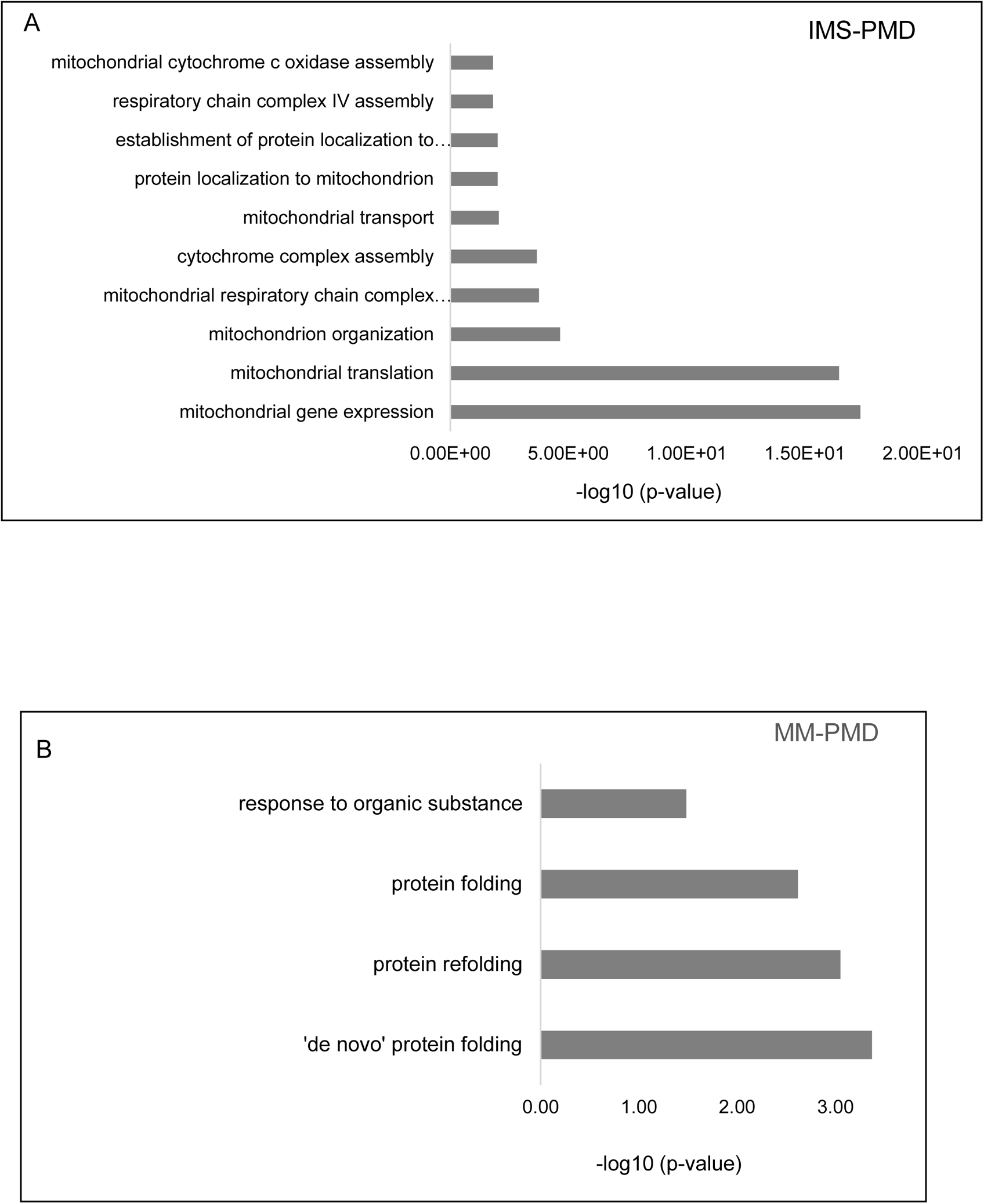
GO enrichment analysis of biological processes during mitochondrial proteotoxic stress. **A.** The differentially upregulated genes (with z-scores above 2) of PMD-mediated IMS stress (after 16 hours of galactose induction) from first set of transcriptomics data, were analysed for GO enrichment analysis with Holm-Bonferroni test for multiple comparison (p-value <0.05) using Yeastmine webserver. The enriched pathways with corresponding p-values were plotted as 2D-bar diagram. **B.** Similar to panel A, the differentially upregulated genes of PMD-induced matrix stress from the transcriptomics data were analysed and plotted for GO enrichment of biological processes using same criteria as described in panel A.

Taken together our data highlight that mitochondrial IMS or matrix has distinct pattern of stress response to proteotoxic stress and are catered by different adaptive mechanisms.

### Various components of Mitochondrial PQC and Translocases follow distinct kinetics during proteotoxic stress in IMS and matrix

Although the Z-score based analysis of gene expression changes across different yeast strains expressing misfolded proteins in different subcellular compartment gave us exclusive signature of transcriptional response of proteotoxic stress specific to IMS or matrix, it portrayed a snapshot of the late response (post 16 hours of induction of stressor protein expression). To capture the early response and ensuing kinetics of gene expression patterns, we performed RNA sequencing at earlier time points (4 hrs, 8 hrs and 12 hrs of induction), post expression of Cyb2SS and Cyb2ΔSS-fused PMD, DMMBP and MBP (as control of folded protein) proteins to target the proteins to IMS and matrix, respectively (Table S3). Principal Component Analysis (PCA) of the gene expression changes with respect to zero time, shows that the responses are dependent both on the type of misfolding property of the stressor protein and their localization in the mitochondrial sub-compartments (Figure 5A). At the earlier hours of proteotoxic stress, at both 4 and 8-hours post-expression of stressors, gene expression profiles for IMS-PMD and MM-PMD segregate along the first principal component axis suggesting that for the same stressor protein, a large part of the response is dictated by the localization of the misfolded protein within mitochondria (Figure 5A, left and middle panels). At later time point (12 hours post-induction) the points segregate primarily based on the extent of toxicity of stressor proteins expressed and its effect on cell growth along PC1 but segregates according to intra-mitochondrial localization of stressor along PC2 (Figure 5A, right panel).

**Figure 5:**
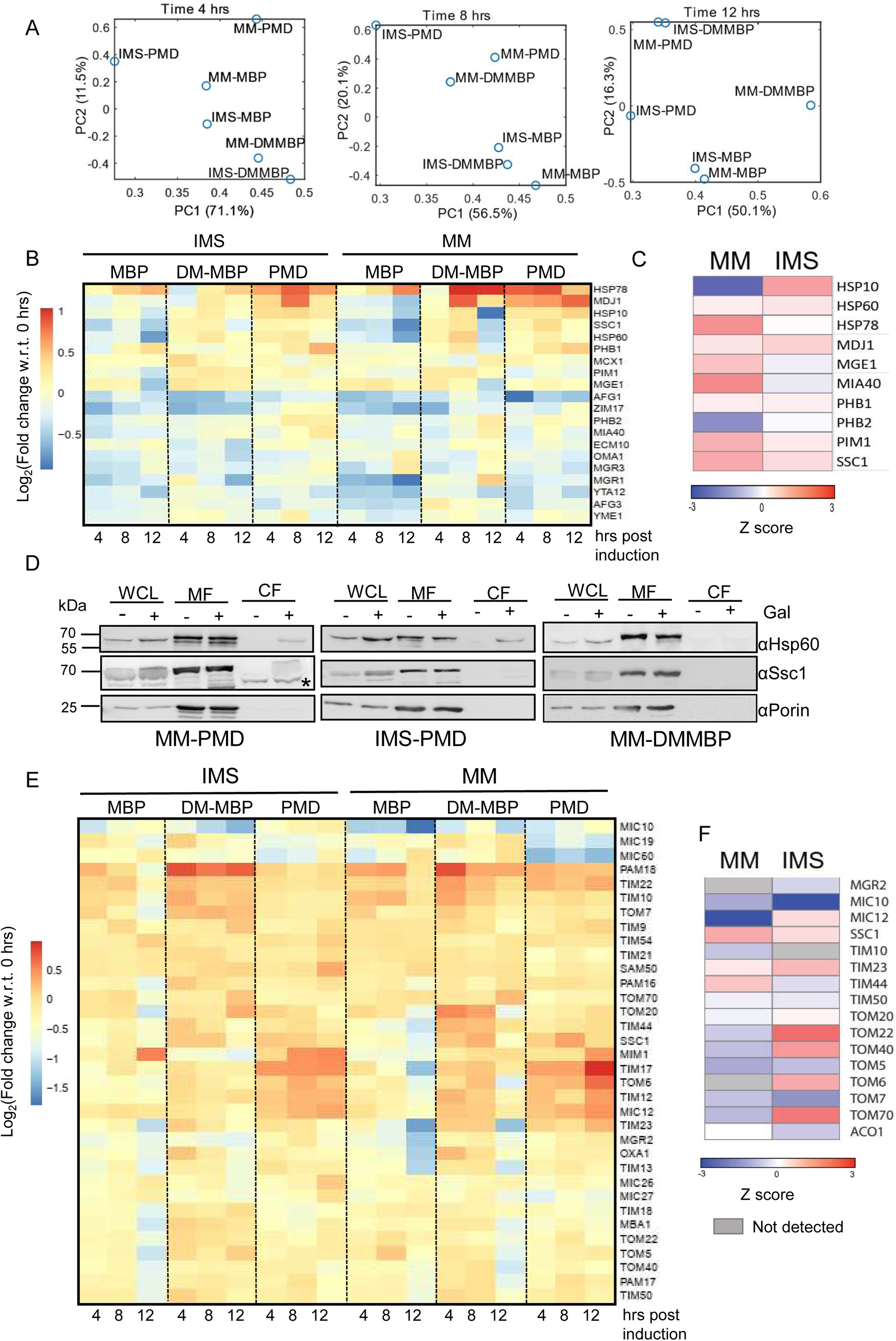
Time-dependent changes in stress response during mitochondrial proteotoxic stress. **A.** Principal components 1 (PC1) and 2 (PC2) obtained after Principal Component Analysis (PCA) of gene expressions are shown for each of the yeast strains in three different time points. The percentage variation explained by each of the PCs are show in the axis title. As evident, at 4th and 8th hour post-induction, IMS-PMD and MM-PMD are separated on PC1 as well as PC2 indicating that the responses are different when the same aggregation prone protein is targeted to mitochondrial matrix or IMS. Notably, IMS-DMMBP clusters along with matrix-targeted proteins indicating that the overall response is different for a soluble slow folding protein and an aggregation prone protein targeted to the IMS. However, there are similarities in a small subset of genes (as shown in subsequent sections) and the PCA is unable to capture this as the genes responsible for IMS proteostasis forms only a miniscule subset for all the genes and contributes less to the overall variation in the dataset. **B.** Gene expression changes mitoQC components of yeast strains with IMS and matrix (MM) misfolding stress due to overexpression of stressor proteins post 4 hrs, 8 hrs and 12 hrs of galactose induction are depicted as heatmap. Wt MBP has been kept as control of overexpression of a folded protein in the respective compartments. The fold changes of gene expressions in log2 scale in comparison to 0 time point of each strain have been plotted as heat map. **C**. Heatmap of Z-scores depicting protein-abundance of mitochondrial chaperones and proteases after 12 hours of expression of PMD protein in matrix (MM) or IMS (n=2). **D**. Left and middle panel: Western blot of mitochondrial matrix chaperones, Hsp60 and Ssc1 showing cytosolic accumulation of chaperones after 12 hours of misfolding stress by PMD protein in mitochondrial matrix (left panel) and IMS (middle panel). Western blot for same chaperones done under similar condition for MM-DMMBP strain which does not show any growth phenotype (right panel). Here, no cytosolic accumulation of Hsp60 or Ssc1 are observed. Lanes without galactose are shown as controls. Porin blot has been shown as marker for mitochondrial fraction. **E.** Gene expression changes of translocase (TOM, TIM, OXA and MICOS complexes) components of yeast strains with IMS and matrix (MM) misfolding stress are shown as heat map as described in panel A. **F**. Heatmap of Z-scores depicting protein-abundance of mitochondrial translocases after 12 hours of expression of PMD protein in matrix (MM) or IMS (n=2).

Subsequently, we analysed kinetics of expression changes of genes involved in mitochondrial protein homeostasis, biogenesis and functions with proteotoxic stress. First, we checked the kinetics of expression of mitochondrial chaperones and proteases with mounting of stress response. Core molecular chaperones of mitochondrial matrix like Hsp60 and Ssc1 (mitochondrial Hsp70) were upregulated, and highest upregulation was observed after 8 hours of induction of PMD or DMMBP protein expression which subsequently decreased in next 4 hours. Importantly, with expression of folded protein MBP, similar overexpression of *HSP60* or *SSC1* was not observed. This finding reiterated that the mitochondrial stress response is specifically mounted by the presence of misfolded or aggregation prone proteins and not by mere presence of any overexpressed protein. Notably, the amplitude of upregulation of these core molecular chaperones was only moderate. Other chaperones like, *MDJ1* and *HSP78* were substantially upregulated with the targeting of misfolded proteins in both the sub-compartments showing unique and distinct kinetics (Figure 5B). Notably, these two chaperones mainly Hsp78 was also upregulated with expression of wild type MBP indicative of overburdening of mitoQC machinery irrespective of presence of misfolding stress. The protein levels of the chaperones post 12 hours of induction of misfolding stress were in nice corroboration with the transcriptomics data (Figure 5C). At protein level too, upregulation of Hsp60 was only marginal and upregulation of Ssc1 was mainly prominent in matrix stress. Interestingly, matrix chaperones especially Hsp60 (occasionally Ssc1 too) showed cytosolic accumulation in strains with visible toxicity due to misfolding stress (MM-PMD, IMS-PMD) (Figure 5D, left and middle panel). Strains with no detectable toxicity and growth phenotype (MM-DMMBP) did not show any such cytosolic accumulation of chaperones (Figure 5D, right panel).

To test specific roles of mitochondrial chaperones during proteotoxic stress, we crossed the chaperone deleted (or depleted) strains with misfolded protein expressing strains. Among many such strains tested, only *ssc1*-knock-down (*SSC1*-DAMP denoted as *ssc1d*) showed further aggravation of misfolding induced growth phenotypes in DMMBP/PMD-IMS and PMD-MM strains indicating general importance of Ssc1 during protein misfolding stress in mitochondria irrespective of its location (Figure S5A). Interestingly deletion of J-domain cochaperone of Ssc1, MDJ1 (*mdj1*Δ), did not aggravate the growth phenotype further while deletion of another mitochondrial matrix Hsp70 (*ECM10* deletion exhibited alleviation of growth phenotypes in both IMS and MM-PMD strains (Figure S5A). The single deletion (or depletion strains) strains of chaperones did not show any growth phenotypes in absence of the stressor proteins (Figure S5D, left panel), indicating that alterations in the observed growth phenotype after crossing with the chaperone deleted-strains following proteotoxic stress is due to absence of modulatory role of these chaperones in the stress response pathways. Simultaneous to chaperones, effect of deletion of other major components of mitoPQC (mitochondrial Protein Quality Control), the mitochondrial proteases were checked. Crossing with deletion strains of mitochondrial proteases like m-AAA, i-AAA and lon proteases namely *yta12*Δ, *yme1*Δ, *pim1*Δrespectively (these deletions strains do not show any observable growth phenotype in absence of stressor protein expression; Figure S5D, middle panel), did not aggravate the phenotype of misfolding induced stress in mitochondria. This data indicates absence of significant contribution of these proteases, at least individually, in IMS or matrix stress response, during mitochondrial proteotoxic stress (Figure S5B). With genetic interaction with *cox14*Δ and *cox17*Δ (these deletions strains also do not exhibit any observable growth phenotype in absence of stressor protein expression; Figure S5D, right panel) strains (both genes were specifically upregulated in IMS-UPR), we observed further alleviation or aggravation in IMS or matrix stress, but the effects were variable between compartments and with different stressor proteins. Thus, their role as specific modulator of IMS-UPR remained inconclusive (Figure S5C).

As we observed cytosolic accumulation of mitochondrial matrix chaperones like Hsp60 or Ssc1 (Figure 5D) with severe stress, it pointed out a possible block in mitochondrial translocation following mounting of misfolding stress in the organelle. Thus, it was interesting to check the gene expression changes of the translocase components with stress. Upon analysing the expression of components of various TIM, TOM and MICOS complexes, we observed time dependent transcriptional upregulation of few translocase components of both outer and inner membrane. Components of different translocases like *TOM6, TIM17, TIM23, TIM12, MIC12* and *MIM1* were upregulated in a time-dependent manner in both IMS and matrix stress. All these subunits were more prominently upregulated in the later part after induction (8-12 hrs) of stress (Figure 5E). At protein level, we found several TOM complex components (Tom6, Tom40, Tom22, Tom70) to be specifically upregulated in IMS stress after 12 hours of stress with PMD protein (Figure 5F) in accordance with first set of transcriptomics data (Figure S4A). Notably, there was no prominent transcriptional downregulation of any of the TIM or TOM complex components indicating absence of any adaptive response to decrease mitochondrial translocation and curtail the incoming protein load during proteotoxic stress. We speculate that translocation block probably happens due to physical block of translocase pores due to decreased translocation rate or clogging of pores with misfolded/aggregated proteins and not due to transcriptional rewiring of translocase components. To assess whether mitochondrial import is hampered in stressed mitochondria, we did an import assay with previously described fluorescently-tagged COXIV pre-sequence peptide [32]. Non-stressed mitochondria isolated from wild type yeast cells grown in either in absence or presence of the inducer, imported the fluorescent peptide efficiently (Figure S6A, left panel). Mitochondria from IMS-DMMBP strain in absence of stress efficiently imported the peptide (Figure S6A, middle upper panel). Mitochondria isolated from the same strain after protein expression, showed fragmented mitochondria (Figure 3A, middle panel) but imported the peptide efficiently like non-stressed mitochondria (Figure S6A, middle upper panel). When we checked the physical interaction with the stressor proteins with TOM complex, we found Cyb2SS-fused DMMBP interacted with Tom40 (Figure S6B) by co-immunoprecipitation with DMMBP protein from isolated mitochondria from IMS-DMMBP strain. Despite such physical interaction of stressor proteins with TOM complex, mitochondria remain import competent in IMS-DMMBP strains. Mitochondria from induced IMS-PMD (Figure S6A, middle lower panel) and MM-PMD (Figure S6A, right panel) strains were extremely fragmented and imported the peptide less efficiently. Import of the peptide in induced MM-PMD mitochondria was least efficient (Figure S6A, right panel). This inefficient mitochondrial import of MM-PMD strain was also evident from partial accumulation of the PMD-protein in cytosol (Figure S1E) and cytosolic accumulation of matrix chaperones (Figure 5D, left panel) as shown before. Similar cytosolic accumulation of Hsp60 in IMS-PMD strain (Figure 5D, middle panel) can be explained by less efficient import of the peptide in these mitochondria. It is interesting to note that, stressed and fragmented mitochondria from IMS-DMMBP was efficient in import indicating nature of stressor proteins in same sub-mitochondrial compartment would determine the toxic phenotypes and the resulting stress response. Due to expression of large aggregate forming stressor proteins like PMD may lead to increased dwelling time of pre-proteins through TOM-TIM translocon pores leading to physical blockage of translocase channels. This ultimately mimic the cellular scenario as observed for mitochondrial precursor over-accumulation stress (mPOS) [33] especially at the late hours after commencement of misfolding stress.

Cytosolic accumulation of matrix chaperones following long-term mitochondrial proteotoxic stress precisely indicated mounting of a secondary cellular stress similar to mPOS or UPRam (Unfolded Protein Response activated by mistargeting of proteins) [33, 34]. A prominent network of upregulated cytosolic quality control machineries in long-term matrix stress by PMD protein indicated mounting of a cytosolic stress response (Figure S4B). Subsequently, we checked the gene expression levels of cytosolic protein quality control (cytoQC) machineries at different time points of stress. We observed upregulation of cytosolic chaperones like Ssa4, Hsp26, mainly in the late hours (8 hrs and 12 hrs post induction) of induction by stressor proteins PMD and DMMBP protein in matrix (Figure S6C). In contrast, only PMD-induced stress in IMS overexpressed the Ssa4 and Hsp26 but same was not observed with DMMBP stress in IMS. Chaperones like Ssa2, Hsp82 were found to be overexpressed in the early hours (4 hrs and 8 hrs post-induction) with expression of stressor proteins like DMMBP and PMD in matrix. These chaperones were similarly overexpressed with IMS-PMD but not with IMS-DMMBP stress (Figure S6C). In agreement with the transcriptomic data, at protein level too, many components of cytoQC like Ssa1, Ssa2, Ssa4, Hsc82, Hsp82, Ydj1, Hsp42, Hsp12, were substantially upregulated due to stress in mitochondrial matrix as well IMS (Figure S6D). This data indeed indicated mounting of a secondary cytosolic stress response most possibly but not exclusively due to inefficient mitochondrial import. As shown by in vitro import assays (Figure S6A), IMS-PMD and MM-PMD-stressed mitochondria are less efficient in import which is in corroboration to overexpression of cytosolic chaperones as an indicator of cytosolic stress. IMS-DMMBP despite showing toxicity, the stressed mitochondria show efficient import activity which may explain the absence of overexpression of cytosolic chaperones as shown by PMD-toxicity in both IMS. The overexpression of cytosolic chaperones in MM-DMMBP strain is puzzling as the strain do not show any toxic phenotype or problem in import. In contrast to mPOS and UPRam [33, 34], most of the proteasome components were repressed upon misfolding stress in IMS or matrix (Figure S6E).

Taken together, we show the time-dependent changes of expression of mitochondrial protein quality control machineries, translocase components during both IMS and matrix stress. Interestingly, all mitochondrial chaperones are encoded by nuclear genome and need to be translocated inside the organelle which becomes challenging during overwhelming stress leading to translocation block (as in case of MM-PMD and IMS-PMD strains). In consequence, cellular adaptive response to mitochondrial proteotoxic stress has evolved in a unique manner and does not prominently rely on prominent upregulation of mitochondrial chaperones or proteases, in contrast to ER-UPR [35, 36] or cytosolic heat shock response (HSR) [30, 37].

### Mitochondrial respiration is abrogated as an adaptive cellular response to proteotoxic stress in the organelle

In the transcriptome data, strikingly we find prominent repression of majority of the components of mitochondrial Electron Transport Chain (ETC) or respiratory chain complex (RCC) due to stress elicited by both Cyb2SS and Cyb2ΔSS-fused stressor proteins in IMS and matrix, respectively (Figure 6A). While the transcriptional repression is severe for majority of the components of complex III, V, II and half of the components of complex IV in MM-PMD induced matrix stress, the repression is not so prominent for components of complex III and V in IMS stress with the same stressor protein (Figure 5A). DMMBP elicited stress in IMS resulted repression of most of the components of complex II and half of the components of complex IV. Interestingly, proteotoxic stress in IMS by two different stressor proteins (DMMBP and PMD) of distinct (mis)folding properties showed different patterns of expression of some components of ETC (e.g., *QCR7/8/9* of complex III and *ATP15/17/4*, *TIM11* of complex V) indicating stressor-protein specific transcriptional stress response in the same mitochondrial sub-compartment (Figure 6A). DMMBP did not exhibit prominent repression of ETC components except few components of complex IV in matrix which corroborated well with no observed toxicity of this strain shown before (Figure 1H). Importantly NADH dehydrogenases like *NDE1/NDE2* or the subunits which are encoded by the mitochondrial genome like COB (component of complex III), *ATP6, ATP8, OLI1* (components of complex V) are not repressed suggesting that the primary route of response to mitochondrial stress is through the nuclear genome. Downregulation of most of the RCC components are recapitulated at the proteome level after PMD-elicited stress in matrix and IMS (Figure 6B, Figure S7A, table S4). Surprisingly, complex 2 components (Sdh1, Sdh2, Sdh3 and Sdh4) are found to be upregulated at protein level during matrix stress, the exact cause of this upregulation of complex 2 subunits remain to be explored.

**Figure 6:**
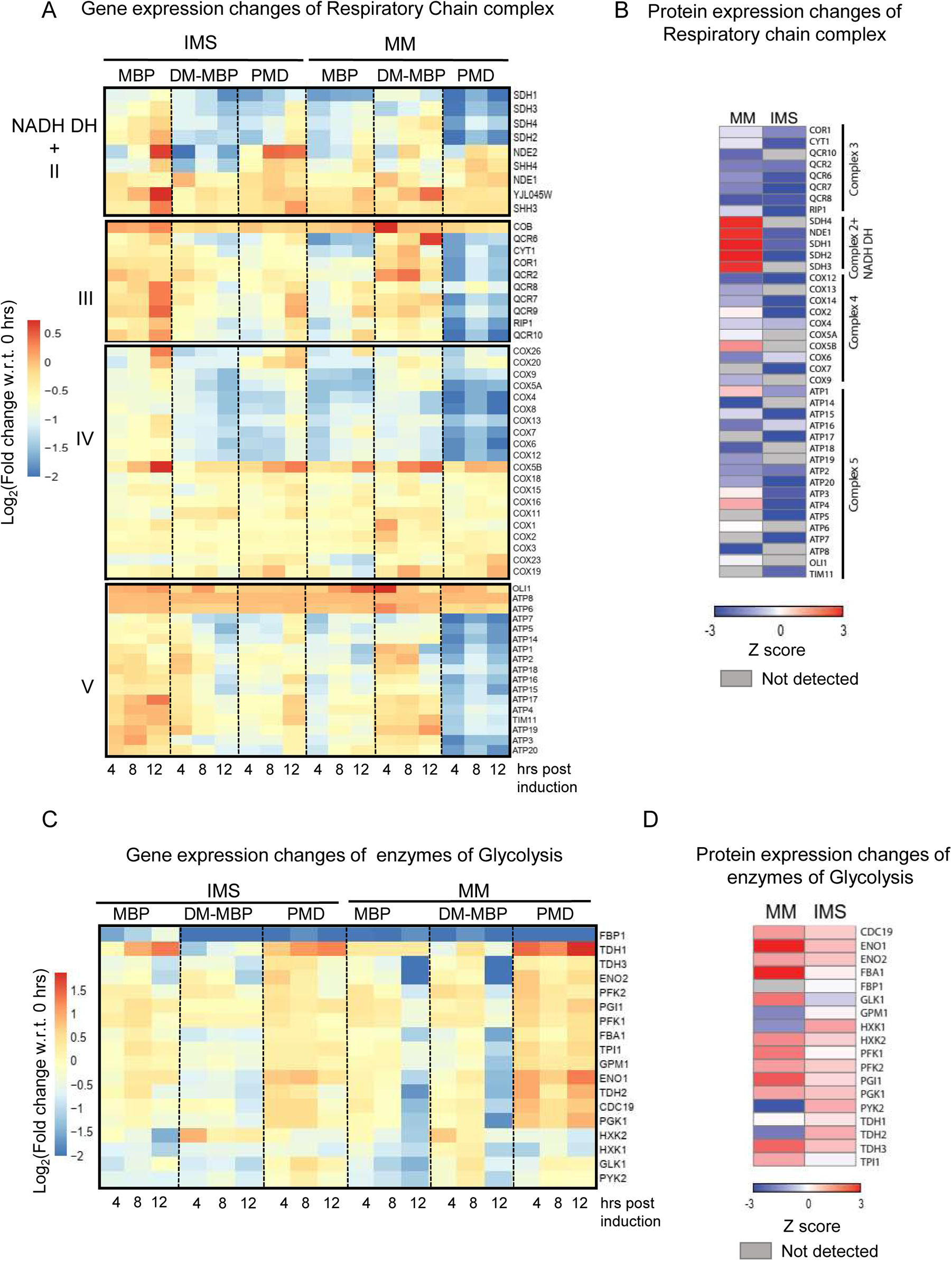
Components of respiratory chain complexes are substantially downregulated as an adaptive response mitochondrial proteotoxic stress. **A.** Gene expression changes of components of respiratory chain complex (or components of OX-PHOS complexes) of yeast strains with IMS and matrix (MM) misfolding stress due to overexpression of stressor proteins post 4 hrs, 8 hrs and 12 hrs of galactose induction are shown as heatmap. Wt MBP has been kept as control of overexpression of a folded protein in the respective compartments. The fold changes of gene expressions in log2 scale in comparison to 0 time point of each strain have been plotted as heat map. **B.** Heatmap of z-scores depicting protein-abundance of components of Respiratory Chain complex, after 12 hours of expression of PMD protein in matrix (MM) or IMS (n=2). **C.** Gene expression changes of components of Glycolysis with IMS and matrix (MM) misfolding stress due to overexpression of stressor proteins post 4 hrs, 8 hrs and 12 hrs of galactose induction are shown as heatmap similarly to panel A. D. Heatmap of z-scores depicting protein-abundance of components of glycolytic enzymes, after 12 hours of expression of PMD protein in matrix (MM) or IMS (n=2).

To determine whether abrogation of mitochondrial respiration by downregulation of OX-PHOS components is an adaptive response to mitochondrial misfolding stress, we forced the cells to undergo mitochondrial respiration in non-fermentable media in presence of misfolding stress. The growth phenotype due to misfolding stress deteriorated further in non-fermentable (glycerol) media in matrix stress indicating an adverse effect of forceful mitochondrial respiration during proteotoxic stress in the organelle (Figure S7C).

Furthermore, treatment with small molecules like ascorbate, which increases the cellular respiration specifically of the stressed cells (the molecular explanation of this effect of ascorbate needs further exploration) (Figure S7D), aggravates the toxic phenotype due to misfolding stress in mitochondria (Figure S7E). Collectively, we show that shutdown of mitochondrial respiration is implemented by yeast as an adaptive mechanism to misfolding stress in the organelle and forcing cells to continue respiration during proteotoxic stress proves damaging for the cells and the organism.

In the previous section, we have shown that yeast cells remain viable even after long-standing mitochondrial misfolding stress (Figure S2E) and after withdrawal of the stress cells grow like non-stressed cells without apparent growth phenotype (Figure S3A). We determined if cells adopt alternate energy metabolism strategy while suspending mitochondrial respiration to circumvent mitochondrial stress. On analysing the transcriptome (Figure 6C) and proteome data (Figure 6D and Figure S7B), we observed substantial upregulation of genes involved in glycolysis and the upregulation was more prominent during matrix stress. This finding aligns well with the altered respiratory status of mitochondria especially during matrix stress (Figure 3C-F) demanding activation of alternate energy metabolism pathways. Concordantly, TCA cycle components were found to be downregulated at the transcriptome as well as proteome level (Figure S7F and G). This also suggests that yeast cells bypass the mitochondrial misfolding stress by shifting the metabolism towards glycolysis. Taken together, we report that misfolding stress in mitochondria remodel the nuclear transcriptome to repress mitochondrial respiration and TCA cycle while activating alternate energy producing pathways like glycolysis. This rewiring is critical in maintaining cellular homeostasis during mitochondrial stress.

### IMS proteotoxic stress: Specific Role of TOM complex components

We have shown in the previous sections that upregulation of components of TOM complex are specifically evident upon Cyb2SS-fused-PMD eliciting IMS stress (Figure S4A) which is further validated at the protein level (Figure 5F). This result prompted us to do a systematic genetic interaction with deletion strains of TOM complex components. We found that deletion of *TOM6, TOM7 or TOM22,* aggravated the growth phenotype of IMS-PMD strain, (Figure 7A). Similar effect of deletion of these TOM complex subunits (*TOM6, TOM7*) are also observed in IMS-DMMBP strain (data not shown). Notably, none of the single deletion strains of TOM subunits show any growth phenotype in absence of the stressor proteins (Figure 7C) indicating important modulatory role of these TOM complex components during IMS stress response. In contrast, deletion of other outer mitochondrial membrane protein like porin (*POR1*), does not affect the IMS-UPR (Figure 7A). Notably, the effect of deletion of these TOM subunits is exclusive for IMS stress and does not affect the phenotype of matrix stress (Figure 7B) as observed during IMS stress. This clearly indicate the importance of TOM complex in modulating IMS proteotoxic stress response or IMS-UPR.

**Figure 7:**
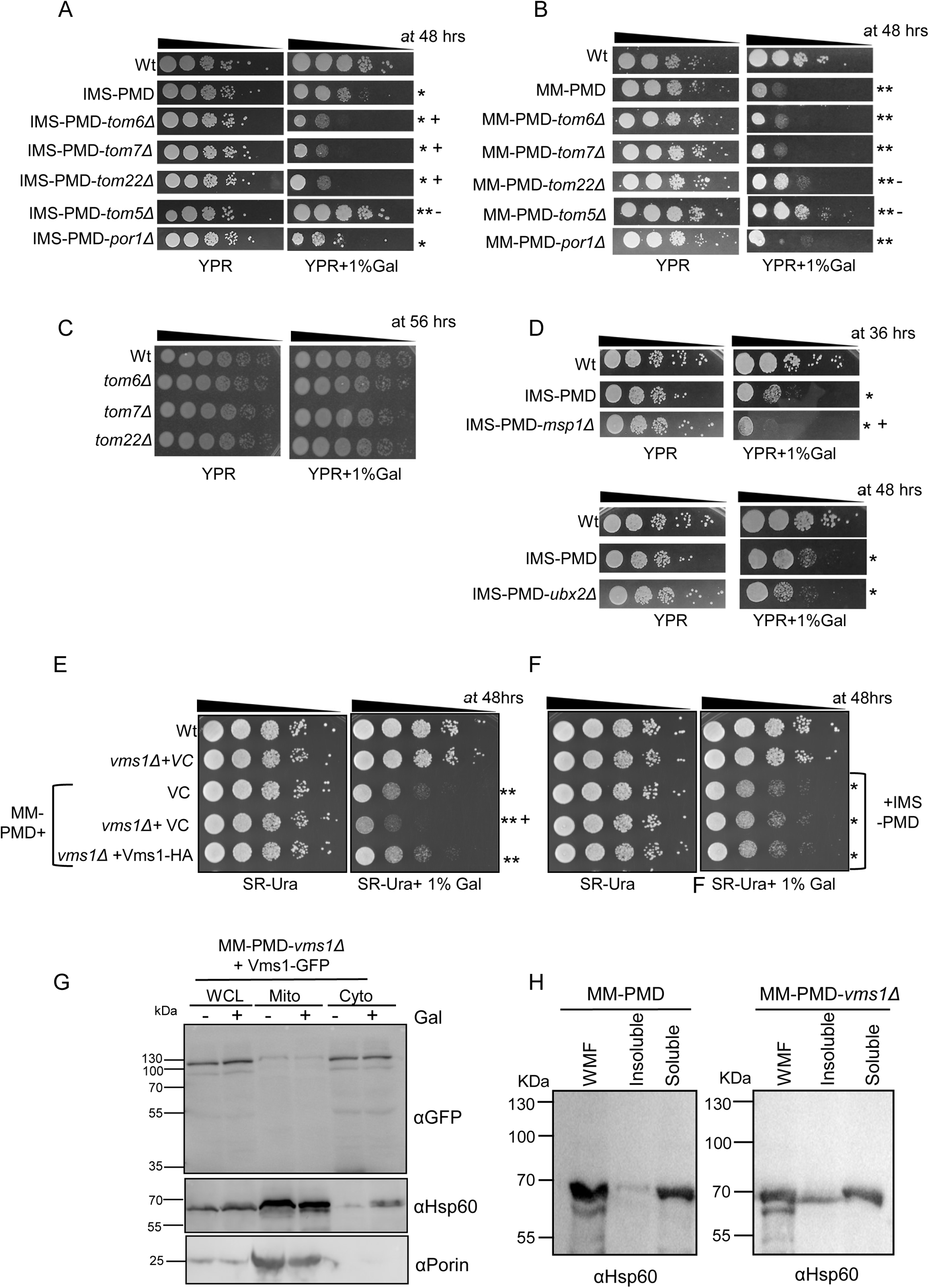
Identification of IMS and matrix-stress response modifiers by genetic interaction. **A.** Drop-dilution assay as of yeast strains containing single deletion of different single components of TOM complex expressing mitochondrial IMS targeted stressor protein PMD in presence and absence of the inducer, as described in Figure 1F. “*” indicates spots with visible growth phenotype in case of IMS-PMD. “*+” indicates spots with further aggravation of growth phenotype due to deletion of TOM components, *TOM6* or *TOM7* or *TOM22*. “*-” indicates spots with alleviated growth phenotype due to deletion of *TOM5*. Porin (*POR1*) deletion was taken as control. The spots are 1:10 serial dilutions and time mentioned in hours at the upper right corner of the panel indicates the time of incubation of the spotted plates at 30°C before taking pictures. All rows of spots were taken from same plate but for better alignment of spots for easy comparison amongst strains, the rows are shown as separate strips. **B**. Similar to panel A, effect of deletion of TOM complex single components are checked during matrix stress by drop-dilution assay of yeast strains containing single deletion of different single components of TOM complex containing mitochondrial matrix targeted stressor protein PMD (MM-PMD) in presence and absence of the inducer. “**” indicates spots with visible growth phenotype of MM-PMD strain. As there was no further aggravation of the growth phenotype of MM-PMD strains when combined with single Tom components deletion (*TOM6* or *TOM7*), the phenotypes are indicated as “**”, similar to MM-PMD strain. “*-” indicates spots with alleviated growth phenotype due to deletion of *TOM5* or *TOM22*. The spots are shown as strips from same plates as explained in panel A. **C. D.** Drop-dilution assay as of yeast strains having deletion of either *MSP1* (upper panel) or *UBX2* (lower panel) in the strains expressing mitochondrial IMS targeted stressor protein PMD in presence and absence of the inducer, as described in Figure 1F. “*” indicates spots with visible growth phenotype in case of IMS-PMD. “*+” indicates spots with further aggravation of growth phenotype due to deletion *MSP1*. The spots are shown as strips from same plates as explained in panel A. **E-F**. Drop-dilution assay as of MM-PMD (panel E) and IMS-PMD (panel F) strains with *vms1*Δin presence and absence of the inducer. Vms1-HA expressed under endogenous promoter from a centromeric Ura plasmid has been shown to complement the growth defect observed in MM-PMD-*vms1*Δ. **+ indicates spots of MM-PMD-*vms1*Δ with further aggravation of growth phenotype of MM-PMD strain due to deletion of *VMS1*. **G**. Western blot of Vms1-GFP protein expressed in MM-PMD-*vms1*Δ cells. Subcellular fractionation was done as described in methods section and equal amount of total protein from the whole cell lysate (WCL), mitochondrial fraction (Mito) and cytosolic fractions (Cyto) were loaded and probed with anti-GFP antibody to detect the GFP-tagged Vms1 protein. Western blot from same fractions were also probed with Hsp60 and porin specific antibodies. **H.** Mitochondria of MM-PMD and MM-PMD-*vms1*Δ were isolated after expressing the proteins for 12 hours with 1% galactose in YPR. Isolated mitochondria were subjected to solubilization with 0.1% Tx-100 in SH buffer and was separated into detergent soluble and insoluble fractions by high-speed centrifugation. The pellet (insoluble fraction) and supernatant (soluble fraction) were loaded, and western blot was performed with anti-Hsp60 antisera. Part of Hsp60 protein is found in detergent-insoluble fraction in MM-PMD-*vms1*Δ mitochondria which remains completely in soluble fraction in absence of deletion of *VMS1* (left panel and is also shown in Figure 2F and I).

Recently, identification of various stress response pathways like mitochondrial compromised protein import response (mitoCPR) [17], mitochondrial translocase associated degradation (mitoTAD) [38] have proved the importance of TOM receptor subunits like Tom70 in mitochondrial and overall cellular protein homeostasis. During mito-CPR, Tom70 recruits Msp1 (an AAA-ATPase with extractase [14–16] activity for mislocalized ER tail-anchored proteins to mitochondrial outer membrane (MOM) to maintain MOM proteostasis) for clearing out the jammed TOM-complex with overexpressed bipartite signal sequence containing preproteins [17]. On the other hand, Ubx2, a bridging factor for Endoplasmic Reticulum Associated Degradation (ERAD), is recruited to outer mitochondrial membrane through Tom70 and performs continuous surveillance of TOM complex by mito-TAD under non-stress conditions and maintains protein homeostasis [38]. Our model of IMS misfolding stress (imparted by bipartite signal sequence containing stressor proteins) would mimic the cellular response like mitoCPR especially during long standing chronic stress. To check the role of *MSP1* or *UBX2*, we deleted the corresponding genes from IMS-PMD and MM-PMD background. Deletion of *MSP1* prominently aggravated the growth phenotype of IMS-PMD strain (Figure 7D, upper panel). *ubx2*Δalso aggravated the phenotype of IMS-PMD although to a much lesser extent than *msp1*Δ (Figure 7D, lower panel). In contrast, *msp*1Δ or *ubx2*Δ did not impart any additional effect on MM-PMD phenotype indicating important role of Msp1 or Ubx2-governed proteostasis pathways during IMS proteotoxic stress (Figure S8A).

To check any redundancy among the TOM core subunits in the stress response modulatory function, we overexpressed the individual components of TOM core-complex (*TOM6, TOM22, TOM40*) from yeast ORF overexpression library in the background of IMS-PMD-*tom6*Δ and IMS-PMD-*tom7*Δ. Additionally we also overexpressed *TOM70* in the same strains to check whether increasing Tom70-mediated response pathways (through Msp1, Ubx2) by its overexpression can compensate for loss of modulatory function of TOM core subunits. Interestingly, overexpression of *TOM6, TOM22, TOM40* efficiently rescued the growth defects due to *tom6*Δ (Figure S8B) or *tom7*Δin the IMS-PMD strain (Figure S8C), although *TOM70* overexpression was unable to do so. This data indicates that TOM core complex subunits possess modulatory functions in IMS stress responses, and this function is redundant among Tom6, Tom7 and Tom22.

### Matrix proteotoxic stress: Specific Role of Vms1

Upon systematic genetic interactions with various deletion strains of matrix stress specific upregulated genes, we found role of *VMS1* to be critical for mito-matrix-UPR. Vms1 is known as a key component of Mitochondria Associated Degradation (MAD) process [22, 39, 40] and an important player of Ribosome Quality Control pathway [18–21, 41]. Deletion of *VMS1* in the matrix (Figure 7E) and IMS (Figure 7F) misfolding strains, revealed its prominent role in matrix misfolding stress. The growth phenotype due to matrix stress worsened in absence of *VMS1* ((Figure 7E), while similar effect was not apparent in IMS stress (Figure 7F). The *vms1*Δstrain without expression of misfolded proteins did not exhibit any growth phenotype [*vms1*Δ+VC (vector control) in Figure 7E and F) indicating that the aggravation of toxic phenotypes of MM-PMD-*vms1*Δ strain compared to MM-PMD strain is due to the absence of modulatory functions of *VMS1* on matrix stress response pathways.

As Vms1 contains a Mitochondria Targeting Domain (MTD) which naturally remains shielded and the protein remains in cytosol, in some stress conditions like in oxidative stress to the protein was shown to localize in mitochondria [42]. Thus, it was interesting to check whether the protein alternately localizes to mitochondria during proteotoxic stress. To check the localization of Vms1 during matrix misfolding stress, we expressed Vms1-tagged to C-terminal GFP under its native promoter in the MM-PMD­*vms1*Δ cells. We found two bands for Vms1-GFP of ~115kDa and ~90kDa (molecular weight with the GFP-tag) (Figure 7G). Bands of similar size were also observed with Flag-tagged Vms1 (data not shown). Both forms of the protein were majorly enriched in cytosolic fraction and both forms were detected in the mitochondrial fraction only in minor amounts (Figure 7G). In agreement with the western blot result, Vms1-GFP was found to be majorly localized in cytosol and no significant co-localization was observed with mitochondria (Figure S8D). This data indicates that cytosolic Vms1 exerts its modulatory function during mitochondrial matrix stress.

Previous study showed that CAT-tailed mitochondrial pre-proteins accumulate and aggregate in absence of antagonistic effect of Vms1 on Rqc2’s CAT-tailing activity [18]. We speculate that this antagonistic activity of Vms1 on Rqc2 would be extremely important during matrix stress as uncontrolled CAT-tailed proteins may pose further problem to already stressed mitochondria.

Uncontrolled CAT tailing by Rqc2 in absence of *VMS1* thus is expected to increase protein aggregation in the mitochondrial matrix [18]. To check whether mitochondrial proteins are more aggregation-prone during matrix proteotoxic stress when combined with *vms1*Δ, we solubilized the mitochondria with detergent (Tx-100) isolated from MM-PMD and MM-PMD-*vms1*Δstrains after expressing the PMD protein. We checked the distribution of Hsp60 protein (which is shown before to be exclusively present in detergent-soluble mitochondrial fraction even in presence of mitochondrial matrix proteotoxic stress, Figure 2E and H) between detergent-soluble and detergent-insoluble fractions from MM-PMD and MM-PMD-*vms1*Δmitochondria. Interestingly, a substantial amount of Hsp60 protein was found in the detergent insoluble fraction of MM-PMD-*vms1*Δin contrast to no discernible Hsp60 in the insoluble fraction of MM-PMD mitochondria (Figure 7H). This data indicates that indeed *vms1*Δcombined with matrix proteotoxic stress may pose great threat of aggregation propensity of mitochondrial proteins leading to severe mitochondrial stress.

Other than regulating CAT tailing activity of *RQC2* in RQC process, *VMS1* was previously implicated in mitochondrial respiration [18]. Since we found mitochondrial respiration is suspended as an adaptive response to matrix stress, we wanted to check if Vms1 possesses any role in matrix stress response by modulating mitochondrial respiration. We henceforth measured the OCR of *vms1*Δcells which showed less OCR compared to the non-deleted strain in absence of misfolding stress (un-induced MM-PMD-*vms1*Δ cells and MM-PMD cells) (Figure S8E). Surprisingly, OCR was increased in induced MM-PMD-*vms1*Δ cells compared to MM-PMD cells (Figure S8E). This result indicates that *VMS1* play an important role in cellular adaptive response of limiting the mitochondrial respiration during misfolding stress. This role of *VMS1* corroborates well with the observed growth phenotype aggravation of MM-PMD-*vms1*Δ cells during stress (Figure 7E).

Taken together, we show that matrix misfolding stress when combined with absence of Vms1, becomes severely detrimental due to enhanced aggregation of mitochondrial proteins like chaperones which cause additional problems to previously-stressed mitochondria due to prevailing proteotoxic stress by accumulation of stressor proteins in the organelle. Additionally, by a yet unknown mechanism Vms1 assists in cellular adaptive response to suspend mitochondrial respiration during matrix stress.

## Discussion

In recent years, understanding mitochondrial proteostasis and communication of mitochondria with other sub-cellular compartments during stresses, have gained much attention. Extensive research in the related fields have yielded the discovery of novel stress response pathways like mitoUPR [7, 9, 43], MAGIC [4], mPOS[33], UPRam [34], mitoCPR[17, 44], mitoTAD[38], MISTERMINATE [45, 46], early mitoUPR [47] which have shed lights on extremely specialized and unique cellular pathways to thwart mitochondrial or mitochondria-associated stresses. In the current study, we have moved a step further beyond generalized mitochondrial stress and have asked whether cellular response can be distinct in response to localized stresses within mitochondrial sub-compartments. By fusing various stressor proteins to well-described IMS and matrix-targeting signal sequences, indeed, we show that apart from shared stress responses, some distinct stress responses specific for mitochondrial IMS or matrix can be captured. Importantly, we show that mere overexpression of folded proteins by the same inducer does not noticeably alter mitochondrial dynamics, or the organelle function and cells do not elicit canonical stress response after overexpression of folded proteins. Interestingly, none of the canonical mitochondrial chaperones are overwhelmingly expressed post misfolding stress either at RNA or protein level. We speculate that cells have evolved rather unconventional stress response mechanism for this unique organelle. As the mitoPQC components are exclusively encoded by the nuclear genome, it will be a challenging task to import the chaperones and proteases by an energy-expending process, especially those belonging to mitochondrial matrix, to mitigate the stress. Furthermore, we show that mitochondrial respiration is abrogated as an adaptive response due to stress elicited in both IMS and matrix. This suspended mitochondrial respiration compounded with physical block of the translocases could arise due to strong aggregate forming stressor (like PMD protein) during misfolding stress would lead to inefficient translocation of mitoPQC components, so producing these quality control machineries become a futile exercise. In such a scenario, mitochondria need to circumvent the stress situation and restore homeostasis by not relying on its own quality control machinery, rather it has evolved non-canonical pathways to handle misfolding stress. Import across the outer membrane being energy-independent, outer mitochondrial membrane associated quality control is easy to implement, at least for proteotoxic stress in IMS. We show that components of TOM complex are crucial during IMS stress. We speculate that a longstanding IMS stress would mimic responses similar to mitoCPR involving Tom70 and Cis1/Msp1 mediated response to restore the homeostasis. Indeed, we show that Msp1 and key regulator of mitoTAD pathway, Ubx2 are specifically important during IMS stress. Additionally, TOM core complex subunits (*TOM6*, *TOM7* and *TOM22*) exert specific modulatory role in IMS proteotoxic stress response (Figure 8).

**Figure 8:**
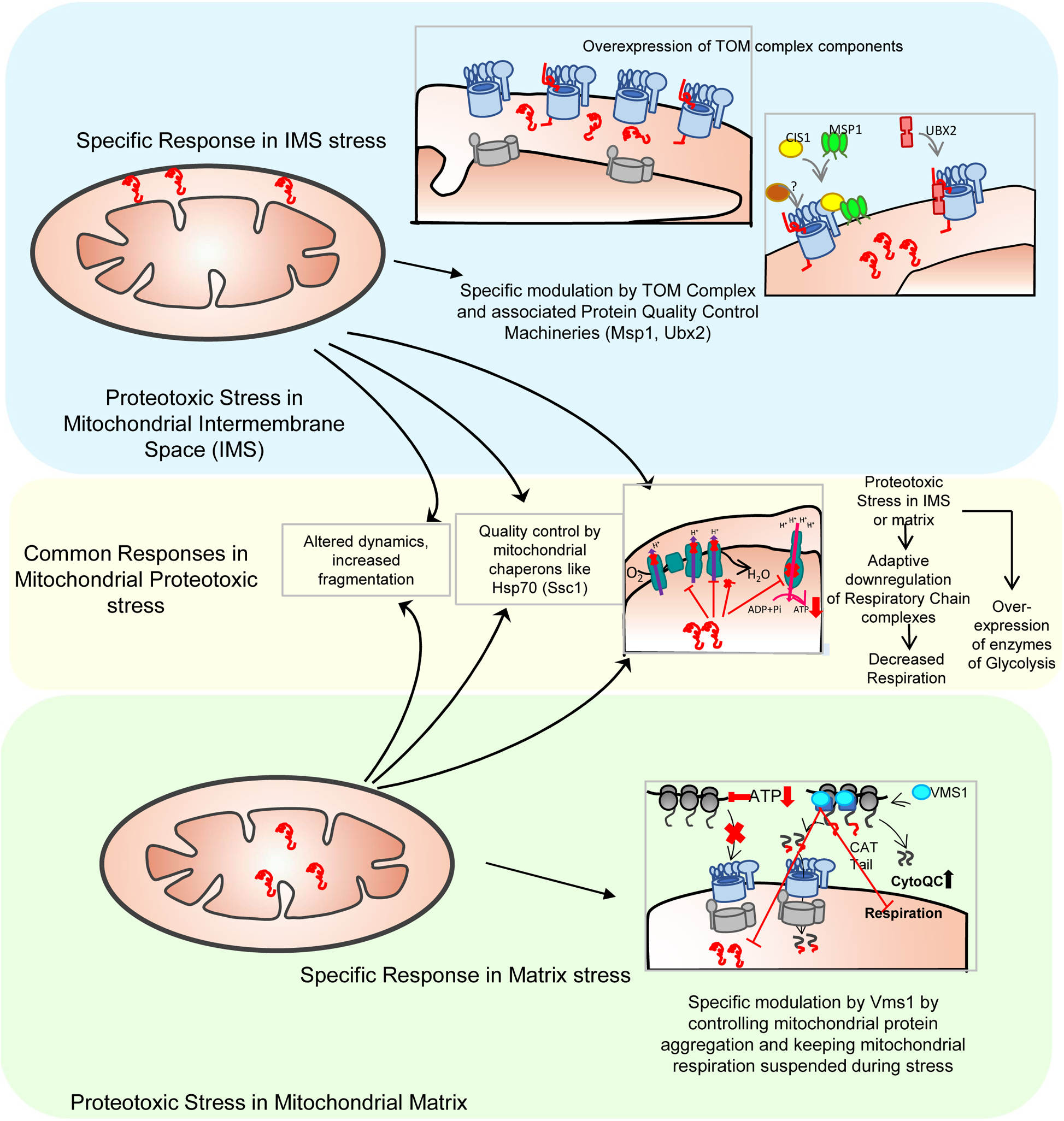
A comprehensive model of general and specific stress responses during proteotoxic stress in Mitochondrial sub-compartments. **Blue box:** Misfolded or aggregated protein from various sources, e.g., cytosolic proteins which needs to be cleared by MAGIC, or mis-targeted proteins of neighbouring organelles like ER and cytosol or endogenous mutated or misfolded proteins accumulate in mitochondrial IMS and stress is mounted. As a response to IMS proteotoxic stress, TOM complex components are overexpressed. TOM-core complex components exert modulatory function by a yet unknown mechanism in restoring the IMS homeostasis. TOM-complex associated protein quality control machineries like Msp1 and Ubx2 also play important role during IMS stress. **Yellow box** in the middle summarizes the global stress response due to proteotoxic stress within mitochondria irrespective of its location within matrix or IMS. Lower **Green box** summarizes two-way modulatory function of Vms1 during matrix proteotoxic stress. The exact molecular mechanism of TOM subunits or Vms1 during IMS or matrix specific stress, remains to be explored in detail.

In case of matrix stress, the response prominently displays the upregulation of mitoQC components (mitochondrial chaperones and co-chaperones) mitochondria-associated Ribosome Quality Control (mito-RQC) and cytosolic quality control (components of HSR). We show that MAD (Mitochondria Associated Degradation) and RQC component Vms1 (Figure 8), plays crucial role during matrix proteotoxic stress by keeping a check on the load of aggregated proteins on mitochondrial matrix by a yet unknown mechanism. Whether the mito-protein aggregation prevention activity of Vms1 during proteotoxic stress is due to its antagonizing activity on Rqc2’s CAT-tailing activity on mitochondrial precursors or beyond it, remains to be explored in-depth in future. We further show that Vms1 is important to limit the mitochondrial respiration during proteotoxic stress which is suspended as an adaptive stress response to mitochondrial proteotoxic stress. Additionally, we found many components of cytosolic quality control are upregulated during long standing mitochondrial stress indicating important role of cytosolic quality control machinery in protecting the stressed organelle. Our current findings and previous works describing stress response pathways like mitoCPR [17, 44], mitoUPR [7, 9, 10], MAGIC [4], mitoTAD [38], mito-protein induced stress by clogger proteins [48] etc., altogether indicate a meticulous symbiosis between mitochondria and cytosol, especially under situations of proteotoxic stress within and around the organelle. In effect, we see IMS-UPR and mito-matrix-UPR successfully tackle the stress with no significant cell death despite severe mitochondrial fragmentation and abrogated respiration, rather cell growth and mitochondrial dynamics are restored efficiently after withdrawal of stress. This capacity of handling mitochondrial proteotoxic stress is similarly efficient in yeast as well as in human cells. Whether specific modulators of IMS-UPR (specific components of TOM complex) and mito-matrix-UPR (Vms1) as found in yeast are conserved across eukaryotes, remains to be checked.

## Acknowledgements

We thank Dr Dejana Mokranjac for sharing the Cyb2-DHFR plasmid. We thank Prof. Jared Rutter for sharing the Vms1 and Vms1 truncation mutant plasmids. We thank Dr. Claes Andreasson for sharing the mutC-DHFR plasmid. We thank Dr Deepak Sharma for sharing yeast overexpression plasmids. Prof Patrick D’Silva is acknowledged for sharing the Tim23 antibody. We thank Dr Kausik Chakraborty for sharing the YMJ003 yeast strain and R script for RNA sequencing data analysis. We thank Aseem Chaphalkar for the UPRE-GFP FACS measurement. We acknowledge Gopal Gunanathan Jayaraj for help in RNAseq library preparation, preliminary data analysis and for critical comments on the manuscript. We acknowledge core Imaging facility and central Instrumentation facility of CSIR-IGIB. Mr. Manish Kumar at CSIR-IGIB and SNU imaging facility is acknowledged for help in imaging data analysis. We thank Monika Verma and Nishtha Bhargava and for critical comments on the manuscript. KBN acknowledges CSIR SRF grant [31/43(350)/2017-EMR-I], PP acknowledges the DST WosA grant (SR/WOS-A/LS-99/2014). PM acknowledges SERB NPDF grant (SERB/F/4161/2018-2019). RS acknowledges ICMR SRF grant (2019-6710/CMB/BMS) and SNU PhD fellowship. MA acknowledges SNU PhD fellowship.

## Availability of data and material

The transcriptomics data (RNAseq) data have been deposited in Bioproject Repository with the accession numbers PRJNA746123 (time-dependent transcriptomics data) and PRJNA747162 (Z-score based analysis of transcriptomics data). The mass spectrometry proteomics data have been deposited to the ProteomeXchange Consortium via the PRIDE partner repository with the dataset identifier PXD027216.

## Consent for publication Competing interests

The authors declare that there are no competing financial interests

## Author Contribution

The work was conceived by KM. All yeast strains were generated by KBN and PP. Yeast experiments were done by KBN, PP and RS. Biophysical characterization of PMD protein was done by KM. RNA sequencing library preparation and sequencing run was performed by AG. Data analysis was done by AG and KM. Samples for proteomics were made by RS, proteomics run, and data analysis were done by SM, KRK and SR. PMD protein simulation and secondary structural analysis were done by AR. Mammalian cell experiments were done by MA and PM. KM analysed the data and wrote the manuscript.

## METHODS

Supplementary Table 1 contains the List of yeast strains, plasmids, primers and Antibodies and other reagents used for this study

**Strains used:** S. cerevisiae strain YMJ003 (MATαhis3Δ1 leu2Δ0 met15Δ0 ura3Δ0 LYS+Δcan1::STE2pr-spHIS5 Δlyp1::STE3pr-LEU2 cyh2 Δura3::UPRE-GFP-TEF2pr-RFP-MET15-URA3. BY4741 (MATa, his3Δ1; leu2Δ0; met15Δ0; ura3Δ0) [49] and E.coli-DH5αstrains were used for this study. All the transformations and plasmid preparations were performed by standard method, which was adopted from the previous studies. Homologous recombination method was adapted for the generation of different compartment specific misfolding induced strains (Supplementary Table 1).

### Construction of plasmids for homologous recombination to generate compartment specific misfolded protein strains in yeast

The construct of different compartment specific misfolded protein, the desired gene is amplified and cloned in *ApaI* and *AvrII* site of pYMN23 plasmid (Supplementary Table 1). The mitochondrial targeting signal sequence of yeast Cytochromeb2 (cyb2) was amplified from Cyb2-DHFR plasmid and was cloned upstream of the ORF to target the protein into mitochondrial inter-membrane space and the deletion of 19 amino acid membrane sorting signal (Cyb2Δ19) was done by overlap PCR from the Cyb2 sequence to target the proteins to mitochondrial matrix. The whole construct is under the control of galactose inducible promoter. The whole DNA cassette was amplified by using Kapa HiFi DNA polymerase and transformed in yeast yMJ003 strain. Further, the integration of the DNA cassette at correct locus was confirmed by genotyping (plating on URA −/+ plates) and PCR based methods (using upstream and internal primers). Using similar methos, different constructs were prepared for different misfolded proteins which targets into different compartments (Supplementary Table 1). pVT100U-mtGFP plasmid was used as a fluorescent marker for mitochondria. For the localization study, misfolded protein (PMD) is tagged with GFP and integrated in the background strain as above.

### Growth conditions and drop dilution assay

All the yeast strains are maintained at 30°C in the commercially available YEPD (1% Yeast extract, 2% Peptone and 2% Dextrose) medium in the presence of nourseothricin antibiotic (cloNAT) antibiotic selection. All the strains are grown in poorly-fementative YPR (1% Yeast extract, 2% Peptone and 2% Raffinose) medium overnight and re-inoculated in the same medium with the initial OD_600_ of 0.1. For drop dilution assay, the different strains were grown in YPR medium till 0.4-0.6 and serially diluted and spotted on different plates containing YP along with 2% dextrose (YPD) or 2% raffinose with (YPR+ gal) and without (YPR) 1%galactose or 3%glycerol (YPG), or for specific treatment with 10mM ascorbate.

### Crossing of misfolded protein expressing yeast strains with deletion strains from YKO library

Yeast strains carrying misfolded protein (referred nomenclature as mentioned in Supplementary Table 1) and genotype (*MAT*α *his3*Δ*1 leu2*Δ*0 met15*Δ*0 ura3*Δ*0 LYS+*Δ*can1::STE2*pr-*spHIS5* Δ*lyp1::STE3*pr-*LEU2 cyh2* Δ*ura3*::UPRE-GFP-*TEF2*pr-RFP­*MET15-URA3*::signal sequence misfolded protein-NAT derived from S288C), referred to as query strain here onwards, were crossed with knockout strain in BY4741 background (MATa, *his3*Δ*1*; *leu2*Δ*0*; *met15*Δ*0*; *ura3*Δ*0*;gene::KANMX) (Yeast KO Library, Mat A complete set, Thermofisher Scientific cat no. 95401.H2) as described previously [50]. Briefly, query strain and single KO strain were cultured in YPD media till saturation. 5ul from saturated culture of each strain were inoculated together into 400 µl of YPD in 2.2 ml 96 well deep-well plates and were co-cultured overnight for mating at 30°C, at 200 rpm continuous shaking in a shaker incubator. For Mat a/ Mat α diploid selection, 10 µl of mated culture were re-inoculated in in 2.2 ml 96 well deep-well plates containing 400 µl of diploid selection medium (SD media with glutamic acid and without lysine, arginine and leucine) with antibiotics Nourseothricin 100 µg/ml and Geneticin 200µg/ml. The culture plate was incubated for 48 hours at 200 rpm continuous shaking at 30°C. For sporulation, 10 µl of diploid cells were re-inoculated in 400 µl of sporulation medium (1% Potassium acetate, 0.1% Yeast extract, 0.05% Glucose, 0.1% Amino acid supplement powder from mixture containing 2 g histidine, 10 g leucine, 2 g lysine, and 2 g uracil) in 2.2 ml 96 well deep-well plates and the plates were incubated at 25° in static condition for 5 days. From sporulation media, 10 µl of culture was seeded in haploid selection media (SD without leu/Arg/Lys) along with canavanine and thialysine to select meiotic cells in Mat α background and plates were incubated at 30°C for next 2 days. Second round of haploid selection was done in similar manner in presence of antibiotics Nourseothricin and Geneticin. After the second round of haploid selection, strains were inoculated for genomic DNA isolation and were confirmed for KANMX cassette integration, gene deletion and mating type PCR.

### Confocal microscopy imaging of yeast cells

Yeast cells were grown in YP-raffinose till it reaches the OD of 0.5 and the misfolded proteins were induced with addition of 1%galactose and further grown at 30C for 6-8hrs. Cells were harvested and washed once with 1XPBS. Cells were fixed with 3.5% formaldehyde solution and coated on Concanavalin A slides and allow 15mins for the cells to adhere on the slide and examined under confocal microscope. Images were captured in different positions of a slide and Z-series was acquired in each position. For the live-cell imaging, cells were coated on thin agarose gel pads and examined under microscope for different time-points.

### Imaging of yeast mitochondria

Cells were transformed with pVT-100U-mtGFP and selected on SD-Ura plates. For microscopy, cells were grown in synthetic raffinose broth (SR-Ura medium) till the OD_600_ of 0.5. Cells were then induced with 1% galactose and were further grown at 30°C for 8-12hrs and then washed with 1X PBS and coated on concanavalin A slides. Imaging was done in Leica TCS SP8 confocal microscope.

### Purification of mitochondria from yeast cells

Purification of yeast mitochondria was performed using standard protocol. Briefly, cells were grown in YP-raffinose medium till the OD6_00_ of 0.4-0.6 and the strains were induced with 1% galactose and incubated at 30°C for 8hrs. Cells were harvested at 4,400xg and washed once and checked the wet weight of the cells. Then cells were suspended in 0.5g/ml of DTT buffer [100mM Tris/H_2_SO_4_ (pH 9.4), 10mM DTT] and incubated at 30°C for 30mins. Cells were spun down at 4,400xg and suspended in zymolyase buffer [20mM potassium phosphate (pH 7.4), 1.2 M sorbitol along with 2.5mg/gm Zymolyase 100T] and incubated at 30°C for 1hr. Cell wall digestion was checked in spectrophotometer. Further cells were pellet down at 3000xg at 4°C and re-suspended in 0.5mg/ml homogenization buffer [10mM Tris/HCl (pH 7.4), 0.6 M sorbitol, 1 mM EDTA, 0.2% (w/v) BSA] and homogenized 15 times by using Dounce homogenizer. The ruptured cell debris were collected by spin at 3000xg for 5mins at 4°C. The collected supernatant was spun down in high-speed at 12,000xg for 15mins at 4°C. The sedimented crude mitochondria was suspended in homogenization buffer. The slow speed and subsequent high-speed centrifugation steps were repeated as described above. The resultant pellet contains crude mitochondria. Further the purified mitochondria was prepared by sucrose-gradient ultra-centrifugation. The different percentage of sucrose gradient was prepared in EM buffer [10mM MOPS/KOH (pH 7.2), 1mM EDTA]. First, 1.5ml of 60% sucrose in Beckman ultra-clear centrifuge tube was taken. Next, it was overlaid with 4ml of 32%, 1.5ml of 23% and 1.5ml of 15% sucrose to get the different gradients. The crude mitochondria were placed on top of 15% sucrose and centrifuge in Beckman SW41Ti swinging bucket rotor for 1hr at 1,34,000xg (33,000rpm) for 1hr at 4°C. The brown coloured intact mitochondrial band was observed in between 60% and 32% sucrose interface. Mitochondrial band was carefully removed with cut micropipette tips and was placed it in Beckman centrifuge tube. The tube is filled with SEM buffer (10 mM MOPS/KOH pH 7.2, 250 mM Sucrose, 1 mM EDTA) and spin at 10,000xg for 30mins at 4°C. The resulting purified mitochondrial pellet was suspended in SH buffer (0.6M Sorbitol, 20mM HEPES pH 7.4) at the final concentration of 10mg/ml. Purified mitochondrial aliquots are made and are flash-frozen in liquid nitrogen and stored at −80°C for further use.

### Mitochondrial Sub-fractionation

Mitochondrial sub-fractionation method was adapted from earlier study [51] with some modification as described below and performed. 50µg of purified mitochondria was taken and resuspended in SH (Sorbitol/HEPES) buffer (0.6M Sorbitol, 20mM HEPES pH 7.4). The mitochondria were treated with 0.1% Triton-X 100 at 4°C for 15mins in slow rocking (10 rpm). Spun down the tubes at 17,500Xg for 15mins at 4°C. The isolated insoluble pellet was suspended in SH buffer and mixed with 5X SDS loading dye, boiled the samples and loaded on 10% SDS PAGE. The collected supernatant was precipitated with 12% TCA (Trichloroacetic acid) at −20°C for 15mins. Further the precipitated samples were spun down at 17,500g for 15mins at 4°C. The isolated soluble fraction was suspended in SH buffer and mixed with 5X loading dye, boiled and loaded on 10% SDS-PAGE.

### Protease protection assay

To determine the localization of misfolded proteins in mitochondrial sub-compartments, samples were treated with Proteinase K as described earlier [51, 52]. 50µg of isolated mitochondria was spun down and resuspended in respective isotonic and hypotonic buffers, 400µl of SH and H buffers (20mM HEPES pH 7.4) respectively for 15mins on ice with intermittent tapping. Then, samples were treated with Proteinase K (total 8 µg) and incubated for 15mins on ice and then stopped the reaction with 2mM PMSF (phenyl methyl sulfonyl fluoride) on ice for 15mins. Tubes were spun down at 17,500Xg for 30mins at 4°C. The pellet was resuspended in SH buffer and 5X SDS loading dye. The supernatant after the centrifugation was precipitated with tri-chloroacetic acid (TCA, 6% final). All the samples were boiled at 95°C for 10mins and loaded on 10% SDS-PAGE.

### Nuclear fractionation

The nuclear fractionation protocol was adapted from Hahn lab with minor modification. The overnight grown cultures were re-inoculated in 100ml of YPR media with the initial OD_600_ of 0.5. The cultures were induced for misfolded protein expression with 1% galactose and incubated at 30°C for 12 hrs. Cultures were centrifuged in 50ml polypropylene conical centrifuge tubes and resuspended in 5 ml buffer A (1M Sorbitol, 50mM Tris-HCL pH 7.5,10mM MgCl_2_ +3 mM DTT) and incubated at 30°C for 15mins with slow rocking (120rpm). Culture was pelleted down and resuspended in 5 ml of buffer A and 2 ml YPD/S (YPD+1M Sorbitol). 2.5 mg of Zymolyase 100T (5mg/g of wet weight) was added and spheroplast preparation was done by incubating the suspension at 30°C for 45-60mins with slow rocking (120 rpm) (The spheroplast was checked by suspending it in water and buffer separately and measured, decrease in OD by 1:10 compared with buffered one). Thereafter 10 ml of YPD/S was added, and cells were centrifuged at 4000 rpm for 10 minutes. Cells were resuspended in 25 ml YPD/S (room temp) and again centrifuged at 4000 rpm for 5 minutes at 4°C. Pellet was resuspended in 10 ml of ice-cold buffer N [25mM K2SO_4_, 30mM HEPES pH 7.6, 5mM MgSO_4_, 1mM EDTA, 10% Glycerol, 0.5% NP40, 7.2mM Spermidine Hexahydrate (phosphate salt), 3mM DTT,1x Protease Inhibitors Cocktail] and was homogenized for 15-20 times on ice, depending on the pellet size. The homogenized suspension was centrifuged at 2000rpm at 4°C for 15mins to remove cellular debris. The supernatant was transferred to a new tube and was centrifuged at 6000 rpm for 25 minutes at 4°C. Supernatant was collected and subjected to TCA precipitation, and loaded in SDS-PAGE along with the pellet for western blot

### Measurement of oxygen consumption rate (OCR)

Yeast strains were grown in YPR and the secondary culture was induced at OD_600_ of 0.4-0.6 with 1% galactose and induced for 12 hours at 30°C. OD_600_ are checked after the induction and cells are diluted with media (to OD_600_ of 1.0) and were used for checking the oxygen consumption rate using Oxygraph (Oroboros) instrument. Initially the chambers were loaded with media to get the standard amount of oxygen present in the chamber, then the cells were loaded to measure the oxygen consumption. The uninduced cultures used as a control to measure the difference in oxygen consumption rate of the induced (stressed) cells due to proteotoxic stress.

### Cloning of mitochondrial IMS or matrix targeted PMD for mammalian cell expression

Cloning of OCT-PMD (for specific targeting to mitochondrial matrix) and SMAC-PMD (for specific targeting to mitochondrial IMS) in pEGFP-N1 vector was achieved after amplification of the mitochondrial targeting sequence (MTS, OCT and SMAC) [31] and PMD by overlap PCR. In the first step, amplification of mitochondrial targeting signals and PMD gene sequence were done separately using gene specific set of primers with overhangs for overlap PCR. In the next step touchdown reaction was performed to overlap the PCR amplified products of MTS and PMD. In the final extension step, overlapped fragments were amplified to with the use of end primers (forward primer of MTS and reverse primer of PMD). OCT-PMD or SMAC-PMD sequences were cloned in pEGFPN1 vector in between XhoI and HindIII sites in frame with EGFP.

### Confocal Imaging of mammalian cell (HeLa cells)

Confocal microscopy was performed using Nikon A1R MP+ Ti-E microscope system equipped with solid state lasers. All imaging was performed using Apochromat 100X 1.4 NA objective lens using oil immersion. 488nm and 561nm lasers were used to excite GFP and MitoTracker Red respectively, while 640nm lasers was used to excite far red signals. Prior to imaging cells were transfected with desired construct and incubated in CO_2_ incubator for 16-20 hours.

### Mitochondria Branch Length Measurement

Mitochondrial morphology and branch length were analyzed using Image J plugin MiNA (Mitochondrial Network Analysis) from (https://imagej.net/MiNA_-_Mitochondrial_Network_Analysis)[53]. Initially images were processed with Fiji software in the following order, granular noise were removed by following path; process →Noise→Despeckle, to improve the image quality and mitochondrial visualization local contrast enhancement was performed by following path; Process→ Enhance Local Contrast (CLAHE). After pre-processing, Tubeness plugin of Imagej was used for mitochondrial segmentation and clear visualization of mitochondrial network by following path Plugins→Analyze→ Tubeness. Finally, MiNA plugin was used to calculate mitochondrial mean branch length by following path Plugins→ Stuart Lab→ MiNA Scripts→ MiNA Analyze Morphology. Statistical analysis by the Student’s t-test.

### Cell Cytotoxicity Assay

To perform cell cytotoxicity assay ~ 10,000 HeLa cells were seeded per well in 96 well plate and incubated in CO_2_ incubator for 24 hours. After that transfection with OCT/SMAC-PMD-GFP and control DNA, plate was kept in the incubator for next 48 hours. After 48 hours incubation, cytotoxicity assay was performed using CyQUANT™ LDH Cytotoxicity Assay kit (Invitrogen) following the manufacturer’s protocol.

### RNA isolation and RNA sequencing

Yeast cells were grown and induced with 1% galactose in YPR media for 4hrs, 8hrs, 12hrs or 16hrs and un-induced cells were kept as controls. After growth, cells were harvested at 6000 rpm for 15mins at 4°C. After centrifugation, the cell pellet was washed with chilled 1X PBS; re-suspended in 1 ml of sterile, chilled 1X PBS (pH 7.4) and was transferred to a sterile microfuge tube. Tubes were centrifuged at 12,000 rpm for 5mins at 4°C and the supernatant was discarded carefully. Acid washed glass beads were added to the pellet (two times the volume of the pellet). To this, 1ml Trizol was added to each tube and kept on bead beater for a total of 5-6mins such that it was on the bead beater for 1 min and on ice for the next 2-3mins. Following this, it was spun down on a table-top centrifuge and the supernatant was collected in a fresh centrifuge tube. To the supernatant, chloroform was added (250µl chloroform was added to 800µl of the supernatant) and was vortexed till the solution turned milky pink. This was further incubated at room temperature for 10mins and then centrifuged at 13,000 rpm for 15mins at 4°C. The aqueous phase on top was collected very carefully without disturbing the white layer in the middle, in a fresh centrifuge tube. To this supernatant, 1mL of isopropanol was added, mixed by rotating and kept at −20°C for 2hours. This was then centrifuged at 13,000rpm for 30mins at 4°C. The supernatant was discarded and 750µl of chilled 80% ethanol was used to wash the pellet. The tubes were kept open for air drying the pellet and the pellet was finally re-suspended in 60µl of sterile, nuclease free water. The RNA was visually inspected on ethidium bromide-stained agarose gel to verify the integrity of RNA. Using TruSeq RNA sample prep kit v2 (Stranded mRNA LT kit), the transcriptomic library was prepared according to the manufacturer’s instructions (www.illumina.com). Briefly, 700ng of RNA with high purity and integrity from each sample was used to generate the library. Adaptor-ligated fragments were purified using AMPure XP beads (Agencourt). The library thus generated was run on BioAnalyzer DNA1000 LabChip (Agilent Technologies) to determine the fragment size. The concentration was estimated using Qubit high sensitivity (HS) kit for double stranded DNA on the Qubit instrument (Invitrogen). The average fragment size of the libraries was determined with a BioAnalyzer DNA1000 LabChip (Agilent Technologies). Diluted libraries were multiplex-sequenced and run on a HiSeq 2000 Illumina platform in HiSeq Flow Cell v3 (Illumina Inc., USA) using TruSeq SBS Kit v3 (Illumina Inc., USA) for cluster generation as per manufacturer’s protocol. All transcriptomic were performed with two biological replicates of each sample. From the bcl files that were generated after sequencing, “.fasta” files were created. FastQ was used to check the quality of the reads. Reads with Phred score of 30 or above were taken forward for further analysis. Trimmomatic (v0.43) was used to trim the read sequences, if required. The resulting reads were then aligned to the transcriptome of *S. cerevisiae* strain S288c as available from ENSEMBL. Kallisto was used to perform the alignment and we had a minimum of 80% alignment to the reference. Using Kallisto, we were able to estimate gene expression levels as TPM (transcripts per million) values.

### Transcriptome Data Analysis

The transcriptome data was analyzed with a standard Kallisto pipeline as described previously [54]. Briefly, fastq files were trimmed using Trimmomatic and the fastqc files were aligned to the yeast transcriptome using Kallisto. The TPM values that were obtained were used to analyze the fold changes. Two types of analysis were performed. First, we used a large-scale RNA-seq with 40 samples (uninduced and induced cells of wild type and misfolded protein expressing to specific sub-cellular compartments namely mitochondrial IMS, mitochondrial matrix, ER, cytosol and nucleus). With 40 samples we could verify that the expression patterns of each gene which were normally distributed about their mean over the different conditions. To check if a gene is significantly upregulated or downregulated in a particular sample, we obtained the Z-score value for each gene, with the mean and standard deviation obtained over all the conditions. For each condition, genes that had a Z-score > 2 or <−2 were used to obtain condition-specific upregulated or downregulated gene list, respectively. Each gene shown in the network has p-value less than 0.023 and Z-score greater than 2.

Second, for time course analysis of transcriptome we performed RNA-seq for the strains (IMS-DMMBP, IMS-PMD with IMS-MBP strain as control and MM-DMMBP, MM-PMD with MM-MBP strain as control) at 0hrs, 4 hrs, 8 hrs and 12 hrs post induction of the exogenous proteins. The TPM values were quantile normalized, averaged over replicates and fold change was calculated with respect to 0 hrs TPM value (un-induced set) for each of the compartment-specific targeted exogenous proteins. The fold change values were then used to obtain gene-set specific information using R-package. The fold changes of gene expressions in log2 scale in comparison to 0 time point of each strain have been plotted as heat map. All the fold changes are shown as colormaps with blue indicating down-regulation (scale provided beside each heatmap) and red indicating upregulation (scale provided along each heatmap). In this analysis p-value for each gene is not provided as that would not be a faithful indication of genes that qualify to be upregulated in any other competing condition due to a hard cut-off in p-value (the relative comparison of fold changes show the genes that show contrasting behaviour between IMS and MM stress) and it would not be able to capture the detailed changes that are associated with change in gene expression pattern over time.

### Mass Spectrometry

Yeast strains (Wt, IMS-PMD and MM-PMD) were grown overnight in YPR. Next morning, secondary culture was inoculated at OD_600_ of 0.1 and grown till OD_600_ of 0.5. Each culture was divided in two culture tubes and in one half of each strain was induced with 1% galactose for 12 hours. The other half was grown in YPR as uninduced controls. After induction, equal number of cells were taken from uninduced and induced culture of each strain, cells were harvested by centrifugation and were re-suspended in resuspension buffer (50mM Tris-HCl, pH 7.5, 150 mM NaCl, 1mM EDTA, 5mM MgCl_2_, 1% NP-40, Protease inhibitor cocktail). The re-suspended cells were subjected to lysis by using glass beads in bead beater. Each cycle of bead beating of 3 minutes were followed by incubation in ice for 5 minutes and 3-4 cycles of bead beating were done. After cell lysis, the whole cell lysate was centrifuged at 18,000Xg for 15 minutes and the supernatant were collected in fresh tubes. All lysates were quantified by BCA protein estimation kit (Thermo Scientific) and the concentrations were made equal for all samples (~10mg/ml). The whole cell lysates were boiled at 95°C for 10 minutes with 4X SDS loading buffer [0.2 M Tris–HCl (pH 6.8), 8% SDS, 0.05 M EDTA, 4% 2-mercaptoethanol, 40% glycerol, 0.8% bromophenol blue] and heating at 95 for 10 min and were centrifuged shortly to remove the debris. 150 µg of protein extract from each sample were ran on NuPAGE 4%–12% Bis–Tris Protein Gels (Invitrogen) using MES running buffer (100 mM MES, 100 mM Tris–HCl, 2 mM EDTA, 7 mM SDS) at 200 V for 40 min and fixed and stained with Coomassie brilliant blue. Reduction, alkylation and In-gel trypsin digestion was done as described in Shevchenko et al [55]. Trypsin digested peptides were eluted, desalted and vacuum dried as described in Rappsilber et al [56], and stored in −20 until Mass spectrometry analysis.

Dried peptides were dissolved in 2% formic acid and sonicated in bath sonicator for 5 min. Each sample were loaded in reverse phase liquid chromatography followed by mass spectrometry analysis. Peptides were analysed on Q Exactive (Thermo Scientific) mass spectrometer interfaced with nano-flow LC system (Easy nLC II, Thermo Scientific). EasySpray Nano Column PepMap™ RSLC C18 (Thermo Fisher) (75 μm× 15 cm; 3 μm; 100Å) using 60 min gradient of mobile phase [5% Can containing 0.1% formic acid (buffer A) and 90% ACN (acetonitrile) containing 0.1% formic acid (buffer B)] at flow rate 300nL/min was used for separation of peptide. Full scan of MS spectra (from/z 400 to 1650) were acquired followed by MS/MS scans of top 10peptide with charge state 2 of higher.

Raw files obtained were analysed in MaxQuant Computational platform (Ver. 1.6.10.43) using UniProt database of *Saccharomyces cerevisiae* (released Nov, 2019) [57]. LFQ option was selected for label free quantification with minimum 2 unique peptides for ratio count along with oxidation (M), acetylation (Protein N-term) as variable and carbomethylation as fixed modification. Additional parameters: 2 trypsin missed cleavages, 20 ppm peptide mass tolerance and 1% peptide false discovery rate (FDR) were allowed. Further data analysis and statistical tests were performed in Perseus (Ver. 1.6.0.2) [58]. To compare the control and stressed samples, ratio of IMS-PMD by control and MM-PMD by control is calculated using average LFQ intensities of two biological repeat experiments.

The ratio is converted into Log2 space and mean and standard deviation is calculated for both conditions. Z-score normalization was done for both conditions using the formula

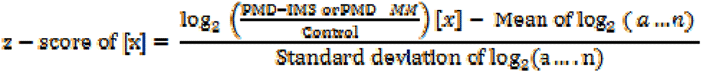

where X is single protein and “a” to “n” is dataset of proteins. Z-score cut-off ±1.96 indicates population lies outside the 95 % interval hence considered significant. The z-score cut-off (1≤ Z) and (−1≥ Z) of proteins is considered as enriched and depleted respectively. Heatmaps were prepared using Morphius (https://software.broadinstitute.org/morpheus).

### Yeast cells viability

To check the viability of yeast cells after proteotoxic stress, cells were grown in YPR and YPR+ 1%-gal media. Cells were harvested after 24 and 48 hours of post-galactose induction. Cells were pelleted down and washed with 1x PBS to wash the media and were resuspended in 1x PBS. As positive control, yeast cells were heated at 100°C for 15 min and for negative control cells were kept unstained. Yeast cells were stained with 5ug\ml propidium iodide for 10 minutes and samples were examined in CytoFlex flow cytometer (Beckman Coulter) using PE channel. A total of 50,000 events were recorded in each experiment, live and dead cell populations were selected from negative and positive control respectively and applied to all test samples.

### Co-immunoprecipitation

Mitochondria from IMS-DMMBP strain were isolated after 4 hrs of induction with 1& galactose. Isolated mitochondria were solubilized with using the solubilization buffer (10mM Tris-HCl pH 7.4,100mM NaCl, 1mM EDTA, 1mM PMSF, protease inhibitor cocktail (Sigma) with 0.5% digitonin. 10% of the solubilized mitochondrial protein extract was used as input. Solubilized mitochondria were bound to Protein A-G Sepharose beads which was already bound to anti-MBP antibody. The resin was washed with the lysis buffer with 0.1% digitonin and bound proteins were eluted by boiling samples at 95°C with SDS sample loading buffer.

### Import Assay

Mitochondria were isolated from yeast cells after 4 hours of induction with the uninduced culture as control. Import of Fluorescein-labelled COXIV signal sequence peptide into isolated mitochondria were carried out in import buffer (0.6M sorbitol, 25mM KCL, 10mM MgCl_2_,2mM KPO_4_,0.5mM EDTA, 2mM ATP, 2mM NADH, 50mM HEPES-KOH pH 7.4 by incubating at 30°C for 10 minutes. 1mM CCCP (Sigma) was used for negative control. Mitochondria was pelleted down by centrifugation at 14000*g for 10 minutes. Mitochondria pellet was washed three times and resuspended in 20µl of Import buffer. 10µl of each sample were applied on glass slides and mounted by a cover slip, then imaging was done with Nikon confocal microscope with 100X oil immersion objective. 488nm laser was used to as excitation source.

