## Supplementary Text for "A comparative study of stress responses elicited by misfolded proteins targeted by bipartite or matrix-targeting signal sequences to yeast mitochondria"

1. **Supplementary References**
2. **Supplementary Table 1**

**Additionally, Supplementary Table S2 (Z-score values across 39 different yeast strains) and Table S3 (TPM values in two replicates for kinetics of gene expression of IMS and matrix strains) contain the raw data of transcriptomics data. Table S4 contains the Z-score values of the proteomics data. Table S5 contains the raw data of GO enrichment analysis by “Yeastmine” of specifically upregulated genes in MM-PMD and IMS-PMD strains (shown in Figure 4).**

1. **Supplementary Figure Legends**

**Figure S1**

**E**. **A.** Confirmation of yeast strains made by homologous recombination of DNA cassette (as shown schematically in Figure 1B) for expression of inducible stressor proteins (as well as folded control proteins) in URA3 locus of parent strain YMJ003. Here, yeast strains containing the cassette for expression of the PMD protein in different subcellular compartments (plating arrangement shown schematically in left panel) are shown in selection media as well as in presence of 5-FOA to confirm the disruption of the URA3 locus. **B.** Expression of different stressor and control proteins targeted to various subcellular compartments shown by western blots. Corresponding yeast strains, after 12 hours of galactose, are subjected to cell lysate preparation followed by western blots. Expression of cyto-MBP and DMMBP along with Wild type (Wt) yeast strain as control (panel i), nuclear-MBP and nuclear-DMMBP in supernatant (following nuclear fraction extraction by Hahn method and TCA precipitation of the supernatant at the last step of protocol (as described in the methodology section) and pellet (panel ii). Western blot with anti-MBP antibody with cell lysates of Wt, IMS-MBP, IMS-DMMBP and MM-MBP and MM-DMMBP strains in uninduced and induced cells (panel iii). Western blot with anti-DHFR antibody with cell lysates of Wt, IMS-DHFR and MM-DHFR strains in uninduced and induced cells (panel iv). Ssc1 western blot is shown as loading control. Two forms of the DHFR protein are visible in precursor (p) and intermediate form (i^m^) and one intermediate form is majorly seen in IMS (i form) which is higher in size than the i^m^ form found in matrix. The without targeting signal form is found in cytosol. **C-F**. Intracellular localization of mitochondria-targeted stressor (or control) proteins by subcellular fraction of mitochondrial and cytosolic fractions. Exclusive enrichment of DMMBP protein in mitochondrial fraction (MF) in IMS-DMMBP (panel C) and MM-DMMBP (panel D) strains are shown by western blot with anti-MBP antibody. As marker for mitochondrial fraction, anti-Porin antibody is used. As marker for cytosolic fraction (CF), anti-PGK1 antibody is used. Minor amount of PGK1 is found in MF but no porin contamination is observed in cytosolic fraction. WCL represents whole cell lysate. Similar fractionation done for MM-PMD-FLAG protein using anti-flag antibody (panel E). Here although majority of protein is enriched in MF, a small fraction of protein is also found in CF indicating problem in mitochondrial import in this strain. Localization of control protein MBP is shown using MM-MBP strain similarly (panel F). **G-H**. Drop-dilution assay of yeast strains expressing the mutC-DHFR (with Wt DHFR as control) (panel F) and DMMBP (with Wt MBP as control) (panel G) proteins in different subcellular compartments are shown in presence and absence of the inducer. ‘ER’ indicates localization stressor (or control) proteins within Endoplasmic Reticulum lumen. ‘Cyto’ indicates localization stressor (or control) proteins in the cytosol. Time mentioned in hours at the upper right corner of the panel indicates the time of incubation of the spotted plates at 30°C before taking pictures.

**Figure S2**

**A.** Secondary structure prediction of the PMD protein. The secondary structures was calculated using PSI-PRED [1] and are marked according to the confidence value at each residue position. The confidence is given with blue bar graphs on top. **B**. 2D binary contact matrix where the X and Y axis denotes the protein sequence of PMD protein. The colours in the heat map correspond to the contact scores, and the range of colours are scaled to blue to red. The plot was generated using DeepMetaPSICOV [2]. **C**. Coomassie brilliant Blue stained gel picture of purified PMD protein after purification by Ni-NTA affinity chromatography followed by gel Filtration chromatography (shown as GFC fraction). **D**. Yeast strain expressing the control protein MBP or wildtype DHFR and stressor protein mutC-DHFR targeted to endoplasmic reticulum (ER) were grown till mid log phase and stressor (or control) proteins were induced with 1% galactose for 4 hours. After induction, GFP fluorescence were measured by flow cytometry. The ratio of mean GFP fluorescence in the induced to uninduced cells was plotted as bar plots for all strains. Error bars represent standard deviation between repeats (n=2). P value was calculated by unpaired Student’s T test (2-tailed). NS indicates non-significant P value. E. Histograms obtained from Propidium Iodide (PI) staining of yeast cells by Flow cytometry are shown. Unstained cells were kept as negative control and yeast cells heated at 100°C were kept as positive control. Different yeast strains (Wt, IMS-DMMBP, IMS-PMD, MM-DMMBP and MM-PMD) were grown in YPR and YPR+1%gal and stained with PI after 24 hrs and 48 hrs of growth. The PI staining profile at both time points were similar and the profile at 48 hrs after growth have been shown. The living and dead cell population were gated into P1 and P2 population, respectively using the fluorescence of the negative and positive control. The same gating into P1 and P2 population were applied to all test samples. Uninduced and induced cells of strains tested show negligible cells in P2 population (less than 2%) indicating no significant cell death in any of the tested strains.

**Figure S3**

**A**. Spot assay of yeast strains as mentioned in the figure. After 16 hours of induction with 1% galactose in YPR broth (indicated as “I” with each strain) with un-induced control (indicated as “U” with each strain), cells were spotted in YPD to stop further expression of the stressor (or control) protein as galactose promoter remains tightly repressed in presence of glucose. Time mentioned on top of each panel indicates time of incubation of plates at 30°C before taking the pictures. **B**. Fluorescence confocal microscopy of yeast cells expressing IMS- or MM-PMD-GFP protein under galactose promoter. Mitochondria-targeted mCherry was expressed from Ura plasmid as a positive control for mitochondria-targeted proteins. **C**. Fluorescence confocal microscopy of yeast cells showing green-fluorescent mitochondria expressing mitochondria targeted yeGFP in the IMS-mutC-DHFR (upper panel), MM-mutC-DHFR (middle panel), and MM-PMD (lower panel) in absence and presence of galactose induction of stressor proteins in respective mitochondrial sub-compartments. As mutC-DHFR is a rapidly degraded misfolded protein, no mitochondrial fragmentation is observed upon expression of this misfolded protein in IMS or matrix. MM-PMD disrupts the tubular network of mitochondria as shown for other stressor protein in Figure 3A. **D**. Bar plot showing percentage viability of HeLa cells after 48 hours of transient transfection of constructs expressing either HA-tagged EGFP or PMD-GFP protein specifically in mitochondrial IMS or matrix. The viability assay was done by LDH cytotoxicity assay kit. For positive control, cells were treated with 500µM H_2_O_2_. Error bars represent standard deviation between repeats (n=3). **E.** Fluorescence confocal microscopy showing uptake of Mitotracker-green probe in MM-PMD (Left panel) and IMS-PMD (Right panel) cells with and without galactose induction.

**Figure S4**

**A.** Network (both genetic and physical) of genes specifically upregulated in IMS-UPR post 16 hours of galactose induction to express PMD protein in IMS. Genes with expression level with Z-score > 2 were taken for making the network in Cytoscape (version 3.8.0) [3]. The size of the nodes indicates closeness centrality in a scale of 0 to 0.46 and the colours indicate betweenness centrality of the nodes in the network in a scale of 0 (blue) to 0.06 (pink). IMS-resident oxidative folding machinery components are shown with black arrows, components of TOM complex are shown with red arrows. **B**. Network (both genetic and physical) of mito-matrix-UPR upregulated genes specifically upregulated in mito-matrix-UPR post 16 hours of galactose induction to express PMD protein in matrix are shown; the genetic network was constructed as described in panel A. Subnetwork of mitochondrial matrix chaperones are highlighted with red dashed box, and a sub-network of cytosolic quality control components is highlighted with blue dashed box. Vms1 has been shown with a red arrow.

**Figure S5**

**A**. Genetic interaction with deletion and depletion strains (*ssc1*d) of IMS and matrix misfolding stress specific strains (with control strains, Wt and IMS-MBP and MM-MBP strains) by drop-dilution assay. “*” and “**” indicate slow growth phenotype of IMS strains (IMS-PMD and IMS-DMMBP) and MM-PMD strains respectively. “*+” and “**+” indicates aggravation of IMS and MM specific phenotypes due to deletion of the genes tested. “*-” and “**-” indicates alleviation of IMS and MM specific phenotypes due to deletion of the genes tested. **B**. Genetic interaction with deletion strains of mitochondrial proteases with IMS and matrix misfolding stress specific strains (with control strains, Wt and IMS-MBP and MM-MBP strains) by drop-dilution assay as described in panel A. C. Genetic interaction with deletion strains of differentially upregulated genes (*COX14* and *COX17*) with IMS and matrix misfolding stress specific strains (with control strains, Wt and IMS-MBP and MM-MBP strains) by drop-dilution assay as described in panel A. **D**. The phenotype of background deletion (or depletion strains) used for the genetic interaction experiment shown in panel A-C are shown by drop-dilution assay on equivalent media and growth condition. All deletion strains shown grow similarly to wild type yeast strain.

**Figure S6**

**A.** In vitro import assay by confocal microscopy of isolated mitochondria using fluorescently tagged COXIV signal peptide as reported previously [4]. Left panel show green-fluorescent mitochondria due to import of fluorescein-tagged COXIV signal peptide isolated from uninduced or induced Wt yeast strain. Lower panel show fluorescence due to treatment of mitochondria with ionophore CCCP. Middle panel: Import assay as shown in panel A for mitochondria isolated from IMS-DMMBP (upper panel) and IMS-PMD (lower panel) strains. -Gal and +Gal panels indicate mitochondria isolated from uninduced and induced strains, respectively. Right panel: Similar import assay shown in panel A and B with mitochondria isolated from MM-DMMBP (upper panel) and MM-PMD (lower panel) strains. **B.** Co-immunoprecipitation of Tom40 protein with IMS-targeted DMMBP protein. Mitochondria isolated after 4 hours of galactose induction from IMS-DMMBP strain was subjected to co-immunoprecipitation using anti-MBP antibody. PI indicates pre-immune sera as control. 10% of solubilized mitochondrial proteins loaded to both PI sera and anti-MBP antibody bound Protein-AG-Sepharose beads are shown as load. **C**. Gene expression changes of components of cytosolic heat shock response pathways of yeast strains with IMS and matrix (MM) misfolding stress due to overexpression of stressor proteins post 4 hrs, 8 hrs and 12 hrs of galactose induction are shown as heatmap. Wt MBP has been kept as control of overexpression of a folded protein in the respective compartments. The fold changes of gene expressions in log2 scale in comparison to 0 time point of each strain have been plotted as heat map. **D**. Heatmap of z-scores depicting protein-abundance of cellular chaperones and proteins known to bind to unfolded protein, after 12 hours of expression of PMD protein in matrix (MM) or IMS (n=2). **E**. Gene expression changes of components of proteasome of yeast strains with IMS and matrix (MM) misfolding stress due to overexpression of stressor proteins post 4 hrs, 8 hrs and 12 hrs of galactose induction are shown as heatmap as described in panel A.

**Figure S7**

**A-B.** Scatter plot of protein abundance (panel A) distribution of mitochondrial respiratory chain complexes after 12 hours of expression of PMD protein in IMS and matrix. Panel B: Distribution of glycolysis enzymes in the same strains (n=2). **C**. Drop-dilution assay of MM-PMD strain in fermentable (YPR, Yeast extract-Peptone-Raffinose) and non-fermentable (YPGly, Yeast extract-Peptone-Glycerol) media in absence and presence of inducer (1% galactose) are shown. “**” indicates spots with visible growth phenotype in case of MM-PMD strains and “**+” indicates aggravation of growth phenotype of MM-PMD strain in non-fermentable media after induction of misfolding stress. The time mentioned in hours at the upper right corner of the panel indicates the time of incubation of the spotted plates at 30°C before taking pictures. **D.** Representative kinetic traces of Oxygen consumption rate (OCR) of wild type (Wt) (upper panel), IMS-PMD (middle panel) and MM-PMD (lower panel) yeast strains with and without inducer (1% galactose) and in presence and absence of ascorbate is plotted. **E.** Drop-dilution assay as of yeast strains containing mitochondrial IMS and matrix targeted stressor proteins DMMBP and PMD (with wt MBP as control) in presence of inducer and inducer plus ascorbate. “*” and “**” indicates spots with visible growth phenotype in case of IMS-PMD or IMS-DMMBP and MM-PMD strains respectively as shown in Figure 1F. “*+” and “**+” indicates spots with further aggravation of growth phenotype in presence of ascorbate in IMS-PMD/IMS-DMMBP and MM-PMD respectively. Time mentioned in hours at the upper right corner of the panel indicates the time of incubation of the spotted plates at 30°C before taking pictures. **F.** Gene expression changes of components of TCA cycle with IMS and matrix (MM) misfolding stress due to overexpression of stressor proteins post 4 hrs, 8 hrs and 12 hrs of galactose induction are shown as heatmap as described in Figure 4A. **G**. Heatmap of z-scores depicting protein-abundance of components of TCA cycle enzymes, after 12 hours of expression of PMD protein in matrix (MM) or IMS (n=2).

**Figure S8**

**A**. Drop-dilution assay as of yeast strains containing deletion of *MSP1* (upper panel) or *UBX2* (upper panel) expressing mitochondrial matrix targeted stressor protein PMD in presence and absence of the inducer. “**” indicates spots with visible growth phenotype in case of MM-PMD or MM-PMD with deletion of *MSP1* or *UBX2*which remained unchanged due to deletion of these components which show further growth phenotype aggravation in IMS-PMD strains (Figure 5C and D). Spots are done in 1:10 serial dilutions and time mentioned in hours at the upper right corner of the panel indicates the time of incubation of the spotted plates at 30°C before taking pictures. **B-C**. Drop-dilution assay of IMS-PMD-*tom6Δ* (panel B) and IMS-PMD-*tom7Δ* (panel C) strains transformed with plasmids overexpressing (OE) individual components of TOM complex (*TOM6, TOM22, TOM40* or *TOM70*) was performed by spotting the along with wild type yeast strain and same strains with empty vector (EV) as controls on the (SR-Ura) plates (Synthetic media without uracil with 2% Raffinose), with inducer plate (SR-Ura+gal). “*-” indicates spots with alleviated growth phenotype of IMS-PMD-*tom6Δ* denoted as “*” due to overexpression of individual TOM complex components. **D**. Fluorescence confocal microscopy of MM-MBP (folded protein control) and MM-PMD yeast strains transformed with Vms1-GFP plasmid and co-stained with MitoTracker Red dye, after induction with 1% galactose. Images of MM-MBP induced cells (left panel) is showing red tubular mitochondrial network and diffused expression of GFP-tagged Vms1 protein in the cytosol. White boxed region is shown in zoomed in form on the left side to show the distinction between tubular red fluorescence of mitochondria-resident MitoTracker-Red dye and diffused green fluorescence of Vms1-GFP. Images of MM-PMD induced cells (right panel) is showing much less staining of mitochondria and fragmented mitochondrial network due to proteotoxic stress in matrix. In the stressed cells also, expression of GFP-tagged-Vms1 protein is mainly observed in the cytosol. **E**. Oxygen consumption rate (OCR) of MM-PMD and MM-PMD –*vms1*Δ strains with and without misfolding stress, is plotted.

2. **Supplementary References**

[1] Buchan DWA, Jones DT. The PSIPRED Protein Analysis Workbench: 20 years on. Nucleic Acids Res. 2019;47:W402-W7.

[2] Kandathil SM, Greener JG, Jones DT. Recent developments in deep learning applied to protein structure prediction. Proteins. 2019;87:1179-89.

[3] Shannon P, Markiel A, Ozier O, Baliga NS, Wang JT, Ramage D, et al. Cytoscape: a software environment for integrated models of biomolecular interaction networks. Genome Res. 2003;13:2498-504.

[4] Martinez-Caballero S, Peixoto PM, Kinnally KW, Campo ML. A fluorescence assay for peptide translocation into mitochondria. Anal Biochem. 2007;362:76-82.

3. **Supplementary Table 1**

Supplementary Table 1 contains the List of yeast strains, plasmids, primers and Antibodies and other reagents used for this study

**List of plasmids used for this study**

| **S.No** | **Plasmid** | **Alias** | **Description** | **Reference/ Source** |
| --- | --- | --- | --- | --- |
| **1** | **pKM1** | pYMN23 | **pAgTEF-NAT-tScADH1,pGAL1-Cyc1t** | **EUROSCARF** |
| **2** | **pKM11** | pYMN23-ERss | **NAT-Gal1p-ERss-Cyc1t** | This study |
| **3** | **pKM12** | pYMN23-MMss | **NAT-Gal1p-MMss-Cyc1** | This study |
| **4** | **pKM13** | pYMN23-NUCLss | **NAT-Galp-NUCLss-Cyc1** | This study |
| **5** | **pKM14** | pYMN23-CYTOss | **NAT-Gal1p-CYTOss-Cyc1** | This study |
| **6** | **pKM15** | pYMN23-IMSss | **NAT-Gal1pIMSss-Cyc1** | This study |
| **7** | **pBN101** | pYMN23-ERss-PMD | **NAT-Gal1p-ERss-PMD-Cyc1** | This study |
| **8** | **pBN102** | pYMN23-MMss-PMD | **NAT-Gal1p-MMss-PMD-Cyc1** | This study |
| **9** | **pBN103** | pYMN23-NUCLss-PMD | **NAT-Gal1p-NUCLss-PMD-Cyc1** | This study |
| **10** | **pBN104** | pYMN23-CYTOss-pMD | **NAT-Gal1p-CYTO-PMD-Cyc1** | This study |
| **11** | **pBN105** | pYMN23-IMSss-PMD | **NAT-Gal1p-IMSss-PMD-Cyc1** | This study |
| **12** | **pBN201** | pYMN23-ERss-wtDHFR | **NAT-Gal1p-ERss-wtDHFR-Cyc1** | This study |
| **13** | **pBN202** | pYMN23-MMss-wtDHFR | **NAT-Gal1p-MMss-wtDHFR-Cyc1** | This study |
| **14** | **pBN203** | pYMN23-NUCLss-wtDHFR | **NAT-Gal1p-NUCLss-wtDHFR-Cyc1** | This study |
| **15** | **pBN204** | pYMN23-CYTOss-wtDHFR | **NAT-Gal1p-CYTOss-wtDHFR-Cyc1** | This study |
| **16** | **pBN205** | pYMN23-IMSss-wtDHFR | **NAT-Gal1p-IMSss-wtDHFR-Cyc1** | This study |
| **17** | **pBN301** | pYMN23-ERss-mutCDHFR | **NAT-Gal1p-ERss-mutCDHFR-Cyc1** | This study |
| **18** | **pBN302** | pYMN23-MMss-mutCDHFR | **NAT-Gal1p-MMss-mutCDHFR-Cyc1** | This study |
| **19** | **pBN303** | pYMN23-NUCLss-mutCDHFR | **NAT-Gal1p-NUCLss-mutCDHFR-Cyc1** | This study |
| **20** | **pBN304** | pYMN23-CYTOss-mutCDHFR | **NAT-Gal1p-CYTOss-mutCDHFR-Cyc1** | This study |
| **21** | **pBN305** | pYMN23-IMSss-mutCDHFR | **NAT-Gal1p-IMSss-mutCDHFR-Cyc1** | This study |
| **22** | **pPR401** | pYMN23-ERss-MBP | **NAT-Gal1p-ERss-MBP-Cyc1** | This study |
| **23** | **pPR402** | pYMN23-MMss-MBP | **NAT-Gal1p-MMss-MBP-Cyc1** | This study |
| **24** | **pPR403** | pYMN23-NUCLss-MBP | **NAT-Gal1p-NUCLss-MBP-Cyc1** | This study |
| **25** | **pPR404** | pYMN23-CYTOss-MBP | **NAT-Gal1p-CYTOss-MBP-Cyc1** | This study |
| **26** | **pPR405** | pYMN23-IMSss-MBP | **NAT-Gal1p-IMSss-DMMBP-Cyc1** | This study |
| **27** | **pPR501** | pYMN23-ERss-DMMBP | **NAT-Gal1p-ERss-MBP-Cyc1** | This study |
| **28** | **pPR502** | pYMN23-MMss-DMMBP | **NAT-Gal1p-MMss- DMMBP -Cyc1** | This study |
| **29** | **pPR503** | pYMN23-NUCLss-DMMBP | **NAT-Gal1p-NUCLss- DMMBP -Cyc1** | This study |
| **30** | **pPR504** | pYMN23-CYTOss-DMMBP | **NAT-Gal1p-CYTOss- DMMBP -Cyc1** | This study |
| **31** | **pPR505** | pYMN23-IMSss-DMMBP | **NAT-Gal1p-IMSss- DMMBP -Cyc1** | This study |
| **32** | **pBN112** | pYMN23-MMss-PMD-GFP | **NAT-Gal1p-MMss-PMD-GFP-Cyc1** | This study |
| **33** | **pBN115** | pYMN23-IMSss-PMD-GFP | **NAT-Gal1p-IMSss-PMD-GFP-Cyc1** | This study |
| **34** | **pBN122** | pYMN23-MMss-PMD-Flag | **NAT-Gal1p-MMss-PMD-Flag-Cyc1** | This study |
| **35** | **pRS316i** | pRS316-CUP1p-**IMSss-mCherry**-CYCt-URA-CEN | **IMS targeting Cherry** | This study |
| **36** | **pRS316m** | pRS316-CUP1p-**MMss-mCherry**-CYCt-URA-CEN | **MM targeting mCherry** | This study |
| **37** | **pVT100U-mtGFP** | pVT100U-mtGFP | **MM targeting GFP** |  |
| **38** | **pRS313** | pRS313-HIS-CEN | **Empty vector** |  |
| **39** | **pRS315** | pRS315-LEU-CEN | **Empty vector** |  |
| **40** | **pRS316** | pRS316-URA-CEN | **Empty vector** |  |
| **41** | **P426-GPD** | P426-GPDp-CYC1t-URA-2u | **Empty vector** |  |
| **42** | **pJH10c** | mutC-DHFR | **T39A, E173D mutation in DHFR** | NKC Gowda et al., PNAS2013 |
| **43** | **pJR8252A** | pRS416-VMS1-6HIS-2HA-URA-CEN | **Full length VMS1** | Heo JM et al., Mol Biol Cell 2013 |
| **44** | **pJR3462D** | pRS416-VMS1-GFP-URA-CEN | **VMS1-GFP** | Heo JM et al., Mol Biol Cell 2013 |
| **45** | **pJR6107B** | pRS416-ΔN-VMS1-GFP-URA-CEN | **ΔN-VMS1-GFP** | Heo JM et al., Mol Biol Cell 2013 |
| **46** | **pJR9532D** | pRS416-VMS1(1-417) -URA-CEN | **ΔC-VMS1-GFP** | Heo JM et al., Mol Biol Cell 2013 |
| **47** | **pETDuet-1-PMD** | pETDuet-PMD | **PMD** | This study |

**List of yeast strains used for this study**

| **S.No** | **Strain Name** | **Genotype** | **Common name** | **Parent strain** | **Reference/ Source** |
| --- | --- | --- | --- | --- | --- |
| **1** | **BY4741** | MATa *his3Δ1 leu2Δ0 met15Δ0 ura3Δ0* | **WT** | S288C | S288C/ EUROSCARF |
| **2** | **BY4742** | *MATα* ; his3Δ 1; leu2Δ 0; lys2Δ 0; ura3Δ 0 | **WT** | S288C | S288C/EUROSCARF |
| **3** | **yMJ003** | *MATα* his3Δ1 leu2Δ0 met15Δ0 ura3Δ0 LYS+Δcan1::STE2pr-spHIS5 Δlyp1::STE3pr-LEU2 cyh2 Δura3::**UPRE-GFP-TEF2pr-RFP-MET15-URA3** | **UPR reporter strain** | S288C | Jonikas et al., 2009 |
| **4** | **KBN101** | *MATα* his3Δ1 leu2Δ0 met15Δ0 ura3Δ0 LYS+Δcan1::STE2pr-spHIS5 Δlyp1::STE3pr-LEU2 cyh2 Δura3::UPRE-GFP-TEF2pr-RFP-MET15-**URA3::NAT-ER-Wt-DHFR** | **ER-wtDHFR** | yMJ003 | This study |
| **5** | **KBN102** | *MATα* his3Δ1 leu2Δ0 met15Δ0 ura3Δ0 LYS+Δcan1::STE2pr-spHIS5 Δlyp1::STE3pr-LEU2 cyh2 Δura3::UPRE-GFP-TEF2pr-RFP-MET15-**URA3::NAT-MM-wtDHFR** | **MM-WtDHFR** | yMJ003 | This study |
| **6** | **KBN103** | *MATα* his3Δ1 leu2Δ0 met15Δ0 ura3Δ0 LYS+Δcan1::STE2pr-spHIS5 Δlyp1::STE3pr-LEU2 cyh2 Δura3::UPRE-GFP-TEF2pr-RFP-MET15-**URA3::NAT-Nucl-wtDHFR** | **Nucl-wtDHFR** | yMJ003 | This study |
| **7** | **KBN104** | *MATα* his3Δ1 leu2Δ0 met15Δ0 ura3Δ0 LYS+Δcan1::STE2pr-spHIS5 Δlyp1::STE3pr-LEU2 cyh2 Δura3::UPRE-GFP-TEF2pr-RFP-MET15-**URA3::NAT-Cyto-wtDHFR** | **Cyto-wtDHFR** | yMJ003 | This study |
| **8** | **KBN105** | *MATα* his3Δ1 leu2Δ0 met15Δ0 ura3Δ0 LYS+Δcan1::STE2pr-spHIS5 Δlyp1::STE3pr-LEU2 cyh2 Δura3::UPRE-GFP-TEF2pr-RFP-MET15-**URA3::NAT-IMS-wtDHFR** | **IMS-WtDHFR** | yMJ003 | This study |
| **9** | **KBN201** | *MATα* his3Δ1 leu2Δ0 met15Δ0 ura3Δ0 LYS+Δcan1::STE2pr-spHIS5 Δlyp1::STE3pr-LEU2 cyh2 Δura3::UPRE-GFP-TEF2pr-RFP-MET15-**URA3::NAT-ER-mutCDHFR** | **ER-mutCDHFR** | yMJ003 | This study |
| **10** | **KBN202** | *MATα* his3Δ1 leu2Δ0 met15Δ0 ura3Δ0 LYS+Δcan1::STE2pr-spHIS5 Δlyp1::STE3pr-LEU2 cyh2 Δura3::UPRE-GFP-TEF2pr-RFP-MET15-**URA3::NAT-MM-mutCDHFR** | **MM-mutCDHFR** | yMJ003 | This study |
| **11** | **KBN203** | *MATα* his3Δ1 leu2Δ0 met15Δ0 ura3Δ0 LYS+Δcan1::STE2pr-spHIS5 Δlyp1::STE3pr-LEU2 cyh2 Δura3::UPRE-GFP-TEF2pr-RFP-MET15-**URA3::NAT-Nucl-mutCDHFR** | **Nucl-mutCDHFR** | yMJ003 | This study |
| **12** | **KBN204** | *MATα* his3Δ1 leu2Δ0 met15Δ0 ura3Δ0 LYS+Δcan1::STE2pr-spHIS5 Δlyp1::STE3pr-LEU2 cyh2 Δura3::UPRE-GFP-TEF2pr-RFP-MET15-**URA3::NAT-Cyto-mutCDHFR** | **Cyto-mutCDHFR** | yMJ003 | This study |
| **13** | **KBN205** | *MATα* his3Δ1 leu2Δ0 met15Δ0 ura3Δ0 LYS+Δcan1::STE2pr-spHIS5 Δlyp1::STE3pr-LEU2 cyh2 Δura3::UPRE-GFP-TEF2pr-RFP-MET15-**URA3::NAT-IMS-mutCDHFR** | **IMS-mutcDHFR** | yMJ003 | This study |
| **14** | **KBN301** | *MATα* his3Δ1 leu2Δ0 met15Δ0 ura3Δ0 LYS+Δcan1::STE2pr-spHIS5 Δlyp1::STE3pr-LEU2 cyh2 Δura3::UPRE-GFP-TEF2pr-RFP-MET15-**URA3::NAT-ER-PMD** | **ER-PMD** | yMJ003 | This study |
| **15** | **KBN302** | *MATα* his3Δ1 leu2Δ0 met15Δ0 ura3Δ0 LYS+Δcan1::STE2pr-spHIS5 Δlyp1::STE3pr-LEU2 cyh2 Δura3::UPRE-GFP-TEF2pr-RFP-MET15-**URA3::NAT-MM-PMD** | **MM-PMD** | yMJ003 | This study |
| **16** | **KBN303** | *MATα* his3Δ1 leu2Δ0 met15Δ0 ura3Δ0 LYS+Δcan1::STE2pr-spHIS5 Δlyp1::STE3pr-LEU2 cyh2 Δura3::UPRE-GFP-TEF2pr-RFP-MET15-**URA3::NAT-Nucl-PMD** | **Nucl-PMD** | yMJ003 | This study |
| **17** | **KBN304** | *MATα* his3Δ1 leu2Δ0 met15Δ0 ura3Δ0 LYS+Δcan1::STE2pr-spHIS5 Δlyp1::STE3pr-LEU2 cyh2 Δura3::UPRE-GFP-TEF2pr-RFP-MET15-**URA3::NAT-Cyto-PMD** | **Cyto-PMD** | yMJ003 | This study |
| **18** | **KBN305** | *MATα* his3Δ1 leu2Δ0 met15Δ0 ura3Δ0 LYS+Δcan1::STE2pr-spHIS5 Δlyp1::STE3pr-LEU2 cyh2 Δura3::UPRE-GFP-TEF2pr-RFP-MET15-**URA3::NAT-IMS-PMD** | **IMS-PMD** | yMJ003 | This study |
| **19** | **PP401** | *MATα* his3Δ1 leu2Δ0 met15Δ0 ura3Δ0 LYS+Δcan1::STE2pr-spHIS5 Δlyp1::STE3pr-LEU2 cyh2 Δura3::UPRE-GFP-TEF2pr-RFP-MET15-**URA3::NAT-ER-MBP** | **ER-MBP** | yMJ003 | This study |
| **20** | **PP402** | *MATα* his3Δ1 leu2Δ0 met15Δ0 ura3Δ0 LYS+Δcan1::STE2pr-spHIS5 Δlyp1::STE3pr-LEU2 cyh2 Δura3::UPRE-GFP-TEF2pr-RFP-MET15-**URA3::NAT-MM-MBP** | **MM-MBP** | yMJ003 | This study |
| **21** | **PP403** | *MATα* his3Δ1 leu2Δ0 met15Δ0 ura3Δ0 LYS+Δcan1::STE2pr-spHIS5 Δlyp1::STE3pr-LEU2 cyh2 Δura3::UPRE-GFP-TEF2pr-RFP-MET15-**URA3::NAT-Nucl-MBP** | **Nucl-MBP** | yMJ003 | This study |
| **22** | **PP404** | *MATα* his3Δ1 leu2Δ0 met15Δ0 ura3Δ0 LYS+Δcan1::STE2pr-spHIS5 Δlyp1::STE3pr-LEU2 cyh2 Δura3::UPRE-GFP-TEF2pr-RFP-MET15-**URA3::NAT-Cyto-MBP** | **Cyto-MBP** | yMJ003 | This study |
| **23** | **PP405** | *MATα* his3Δ1 leu2Δ0 met15Δ0 ura3Δ0 LYS+Δcan1::STE2pr-spHIS5 Δlyp1::STE3pr-LEU2 cyh2 Δura3::UPRE-GFP-TEF2pr-RFP-MET15-**URA3::NAT-IMS-MBP** | **IMS-MBP** | yMJ003 | This study |
| **24** | **PP501** | *MATα* his3Δ1 leu2Δ0 met15Δ0 ura3Δ0 LYS+Δcan1::STE2pr-spHIS5 Δlyp1::STE3pr-LEU2 cyh2 Δura3::UPRE-GFP-TEF2pr-RFP-MET15-**URA3::NAT-ER-DMMBP** | **ER-DMMBP** | yMJ003 | This study |
| **25** | **PP502** | *MATα* his3Δ1 leu2Δ0 met15Δ0 ura3Δ0 LYS+Δcan1::STE2pr-spHIS5 Δlyp1::STE3pr-LEU2 cyh2 Δura3::UPRE-GFP-TEF2pr-RFP-MET15-**URA3::NAT-MM-DMMBP** | **MM-DMMBP** | yMJ003 | This study |
| **26** | **PP503** | *MATα* his3Δ1 leu2Δ0 met15Δ0 ura3Δ0 LYS+Δcan1::STE2pr-spHIS5 Δlyp1::STE3pr-LEU2 cyh2 Δura3::UPRE-GFP-TEF2pr-RFP-MET15-**URA3::NAT-Nucl-DMMBP** | **Nucl-DMMBP** | yMJ003 | This study |
| **27** | **PP504** | *MATα* his3Δ1 leu2Δ0 met15Δ0 ura3Δ0 LYS+Δcan1::STE2pr-spHIS5 Δlyp1::STE3pr-LEU2 cyh2 Δura3::UPRE-GFP-TEF2pr-RFP-MET15-**URA3::NAT-Cyto-DMMBP** | **Cyto-DMMBP** | yMJ003 | This study |
| **28** | **PP505** | *MATα* his3Δ1 leu2Δ0 met15Δ0 ura3Δ0 LYS+Δcan1::STE2pr-spHIS5 Δlyp1::STE3pr-LEU2 cyh2 Δura3::UPRE-GFP-TEF2pr-RFP-MET15-**URA3::NAT-IMS-DMMBP** | **IMS-DMMBP** | yMJ003 | This study |
| **29** | **KBN312** | *MATα* his3Δ1 leu2Δ0 met15Δ0 ura3Δ0 LYS+Δcan1::STE2pr-spHIS5 Δlyp1::STE3pr-LEU2 cyh2 Δura3::UPRE-GFP-TEF2pr-RFP::KanMX -MET15 -**URA3::NAT-MM-PMD-GFP** | **MM-PMD-GFP** | KBN302 | This study |
| **30** | **KBN315** | *MATα* his3Δ1 leu2Δ0 met15Δ0 ura3Δ0 LYS+Δcan1::STE2pr-spHIS5 Δlyp1::STE3pr-LEU2 cyh2 Δura3::UPRE-GFP-TEF2pr-RFP::KanMX--MET15-**URA3::NAT-IMS-PMD-GFP** | **IMS-PMD-GFP** | KBN305 | This study |
| **31** | **KBN322** | *MATα* his3Δ1 leu2Δ0 met15Δ0 ura3Δ0 LYS+Δcan1::STE2pr-spHIS5 Δlyp1::STE3pr-LEU2 cyh2 Δura3::UPRE-GFP-TEF2pr-RFP-MET15-**URA3::NAT-MM-PMD+pVT100U-mtGFP** | **MM-PMD-pVT100U-mtGFP** | KBN302 | This study |
| **32** | **KBN325** | *MATα* his3Δ1 leu2Δ0 met15Δ0 ura3Δ0 LYS+Δcan1::STE2pr-spHIS5 Δlyp1::STE3pr-LEU2 cyh2 Δura3::UPRE-GFP-TEF2pr-RFP-MET15-**URA3::NAT-IMS-PMD+pVT100U-mtGFP** | **IMS-PMD -pVT100U-mtGFP** | KBN305 | This study |
| **33** | **PP522** | *MATα* his3Δ1 leu2Δ0 met15Δ0 ura3Δ0 LYS+Δcan1::STE2pr-spHIS5 Δlyp1::STE3pr-LEU2 cyh2 Δura3::UPRE-GFP-TEF2pr-RFP-MET15-**URA3::NAT-MM-DMMBP+pVT100U-mtGFP** | **MM-DMMBP-pVT100U-mtGFP** | PP502 | This study |
| **34** | **PP525** | *MATα* his3Δ1 leu2Δ0 met15Δ0 ura3Δ0 LYS+Δcan1::STE2pr-spHIS5 Δlyp1::STE3pr-LEU2 cyh2 Δura3::UPRE-GFP-TEF2pr-RFP-MET15-**URA3::NAT-IMS-DMMBP+pVT100U-mtGFP** | **IMS-DMMBP-pVT100U-mtGFP** | PP505 | This study |
| **35** | **KBN332** | *MATα* his3Δ1 leu2Δ0 met15Δ0 ura3Δ0 LYS+Δcan1::STE2pr-spHIS5 Δlyp1::STE3pr-LEU2 cyh2 Δura3::UPRE-GFP-TEF2pr-RFP-MET15-**URA3::NAT-MM-PMD+ pRS426-GPD-VMS1-Flag** | **MM-PMD- pRS426-GPD-VMS1-Flag_OE** | KBN302 | This study |
| **36** | **KBN342** | *MATα* his3Δ1 leu2Δ0 met15Δ0 ura3Δ0 LYS+Δcan1::STE2pr-spHIS5 Δlyp1::STE3pr-LEU2 cyh2 Δura3::UPRE-GFP-TEF2pr-RFP-MET15-**URA3::NAT-MM-PMD+ pJR8252A** | **MM-PMD- pRS416-VMS1-6HIS-2HA** | KBN302 | This study |
| **37** | **KBN352** | *MATα* his3Δ1 leu2Δ0 met15Δ0 ura3Δ0 LYS+Δcan1::STE2pr-spHIS5 Δlyp1::STE3pr-LEU2 cyh2 Δura3::UPRE-GFP-TEF2pr-RFP-MET15-**URA3::NAT-MM-PMD+ pJR3462D** | **MM-PMD- pRS416-VMS1-GFP** | KBN302 | This study |
| **38** | **KBN362** | *MATα* his3Δ1 leu2Δ0 met15Δ0 ura3Δ0 LYS+Δcan1::STE2pr-spHIS5 Δlyp1::STE3pr-LEU2 cyh2 Δura3::UPRE-GFP-TEF2pr-RFP-MET15-**URA3::NAT-MM-PMD+ pJR6107B** | **MM-PMD- pRS416-ΔN-VMS1-GFP** | KBN302 | This study |
| **39** | **KBN372** | *MATα* his3Δ1 leu2Δ0 met15Δ0 ura3Δ0 LYS+Δcan1::STE2pr-spHIS5 Δlyp1::STE3pr-LEU2 cyh2 Δura3::UPRE-GFP-TEF2pr-RFP-MET15-**URA3::NAT-MM-PMD+ pJR9532D** | **MM-PMD- pRS416-VMS1(1-417)** | KBN302 | This study |
| **40** | **KBN382** | *MATα* his3Δ1 leu2Δ0 met15Δ0 ura3Δ0 LYS+Δcan1::STE2pr-spHIS5 Δlyp1::STE3pr-LEU2 cyh2 Δura3::UPRE-GFP-TEF2pr-RFP-MET15-**URA3::NAT-MM-PMD+ pRS426-GPD-ΔN-VMS1-GFP** | **MM-PMD- pRS426-GPD-ΔN-VMS1-GFP** | KBN302 | This study |
| **41** | **KBN335** | *MATα* his3Δ1 leu2Δ0 met15Δ0 ura3Δ0 LYS+Δcan1::STE2pr-spHIS5 Δlyp1::STE3pr-LEU2 cyh2 Δura3::UPRE-GFP-TEF2pr-RFP-MET15-**URA3::NAT-IMS-PMD+ pRS426-GPD-VMS1-Flag** | **IMS-PMD -pRS426-GPD-VMS1-Flag_OE** | KBN305 | This study |
| **42** | **KBN345** | *MATα* his3Δ1 leu2Δ0 met15Δ0 ura3Δ0 LYS+Δcan1::STE2pr-spHIS5 Δlyp1::STE3pr-LEU2 cyh2 Δura3::UPRE-GFP-TEF2pr-RFP-MET15-**URA3::NAT-IMS-PMD+ pJR8252A** | **IMS-PMD - pRS416-VMS1-6HIS-2HA** | KBN305 | This study |
| **43** | **KBN355** | *MATα* his3Δ1 leu2Δ0 met15Δ0 ura3Δ0 LYS+Δcan1::STE2pr-spHIS5 Δlyp1::STE3pr-LEU2 cyh2 Δura3::UPRE-GFP-TEF2pr-RFP-MET15-**URA3::NAT-IMS-PMD+ pJR3462D** | **IMS-PMD - pRS416-VMS1-GFP** | KBN305 | This study |
| **44** | **KBN365** | *MATα* his3Δ1 leu2Δ0 met15Δ0 ura3Δ0 LYS+Δcan1::STE2pr-spHIS5 Δlyp1::STE3pr-LEU2 cyh2 Δura3::UPRE-GFP-TEF2pr-RFP-MET15-**URA3::NAT-IMS-PMD+ pJR6107B** | **IMS-PMD - pRS416-ΔN-VMS1-GFP** | KBN305 | This study |
| **45** | **KBN375** | *MATα* his3Δ1 leu2Δ0 met15Δ0 ura3Δ0 LYS+Δcan1::STE2pr-spHIS5 Δlyp1::STE3pr-LEU2 cyh2 Δura3::UPRE-GFP-TEF2pr-RFP-MET15-**URA3::NAT-IMS-PMD+ pJR9532D** | **IMS-PMD - pRS416-VMS1(1-417)** | KBN305 | This study |
| **46** | **KBN385** | *MATα* his3Δ1 leu2Δ0 met15Δ0 ura3Δ0 LYS+Δcan1::STE2pr-spHIS5 Δlyp1::STE3pr-LEU2 cyh2 Δura3::UPRE-GFP-TEF2pr-RFP-MET15-**URA3::NAT-IMS-PMD+ pRS426-GPD-ΔN-VMS1-GFP** | **IMS-PMD - pRS426-GPD-ΔN-VMS1-GFP_OE** | KBN305 | This study |
| **47** | **PP432** | *MATα* his3Δ1 leu2Δ0 met15Δ0 ura3Δ0 LYS+Δcan1::STE2pr-spHIS5 Δlyp1::STE3pr-LEU2 cyh2 Δura3::UPRE-GFP-TEF2pr-RFP-MET15-**URA3::NAT-MM-MBP+ pRS426-GPD-VMS1-Flag** | **MM-DMMBP- pRS426-GPD-VMS1-Flag_OE** | PP402 | This study |
| **48** | **PP442** | *MATα* his3Δ1 leu2Δ0 met15Δ0 ura3Δ0 LYS+Δcan1::STE2pr-spHIS5 Δlyp1::STE3pr-LEU2 cyh2 Δura3::UPRE-GFP-TEF2pr-RFP-MET15-**URA3::NAT-MM-MBP+ pJR8252A** | **MM-DMMBP- pRS416-VMS1-6HIS-2HA** | PP402 | This study |
| **49** | **PP452** | *MATα* his3Δ1 leu2Δ0 met15Δ0 ura3Δ0 LYS+Δcan1::STE2pr-spHIS5 Δlyp1::STE3pr-LEU2 cyh2 Δura3::UPRE-GFP-TEF2pr-RFP-MET15-**URA3::NAT-MM-MBP+ pJR3462D** | **MM-DMMBP- pRS416-VMS1-GFP** | PP402 | This study |
| **50** | **PP462** | *MATα* his3Δ1 leu2Δ0 met15Δ0 ura3Δ0 LYS+Δcan1::STE2pr-spHIS5 Δlyp1::STE3pr-LEU2 cyh2 Δura3::UPRE-GFP-TEF2pr-RFP-MET15-**URA3::NAT-MM-MBP+ pJR6107B** | **MM-DMMBP- pRS416-ΔN-VMS1-GFP** | PP402 | This study |
| **51** | **PP472** | *MATα* his3Δ1 leu2Δ0 met15Δ0 ura3Δ0 LYS+Δcan1::STE2pr-spHIS5 Δlyp1::STE3pr-LEU2 cyh2 Δura3::UPRE-GFP-TEF2pr-RFP-MET15-**URA3::NAT-MM-MBP+ pJR9532D** | **MM-DMMBP- pRS416-VMS1(1-417)** | PP402 | This study |
| **52** | **PP482** | *MATα* his3Δ1 leu2Δ0 met15Δ0 ura3Δ0 LYS+Δcan1::STE2pr-spHIS5 Δlyp1::STE3pr-LEU2 cyh2 Δura3::UPRE-GFP-TEF2pr-RFP-MET15-**URA3::NAT-MM- MBP + pRS426-GPD-ΔN-VMS1-GFP** | **MM-DMMBP- pRS426-GPD-ΔN-VMS1-GFP** | PP402 | This study |
| **53** | **PP532** | *MATα* his3Δ1 leu2Δ0 met15Δ0 ura3Δ0 LYS+Δcan1::STE2pr-spHIS5 Δlyp1::STE3pr-LEU2 cyh2 Δura3::UPRE-GFP-TEF2pr-RFP-MET15-**URA3::NAT-MM-DMMBP+ pRS426-GPD-VMS1-Flag** | **MM-DMMBP- pRS426-GPD-VMS1-Flag_OE** | PP502 | This study |
| **54** | **PP542** | *MATα* his3Δ1 leu2Δ0 met15Δ0 ura3Δ0 LYS+Δcan1::STE2pr-spHIS5 Δlyp1::STE3pr-LEU2 cyh2 Δura3::UPRE-GFP-TEF2pr-RFP-MET15-**URA3::NAT-MM-DMMBP+ pJR8252A** | **MM-DMMBP- pRS416-VMS1-6HIS-2HA** | PP502 | This study |
| **55** | **PP552** | *MATα* his3Δ1 leu2Δ0 met15Δ0 ura3Δ0 LYS+Δcan1::STE2pr-spHIS5 Δlyp1::STE3pr-LEU2 cyh2 Δura3::UPRE-GFP-TEF2pr-RFP-MET15-**URA3::NAT-MM-DMMBP+ pJR3462D** | **MM-DMMBP- pRS416-VMS1-GFP** | PP502 | This study |
| **56** | **PP562** | *MATα* his3Δ1 leu2Δ0 met15Δ0 ura3Δ0 LYS+Δcan1::STE2pr-spHIS5 Δlyp1::STE3pr-LEU2 cyh2 Δura3::UPRE-GFP-TEF2pr-RFP-MET15-**URA3::NAT-MM-DMMBP+ pJR6107B** | **MM-DMMBP- pRS416-ΔN-VMS1-GFP** | PP502 | This study |
| **53** | **PP572** | *MATα* his3Δ1 leu2Δ0 met15Δ0 ura3Δ0 LYS+Δcan1::STE2pr-spHIS5 Δlyp1::STE3pr-LEU2 cyh2 Δura3::UPRE-GFP-TEF2pr-RFP-MET15-**URA3::NAT-MM-DMMBP+ pJR9532D** | **MM-DMMBP- pRS416-VMS1(1-417)** | PP502 | This study |
| **54** | **PP582** | *MATα* his3Δ1 leu2Δ0 met15Δ0 ura3Δ0 LYS+Δcan1::STE2pr-spHIS5 Δlyp1::STE3pr-LEU2 cyh2 Δura3::UPRE-GFP-TEF2pr-RFP-MET15-**URA3::NAT-MM-DMMBP+ pRS426-GPD-ΔN-VMS1-GFP** | **MM-DMMBP- pRS426-GPD-ΔN-VMS1-GFP** | PP502 | This study |
| **55** | **PP435** | *MATα* his3Δ1 leu2Δ0 met15Δ0 ura3Δ0 LYS+Δcan1::STE2pr-spHIS5 Δlyp1::STE3pr-LEU2 cyh2 Δura3::UPRE-GFP-TEF2pr-RFP-MET15-**URA3::NAT-IMS-MBP+ pRS426-GPD-VMS1-Flag** | **MM-DMMBP- pRS426-GPD-VMS1-Flag_OE** | PP405 | This study |
| **56** | **PP445** | *MATα* his3Δ1 leu2Δ0 met15Δ0 ura3Δ0 LYS+Δcan1::STE2pr-spHIS5 Δlyp1::STE3pr-LEU2 cyh2 Δura3::UPRE-GFP-TEF2pr-RFP-MET15-**URA3::NAT-IMS-MBP+ pJR8252A** | **MM-DMMBP- pRS416-VMS1-6HIS-2HA** | PP405 | This study |
| **57** | **PP455** | *MATα* his3Δ1 leu2Δ0 met15Δ0 ura3Δ0 LYS+Δcan1::STE2pr-spHIS5 Δlyp1::STE3pr-LEU2 cyh2 Δura3::UPRE-GFP-TEF2pr-RFP-MET15-**URA3::NAT-IMS-MBP+ pJR3462D** | **MM-DMMBP- pRS416-VMS1-GFP** | PP405 | This study |
| **58** | **PP465** | *MATα* his3Δ1 leu2Δ0 met15Δ0 ura3Δ0 LYS+Δcan1::STE2pr-spHIS5 Δlyp1::STE3pr-LEU2 cyh2 Δura3::UPRE-GFP-TEF2pr-RFP-MET15-**URA3::NAT-IMS-MBP+ pJR6107B** | **MM-DMMBP- pRS416-ΔN-VMS1-GFP** | PP405 | This study |
| **59** | **PP475** | *MATα* his3Δ1 leu2Δ0 met15Δ0 ura3Δ0 LYS+Δcan1::STE2pr-spHIS5 Δlyp1::STE3pr-LEU2 cyh2 Δura3::UPRE-GFP-TEF2pr-RFP-MET15-**URA3::NAT-IMS-MBP+ pJR9532D** | **MM-DMMBP- pRS416-VMS1(1-417)** | PP405 | This study |
| **60** | **PP485** | *MATα* his3Δ1 leu2Δ0 met15Δ0 ura3Δ0 LYS+Δcan1::STE2pr-spHIS5 Δlyp1::STE3pr-LEU2 cyh2 Δura3::UPRE-GFP-TEF2pr-RFP-MET15-**URA3::NAT-IMS- MBP + pRS426-GPD-ΔN-VMS1-GFP** | **MM-DMMBP- pRS426-GPD-ΔN-VMS1-GFP_OE** | PP405 | This study |
| **61** | **PP535** | *MATα* his3Δ1 leu2Δ0 met15Δ0 ura3Δ0 LYS+Δcan1::STE2pr-spHIS5 Δlyp1::STE3pr-LEU2 cyh2 Δura3::UPRE-GFP-TEF2pr-RFP-MET15-**URA3::NAT-IMS-DMMBP+ pRS426-GPD-VMS1-Flag** | **IMS-DMMBP- pRS426-GPD-VMS1-Flag_OE** | PP505 | This study |
| **62** | **PP545** | *MATα* his3Δ1 leu2Δ0 met15Δ0 ura3Δ0 LYS+Δcan1::STE2pr-spHIS5 Δlyp1::STE3pr-LEU2 cyh2 Δura3::UPRE-GFP-TEF2pr-RFP-MET15-**URA3::NAT-IMS-DMMBP+ pJR8252A** | **IMS-DMMBP- pRS416-VMS1-6HIS-2HA** | PP505 | This study |
| **63** | **PP555** | *MATα* his3Δ1 leu2Δ0 met15Δ0 ura3Δ0 LYS+Δcan1::STE2pr-spHIS5 Δlyp1::STE3pr-LEU2 cyh2 Δura3::UPRE-GFP-TEF2pr-RFP-MET15-**URA3::NAT-IMS-DMMBP+ pJR3462D** | **IMS-DMMBP- pRS416-VMS1-GFP** | PP505 | This study |
| **64** | **PP565** | *MATα* his3Δ1 leu2Δ0 met15Δ0 ura3Δ0 LYS+Δcan1::STE2pr-spHIS5 Δlyp1::STE3pr-LEU2 cyh2 Δura3::UPRE-GFP-TEF2pr-RFP-MET15-**URA3::NAT-IMS-DMMBP+ pJR6107B** | **IMS-DMMBP- pRS416-ΔN-VMS1-GFP** | PP505 | This study |
| **65** | **PP575** | *MATα* his3Δ1 leu2Δ0 met15Δ0 ura3Δ0 LYS+Δcan1::STE2pr-spHIS5 Δlyp1::STE3pr-LEU2 cyh2 Δura3::UPRE-GFP-TEF2pr-RFP-MET15-**URA3::NAT-IMS-DMMBP+ pJR9532D** | **IMS-DMMBP- pRS416-VMS1(1-417)** | PP505 | This study |
| **66** | **PP585** | *MATα* his3Δ1 leu2Δ0 met15Δ0 ura3Δ0 LYS+Δcan1::STE2pr-spHIS5 Δlyp1::STE3pr-LEU2 cyh2 Δura3::UPRE-GFP-TEF2pr-RFP-MET15-**URA3::NAT-IMS-DMMBP+ pRS426-GPD-ΔN-VMS1-GFP** | **IMS-DMMBP- pRS426-GPD-ΔN-VMS1-GFP_OE** | PP505 | This study |

**List of primers used for this study**

| **Primer Name** | **Sequence** |
| --- | --- |
| NAT2-URA3_F | 5'ATGTCGAAAGCTACATATAAGGAACGTGCTGCTACTCATCCTAGTC CGTACGCTGCAGGTCGACGG3' |
| CYC1-URA3_R | 5'GCTGGCCGCATCTTCTCAAATATGCTTCCCAGCCTGCTTTTC TGCTCACTATAGGGAGACCGGC3' |
| UraupF | 5'TATTCTTAACCCAACTGCACAG3' |
| UradownR | 5'GTTCAATGCGTCCATCTTTAC3' |
| NAT_Int_Rev_Primer | 5'CGAGATGACCACGAAGCCCGCC3' |
| MBP(WT)_FP_ApaINheI | 5'CCCGGGCCCGCTAGCATGAAAATCGAAGAAGGTAAACTG3' |
| MBP(WT)_RP_AvrII | 5'CCCCCTAGGTTACTTGGTGATACGAGTCTGCGCGTCTTTCAG3' |
| DHFR_ApaINheI_For | 5'CCCGGGCCCGCTAGCATGGTTCGACCATTGAACTGC3' |
| DHFR_AvrIIXbaI_Rev | 5'TCTAGACCTAGGTTATTAGTCTTTCTTCTCGTAGAC3' |
| PBMD_ApaI_F | 5'CCCGGGCCCATGCTTGGTGATACGAGTCTGCGCG3' |
| PBMD_AvrII_R | 5'CCCCCTAGGCGCGCAGACTCGTATCACCAAGCAT3' |
| UPRE-KanMX_F | 5'GGAACTGGACAGCGTGTCGAAAAAGCTCGACAGGAACTGGACAGCGTGGACATGGAGGCCCAGAATACCCTCC3' |
| KanMX-GFP_R | 5'TTATTTGTACAATTCATCCATACCATGGGTAATACCAGCAGCAGTAACCAGTATAGCGACCAGCATTCAC3' |
| PBMD-yeGFP_F | 5'GCAGATCTTTGTTATAAATCAGCGA ATGTCTAAAGGTGAAGAATTATTCAC3' |
| PBMD-yeGFP_R | 5'GTGAATAATTCTTCACCTTTAGACATTCGCTGATTTATAACAAAGATCTGC3' |
| yeGFP_AvrII_Rev | 5'CCCCCTAGG TTATTTGTACAATTCATCCATACC3' |
| PBMD_Int_For | 5'CCGTTGATGGTCATCGCTGTTTCGC3' |
| PBMD_Int_Rev | 5'GGTTAATAAAGACAAACCGCTGGG3' |
| PBMD_BamHI_F | 5'CCCGGATCCTATGCTTGGTGATACGAGTCTGCGC3' |
| PBMD_SalI_R | 5'CCCGTCGACTTATCGCTGATTTATAACAAAGAT3' |
| PBMD_2xFLAG_AvrII_R | 5'GGGGTCGACCCTAGGTTA CTTATCGTCGTCATCCTTGTAATC CTTATCGTCGTCATCCTTGTAATC TCGCTGATTTATAACAAAGATCTGCTG3' |
| VMS1_SpeI-BamHI_F | 5'GGGACTAGTGGATCCATGAATTCACAAAAGGCGAG3' |
| VMS1_ClaI-SalI_R | 5'GGGGTCGACATCGATTCAGTATTTCTTTTTCATCC3' |
| VMS1-KanMX_F | 5'GGATTTTCAAAAGATCTGCACGCCTGTTGACAAGCTTCCAATAGC CTGAAGCTTCGTACGCTGCAGGTCG3' |
| VMS1-KanMX_R | 5'TGCAAATGCTAAGAAAAATCCTAAAAATTTGAATATGAGATATTC GGCCACTAGTGGATCTGATATCACC3' |
| VMS1_ClaI-SalI_2xFlag_R | 5'GGGGTCGACATCGATTTACTTATCGTCGTCATCCTTGTAATCGCCCTTATCGTCGTCATCCTTGTAATCGTATTTCTTTTTCATCCTTTC3' |
| GFP_ClaI-SalI_R | 5'GGGGTCGACATCGATTTATTTGTATAGTTCATCCATGCC3' |
| VMS1_Int_F | 5'CTGCTGAAGAAAGGAAGGGCACCG3' |
| VMS1_Int_R | 5'GTGCGCGTTCCCCTTGACATTGAGTC3' |

List of antibodies used for this study

| **Antibody** | **Source** |
| --- | --- |
| Anti-MBP Monoclonal Antibody | E8032L(NEB) |
| Porin Monoclonal Antibody (16G9E6BC4) | 459500(Thermo Fisher) |
| DHFR Antibody | #45710(CST) |
| DYKDDDDK Tag Antibody | 2044S(CST) |
| PGK1 Antibody | #68540(CST) |
| Anti-GFP antibody | Ab290(abcam) |
| yHSP60 anti-sera | In-house generated |
| SSC1 anti-sera | In-house generated |
| TOM70 anti-sera | In-house generated |
