## Supplementary figures and images for "A comparative study of stress responses elicited by misfolded proteins targeted by bipartite or matrix-targeting signal sequences to yeast mitochondria"

### Supplementary Figure 1

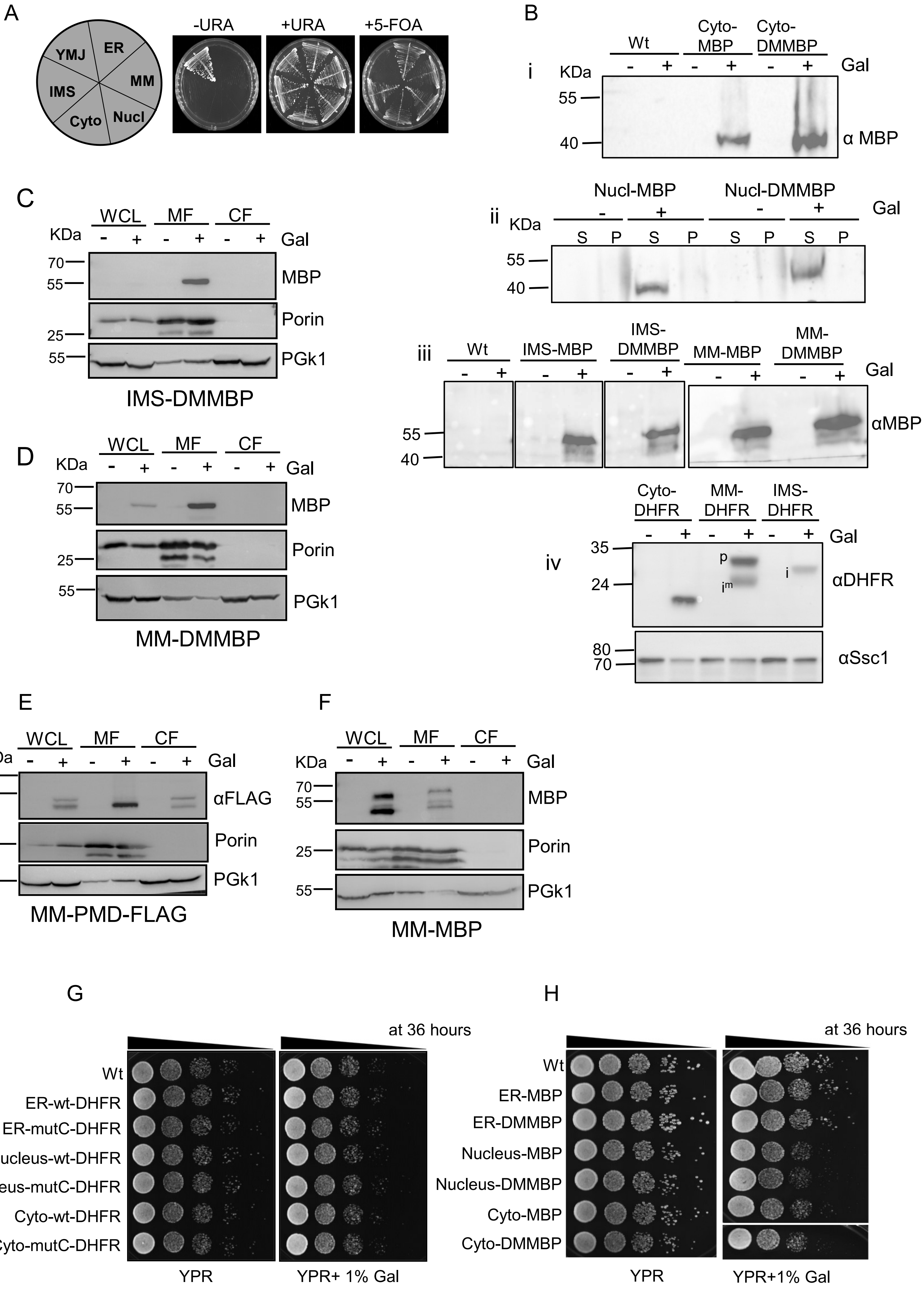

### Supplementary Figure 2

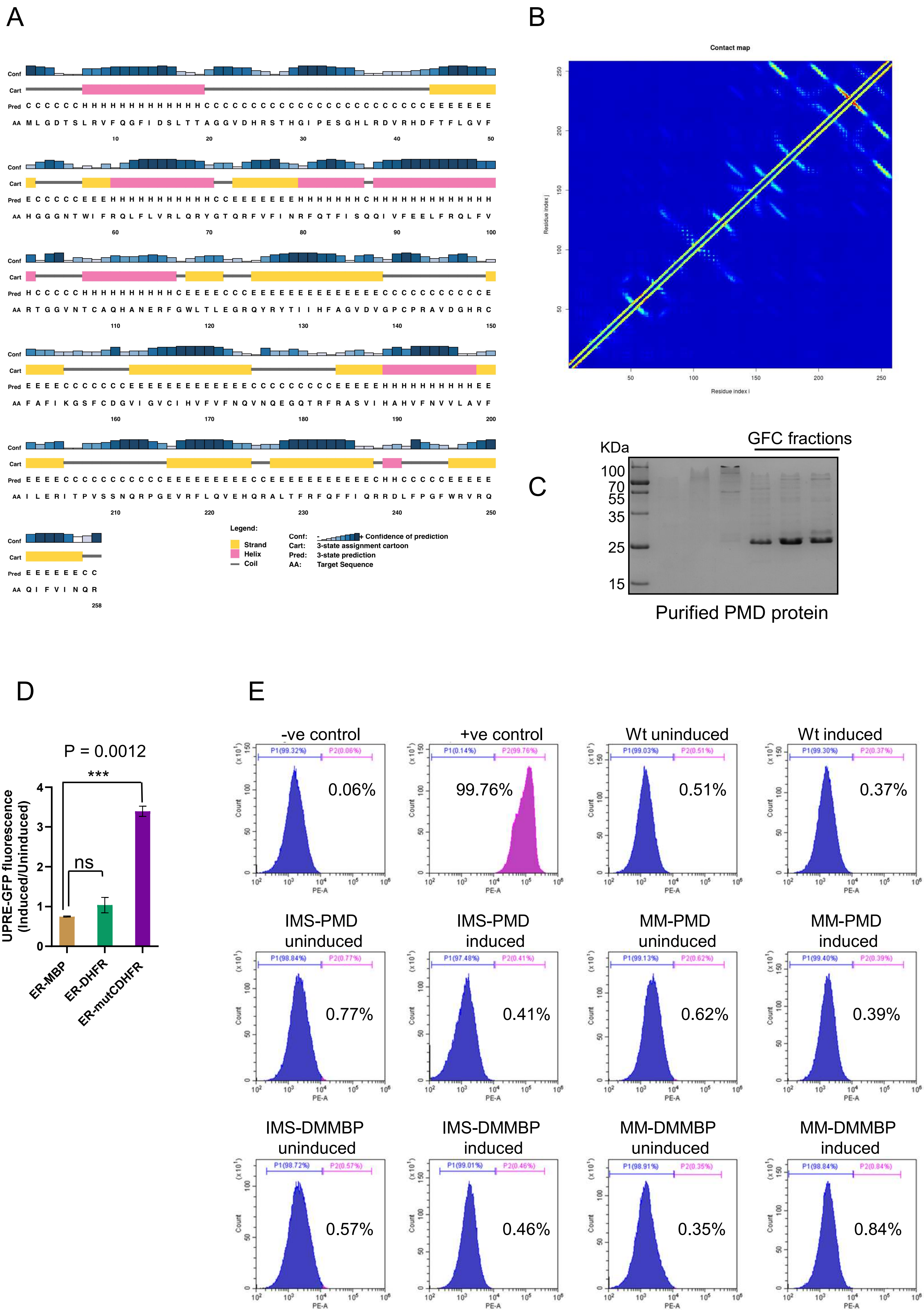

### Supplementary Figure 3

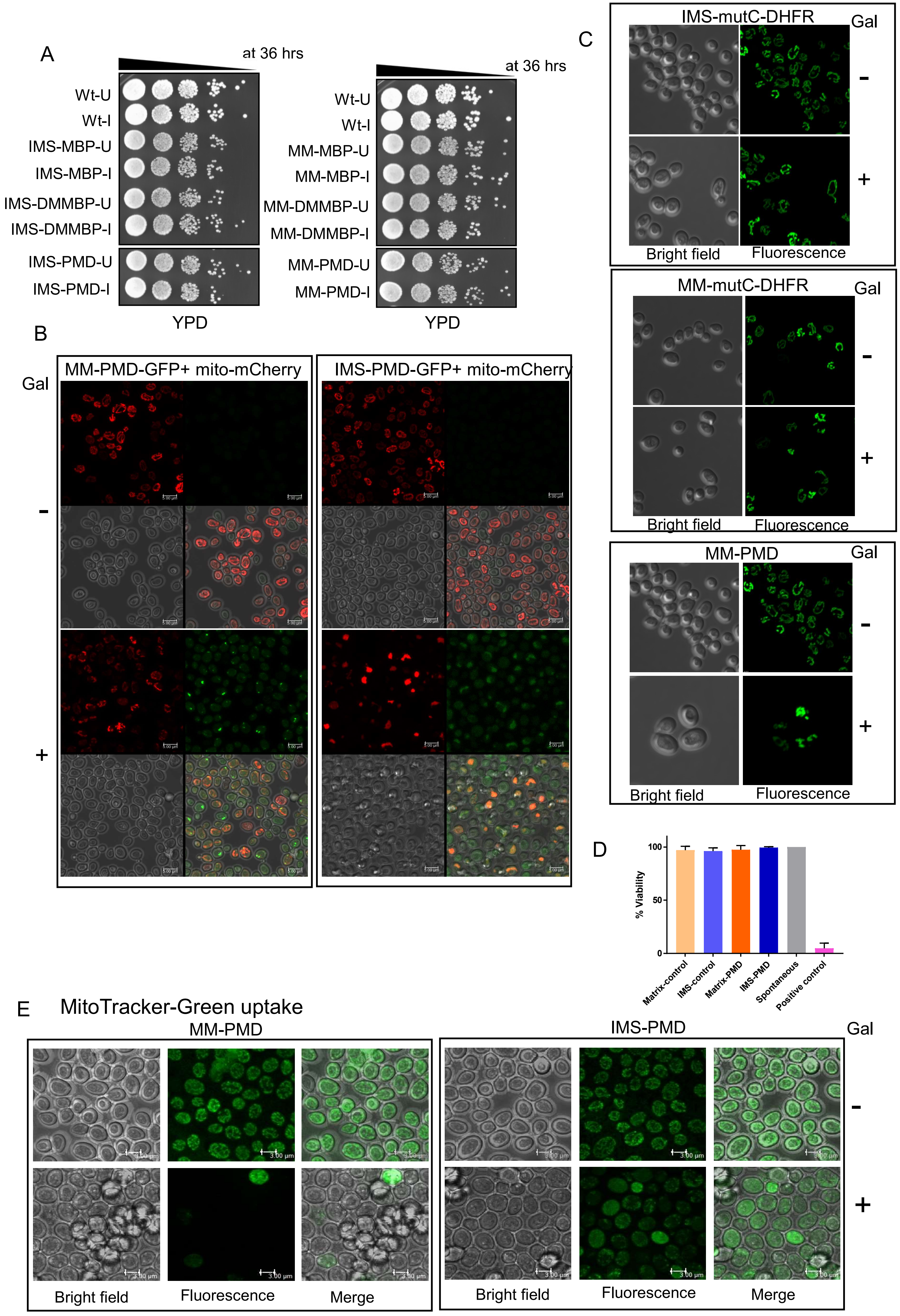

### Supplementary Figure 4

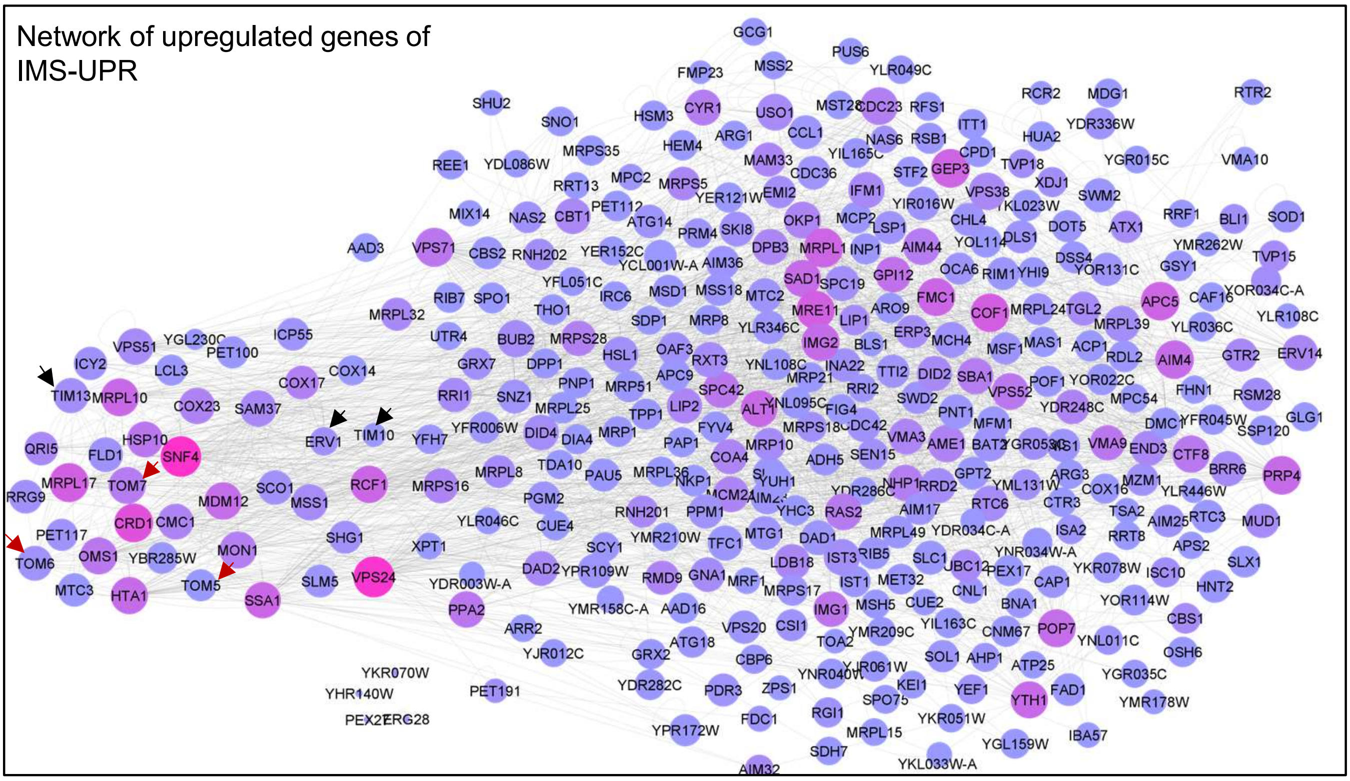

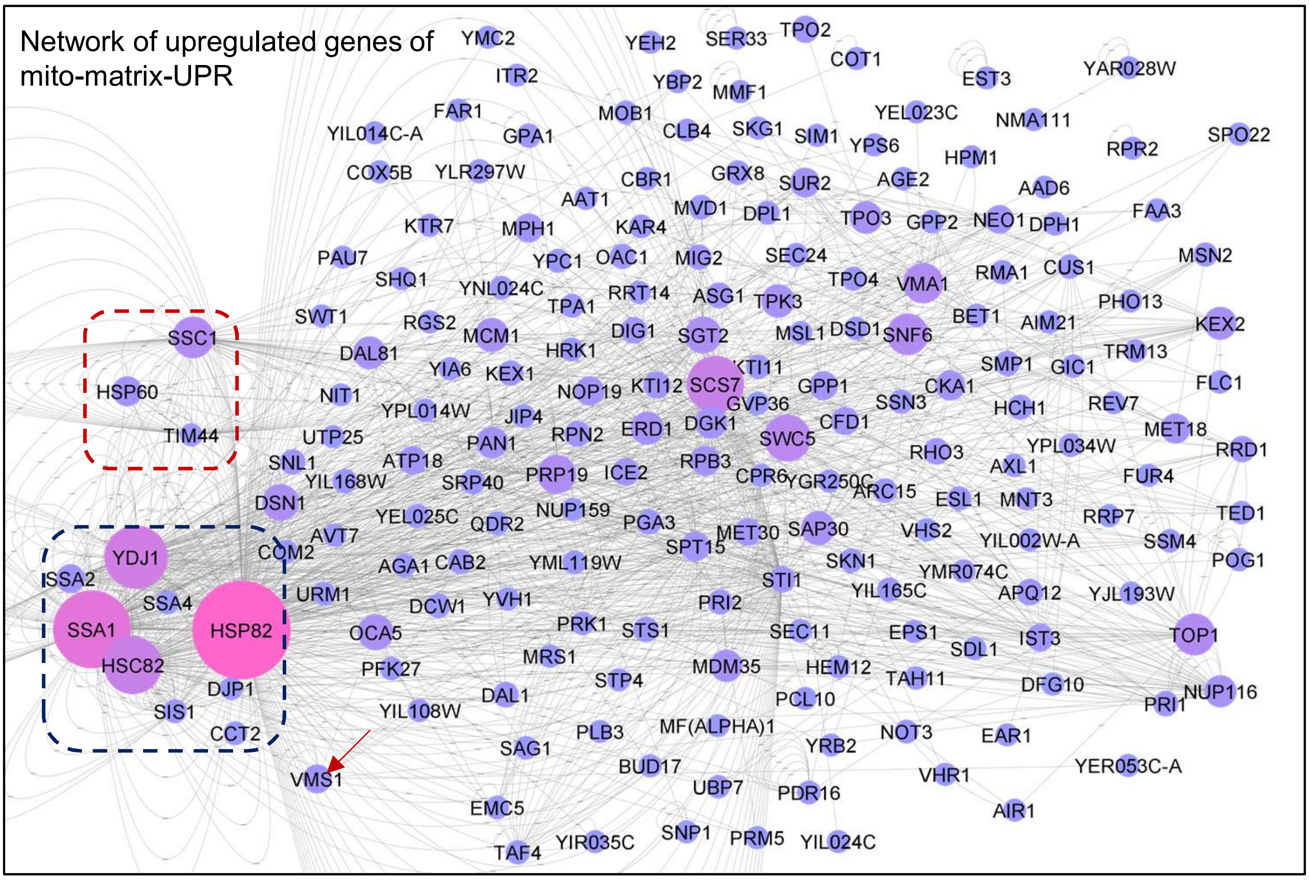

### Supplementary Figure 5

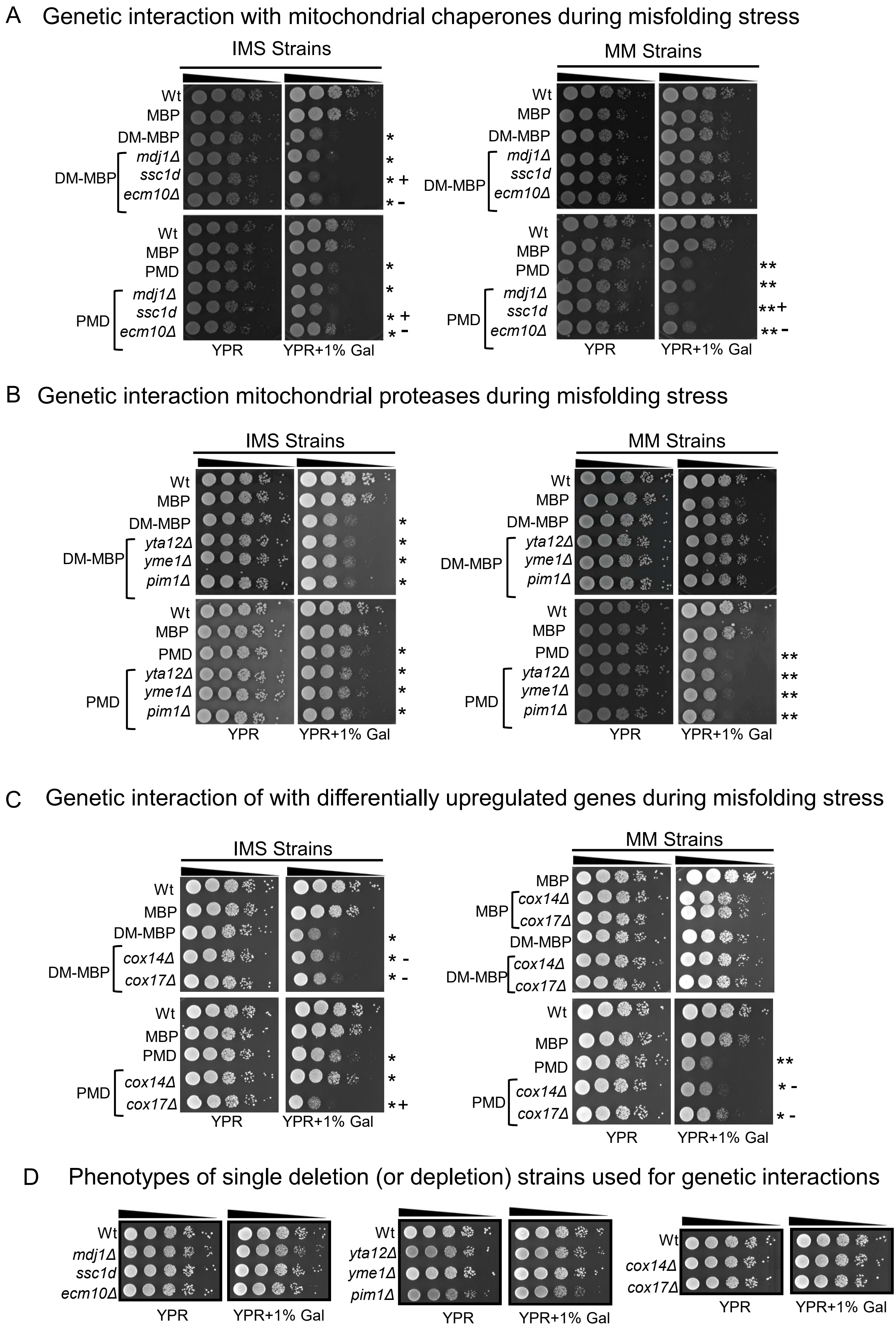

### Supplementary Figure 6

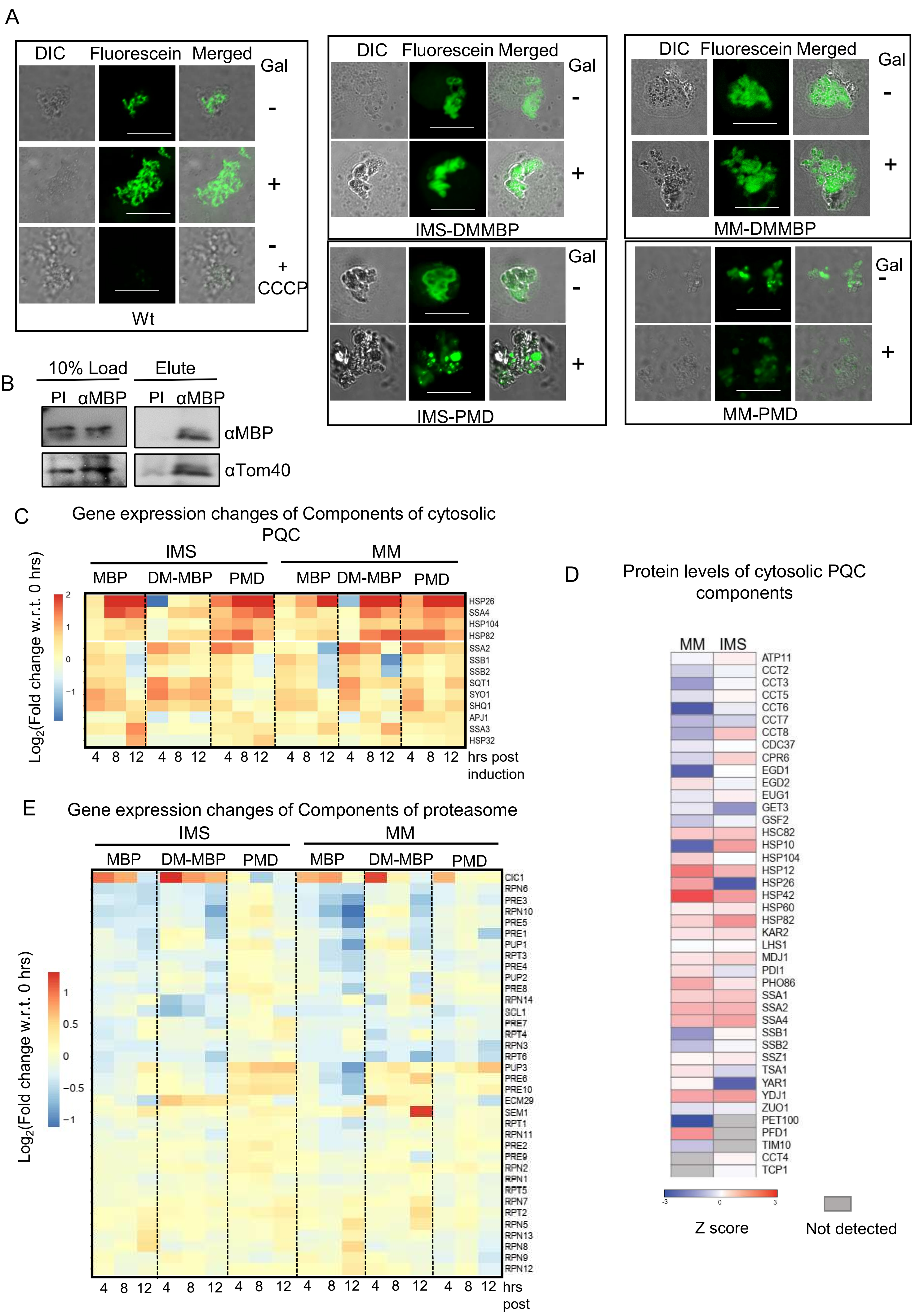

### Supplementary Figure 7

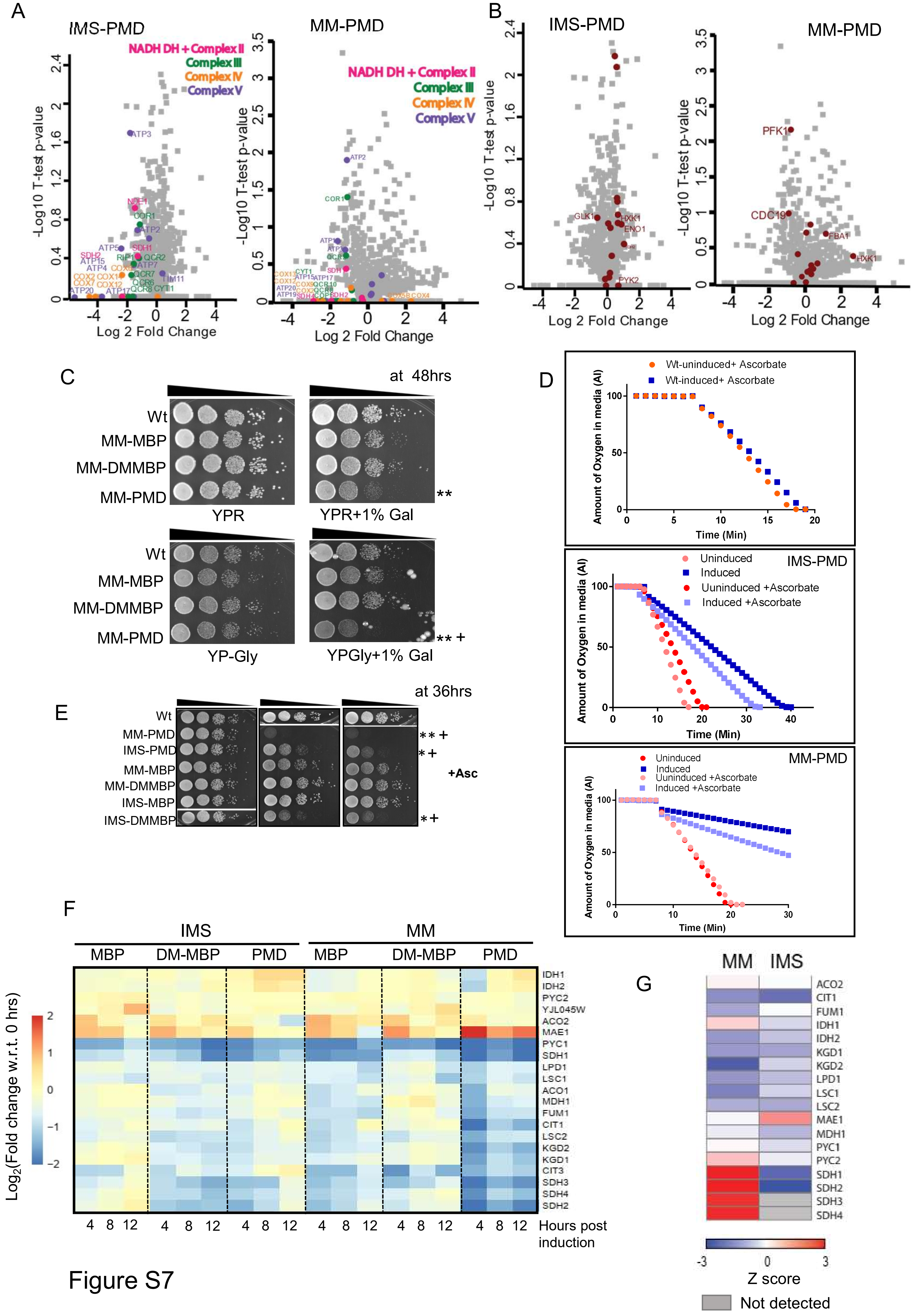

### Supplementary Figure 8

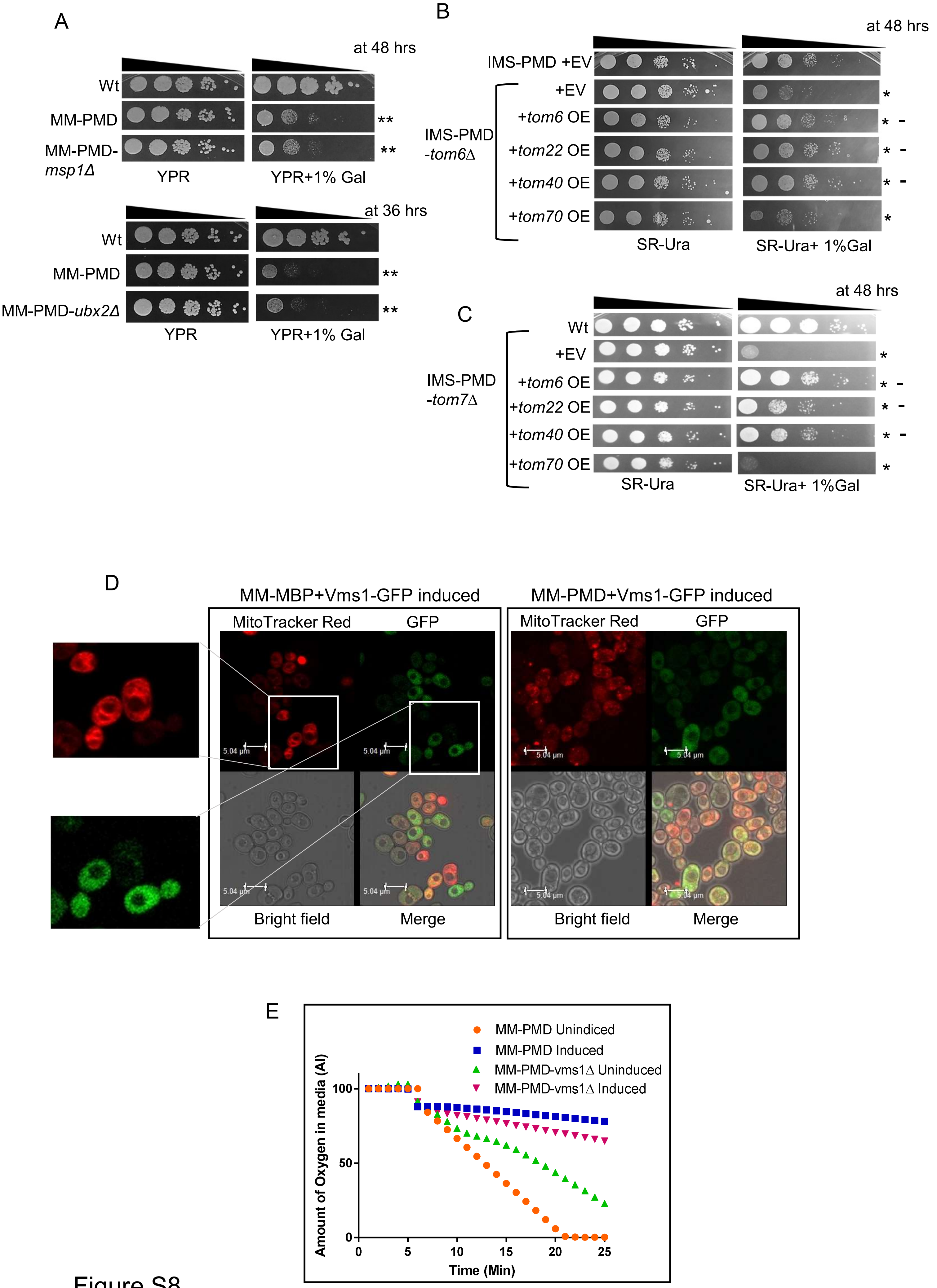
